# Striatal Dopaminergic Dysfunction Constrains Motor Invigoration in Parkinson’s Disease

**DOI:** 10.64898/2026.09.02.748476

**Authors:** Birgitte Liang Chen Thomsen, Vytautas Labanauskas, Nora Raaf, Sune Gram Thomsen, Mikkel Christoffer Vinding, Lisbeth Marner, Annemette Løkkegaard, David Meder, Hartwig Roman Siebner

## Abstract

Bradykinesia is a cardinal motor feature of Parkinson’s disease and is characterised by a reduced ability to generate movements with normal speed and force. Although nigrostriatal dopamine loss is regarded as the primary cause, direct human evidence linking dopaminergic degeneration to striatal dysfunction and impaired motor output and vigour remains limited.

To investigate this relationship, we developed a grip-force paradigm that quantified movement vigour as the ability to rapidly generate force. Fifty-three individuals with Parkinson’s disease, 30 with levodopa-induced dyskinesia and 23 without, and 25 age-matched healthy controls performed the task during 3T functional MRI. Thirty-five patients additionally underwent dopamine transporter PET imaging to assess associations among nigrostriatal dopaminergic integrity, striatal activity and motor performance.

Patients with Parkinson’s disease showed marked reductions in initial movement vigour accompanied by reduced bilateral putaminal activation during force generation. Lower putaminal dopamine transporter binding was associated with both poorer movement vigour and weaker grip-related putaminal activation during functional MRI, establishing a direct link between nigrostriatal degeneration, impaired striatal recruitment and motor slowing. Functional MRI further revealed reduced reward-related responses in the nucleus accumbens. Across participants, stronger reward-related ventral striatal activity was associated with greater movement vigour, suggesting that motivational processes contribute independently to motor performance. Patients with levodopa-induced dyskinesia exhibited more severe putaminal dopaminergic denervation than patients without dyskinesia but showed no additional impairment of movement vigour or striatal task responses.

These findings provide direct *in vivo* evidence that dorsal nigrostriatal dopaminergic degeneration constrains movement vigour in Parkinson’s disease through reduced striatal recruitment during action generation. Impaired reward-related signalling in ventral striatum emerges as an additional, partly independent mechanism influencing motor performance, highlighting distinct motor and motivational contributions to bradykinesia.

## Introduction

Parkinson’s disease (PD) leads to progressive degeneration of dopaminergic neurons in the substantia nigra pars compacta and dopamine depletion within the striatum.^1,2^ Loss of nigrostriatal dopamine disrupts cortico-basal ganglia-cortical circuits, resulting in the cardinal motor syndrome of bradykinesia.^3–5^ While clinically defined by reduced speed and amplitude of movement, bradykinesia comprises several physiological deficits that likely arise from distinct underlying neural mechanisms.^6^

A central component of bradykinesia is impaired motor invigoration, defined as the reduced ability to rapidly mobilise motor output during voluntary action.^7–11^ Motor invigoration is expressed behaviourally as movement vigour and can be quantified using the rate at which force is generated during the earliest phase of movement.^12–17^ In PD, insufficient recruitment of agonist musculature results in segmented force production during ballistic motor tasks and generalises across effectors and behavioural contexts.^15,17^ Such observations suggest that impaired motor invigoration represents a fundamental pathophysiological feature of PD rather than a task-specific motor deficit.

Substantial theoretical, computational, and experimental evidence implicates dopamine in the regulation of motor invigoration.^18,19^ Dopaminergic projections within dorsal striatal motor circuits are thought to facilitate the scaling of motor output, whereas mesolimbic dopaminergic projections to the ventral striatum contribute to the valuation of actions and anticipated outcomes.^20–25^ Consistent with this framework, movement vigour scales with the expected value of the action’s outcome.^18^ These observations have led to the influential hypothesis that dopamine promotes vigorous action both by enabling motor execution and by energising behaviour through value-based motivational mechanisms.^25–28^

A central mechanistic model of PD therefore posits that nigrostriatal degeneration reduces striatal dopaminergic tone, diminishes striatal functional engagement, and consequently impairs motor invigoration.^25,29,30^ Although supported by extensive preclinical research and indirect evidence from human studies, direct *in vivo* demonstrations linking dopaminergic denervation, striatal functional responses, and quantitative measures of movement invigoration within the same individuals remain limited.^24,31–38^ Consequently, it remains unclear whether impaired movement vigour in PD represents a direct consequence of dorsal striatal dopamine loss and reduced striatal engagement during action.

To address this gap, we combined a novel multidomain grip-force paradigm with task-based functional MRI (fMRI) and dopamine transporter (DAT) PET imaging. Patients with PD in a pragmatic OFF-medication state and age-matched healthy controls performed visually cued unilateral isometric hand grip responses. We focused on the first peak in the rate of force development, a sensitive physiological measure of early motor invigoration reflecting the strength of initial agonist motor drive (i.e., initial vigour). In a subgroup of patients, DAT PET was used to quantify nigrostriatal dopaminergic integrity and assess its relationship with task-evoked striatal activity and motor performance.

We hypothesised that PD would be associated with impaired initial vigour and reduced recruitment in the motor territory of the striatum.^4^ Furthermore, we predicted that lower putaminal DAT binding would be associated with attenuated movement-related putaminal activation and diminished motor invigoration, providing direct *in vivo* evidence for a mechanistic link between dorsal striatal dopamine loss, striatal dysfunction, and bradykinesia.

Because motor invigoration is closely linked to motivational processes, we additionally examined reward-related modulation of behaviour and neural activity. The task incorporated contexts with differing reward contingencies, allowing us to determine how reward expectation influences motor invigoration and striatal responses. We hypothesised that PD would be associated with attenuated reward-related signalling within ventral striatal circuits, particularly the nucleus accumbens,^39^ and that reduced reward sensitivity would contribute to impaired motor invigoration independently of deficits in dorsal striatal motor circuits.

Finally, because levodopa-induced dyskinesia (LID) has been linked to maladaptive plasticity within cortico-basal ganglia networks,^40–42^ we explored whether relationships among dopaminergic denervation, striatal activation, reward processing, and motor invigoration differed between patients with and without LID.

## Methods and Materials

### Participants

We included 38 patients with akinetic-rigid type Parkinson’s disease with LID (PD-LID), 27 patients without LID (PD-NoLID), and 26 age-matched healthy controls (HC) for the MRI acquisition.

Patients with PD fulfilled the Movement Disorder Society (MDS) diagnostic criteria.^43^ Exclusion criteria included (i) other neurological or psychiatric disorders, (ii) advanced treatments (e.g., deep brain stimulation or continuous dopaminergic infusion therapies), (iii) changes in antiparkinsonian medication within four weeks, (iv) use of antipsychotics, donepezil, orphenadrine, or GABAergic medication, (v) frequent use of benzodiazepines or opioids (>1/week), (vi) contraindications to MRI, or (vii) severe tremor.

A total of 14 participants dropped out or were excluded from analysis. Reasons for exclusion included withdrawal between study sessions (n = 6), claustrophobia (n = 3), severe symptom exacerbation following medication withdrawal (n = 1), inability to fit within the scanner (n = 1), and misunderstanding of the task (n = 1). The behavioural analysis included 30 patients with PD-LID (15 females), 23 with PD-NoLID (10 females), and 25 HC (13 females).

Two participants, one with and one without LID, were excluded from the imaging analysis because fewer than 10 trials exceeded the response threshold for either right-go or left-go trials. Furthermore, 16 patients with PD-LID and 19 with PD-NoLID participated in DAT PET imaging. One PD-NoLID participant was excluded from the analysis using MRI-defined ROIs due to misalignment of the regions.

All participants provided written informed consent in accordance with the Declaration of Helsinki.^44^ The study was approved by the Regional Research Ethics Committee of the Capital Region of Denmark (H-22010296) and registered on clinicaltrials.gov (NCT06240624).

### Experimental Procedures

Patients with PD participated in up to three test days. On day 1, we performed Parkinson-specific clinical assessments (including the MDS Unified Parkinson’s Disease Rating Scale (MDS-UPDRS),^45^ Unified Dyskinesia Rating Scale (UDysRS)^46^ as well as neuropsychological assessments (Table 1). HC performed the neuropsychological assessments on the day of the MRI scan.

**Table 1.** Demographic and Clinical Characteristics. Values are presented as mean ± SD unless otherwise specified. Group comparisons were performed using one-way ANOVA or independent-samples t-tests for normally distributed variables, and Kruskal– Wallis or Mann–Whitney U tests otherwise. Categorical variables were compared using chi-square or Fisher’s exact tests. Post hoc comparisons were adjusted for multiple testing. Different superscript letters indicate statistically significant differences between groups following post hoc analyses, whereas groups sharing the same superscript letter do not differ significantly. PD: Parkinson’s disease; LID: levodopa-induced dyskinesia. LEDD: Levodopa equivalent daily dose; MDS-UPDRS: Movement Disorder Society Unified Parkinson’s Disease Rating Scale; UDysRS: Unified Dyskinesia Rating Scale, NMSS: Non-Motor Symptoms Scale for Parkinson’s Disease; MoCA: Montreal Cognitive Assessment;^54^ MDI: Major Depression Inventory;^55^ BIS-11: Barratt Impulsiveness Scale;^56^ LARS: Lille Apathy Rating Scale;^57^ QUIP: Parkinson’s Disease Impulsive-Compulsive Disorders Questionnaire.^58^

|  | Healthy Controls | Patients PD without LID | Patients with PD and LID | P-value |
| --- | --- | --- | --- | --- |
| N | 25 | 23 | 30 |  |
| Sex (Female/Male) | 13/12 | 10/13 | 15/15 | 0.83 |
| Handedness (R/L/A) | 22/2/1 | 18/1/4 | 26/2/2 | 0.59 |
| Age [years] | 66.0 ±9.08 | 66.2 ±9.11 | 61.7 ±9.05 | 0.12 |
| Education [years] | 14.2 ±2.85 | 14.3 ±2.91 | 14.5 ±3.36 | 0.72 |
| Disease duration [years] | NA | 6.45 ±2.89 | 7.91 ±3.74 | 0.16 |
| LEDD [mg] | NA | 639 ±249 | 806 ±401 | 0.10 |
| MDS-UPDRS Total (ON) | NA | 46.7 ±18.5<sup>a</sup> | 57.3 ±21.6<sup>b</sup> | 0.048 |
| MDS-UPDRS I | NA | 7.87 ±5.57 | 9.87 ±5.22 | 0.091 |
| MDS-UPDRS II | NA | 12.1 ±6.54 | 13.5 ±5.78 | 0.47 |
| MDS-UPDRS III (OFF) | NA | 33.9 ±11.1 | 39.5 ±12.8 | 0.15 |
| MDS-UPDRS III (ON) | NA | 21.2±9.6 | 23.3 ±12.2 | 0.61 |
| MDS-UPDRS IV | NA | 2.17 ±2.31<sup>a</sup> | 6.20 ±2.67<sup>b</sup> | <0.0001 |
| UDysRS | NA | NA | 27.0 ±9.12 | NA |
| NMSS | NA | 45.1 ±32.4 | 51.8 ±31.3 | 0.33 |
| MOCA | 27.8 ±1.71 | 27.5 ±2.29 | 28.0 ±1.77 | 0.80 |
| MDI | 4.64 ±4.55<sup>a</sup> | 7.95 ±5.05<sup>ab</sup> | 8.10 ±6.09<sup>b</sup> | 0.022 |
| BIS-11 | 45.1 ±7.40 | 43.6 ±9.01 | 44.3 ±9.07 | 0.84 |
| LARS | -29.4 ±3.61 | -28.2 ±4.49 | -28.5 ±3.43 | 0.47 |
| QUIP | NA | 0.652 ±1.97 | 1.20 ±1.86 | 0.091 |

On day 2 (day 1 for HC), patients with PD arrived in a pragmatic OFF-medication state (≥12 h withdrawal, for details see supplementary material). Motor symptom severity (MDS-UPDRS III) was assessed before the MRI scan. Participants were then trained on the task and performed maximal voluntary contraction (MVC) measurements. Structural MRI was acquired, followed by 10 minutes of the fMRI task in the OFF-medication state. Subsequently, patients with PD were administered dopaminergic medication and task-based fMRI resumed shortly after. Data acquired after levodopa administration will be reported separately. On day 3, a subgroup of the participants with PD underwent DAT PET.

### Experimental Task

Participants performed a go/no-go task presented using PsychoPy (v2021.2.0) incorporating emotional and reward components (Fig. 1A). Participants were instructed to respond with rapid grip-force contractions using the right or left hand (go) or to withhold responses (no-go). Grip force was recorded continuously from both hands using MRI-safe transducers (Current Designs Inc.) with a response threshold of approximately 70 N.

**Figure 1.**
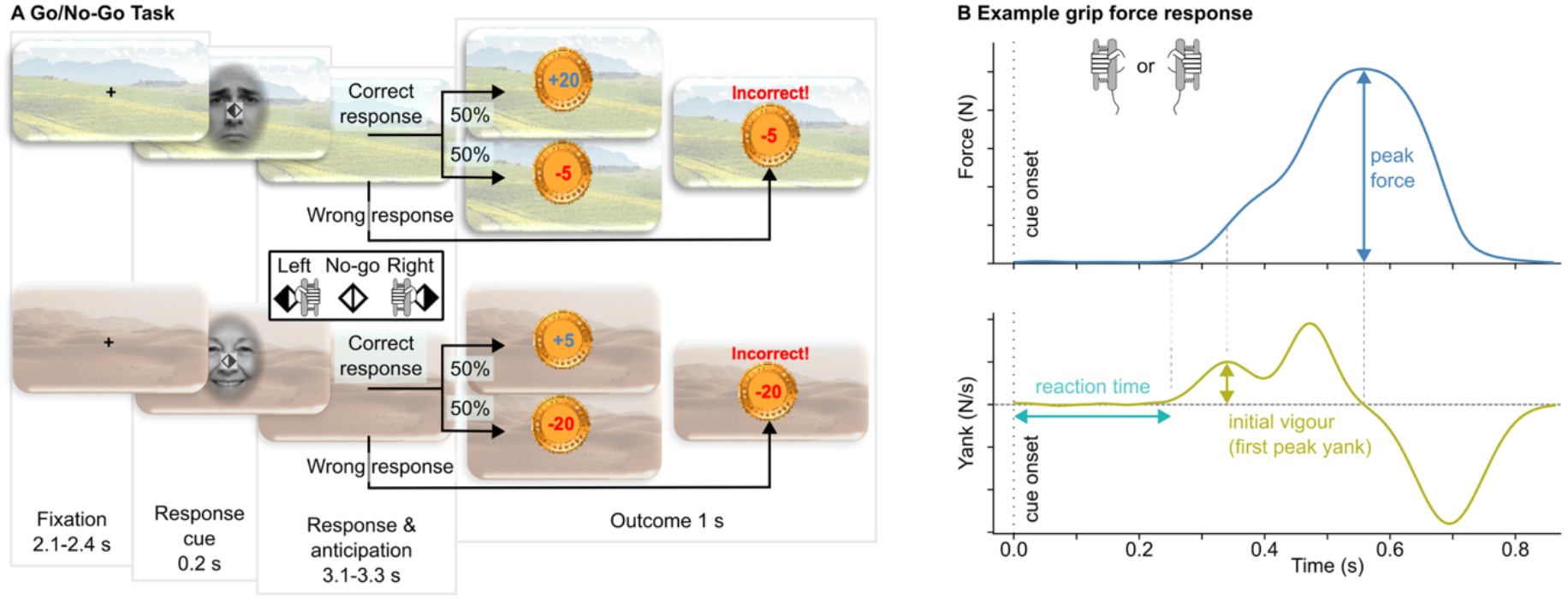
Go/No-Go task. A. Participants performed a visually cued grip-force task, responding by squeezing with either the left or right hand (go trials) or withholding a response (no-go trials) depending on the presented symbol. Correct responses were followed by a 50% probability of either winning or losing money, whereas incorrect responses always resulted in a monetary loss. The background (meadow or desert) indicated the motivational context. The meadow was associated with high reward and low loss, while the desert was associated with low reward and high loss. Response cues were presented in front of faces expressing different emotions (happy, neutral, or sad). B. Example of a grip response with force (top) and yank (first derivative N/s, bottom) and the three main outcome variables: peak force, reaction time and initial vigour. Due to copyright issues, the emotional faces in this figure are not from the FACES database^47^ but were created with AI (Microsoft Copilot based on GPT-5).

For each trial, a symbol indicating the required response was shown overlaid on a face with varying emotional expressions (happy, sad, or neutral)^47^ for 200 ms. The emotional cues were added because the task was also used in a depression trial; effects of emotional expression were not analysed further in this study. Correct responses resulted in a monetary reward or a loss (50% probability). Wrong responses always resulted in a loss, and the text “Forkert!” (Danish for “Incorrect!”) was displayed. The magnitude of rewards and losses depended on the motivational context, conveyed by the background: trials with a meadow in the background indicated a high-reward context with +20 or −5 Danish kroner (DKK) outcomes, while a desert signalled a high-loss context with +5 or −20 DKK outcomes. Trials were grouped into short blocks of four trials of the same motivational context with a maximum of two consecutive blocks of the same context. The task comprised 72 trials (24 of each response type). For additional details see supplementary material.

### MRI Acquisition

We used a 3T Siemens MAGNETOM Prisma scanner to acquire a T1-weighted MPRAGE (voxel size = 1 mm^3^; slices = 208; TR = 2700 ms; TE = 3.61 ms; flip angle = 9°) and functional MRI using a T2*-weighted echo-planar imaging (EPI) sequence (voxel size = 2.5 mm^3^; slices = 62; TR = 1800 ms; TE = 30 ms; flip angle = 65°). Pulse and respiratory cycles were recorded during MRI acquisition.

### DAT PET Acquisition

DAT PET with [^18^F]FE-PE2I assessed presynaptic dopaminergic integrity in the striatum. PET scans were acquired on a Discovery 710 or MI PET/CT (GE Healthcare). Imaging was performed 30 minutes after tracer injection, with 10 minutes of acquisition.^48^

### Behavioural Measures

Grip force data were low-pass filtered at 15 Hz and resampled to 120 Hz. The rate of force development (yank) was calculated as the first temporal derivative (df/dt) (Fig. 1B). Trials were classified as correct and included in the behavioural analyses if force exceeded 5% of MVC. This post hoc criterion was chosen to account for inter-individual differences in the maximum grip strength. Reaction time (ms), peak force (N), and initial vigour (N/s) were extracted from each trial. Reaction time was defined as the time at which the yank exceeded the rectified baseline (1 s before cue-onset) + 2.5 standard deviations and log-transformed for statistical testing. Initial vigour was quantified as the first peak in the df/dt output occurring between 25 ms after the reaction time and the time of peak force. Unusually fast reaction times (< 250 ms) and low initial vigour (< 0.25 AU) values were visually inspected and corrected as necessary. For details see supplementary material.

Omission errors were defined as go trials not reaching 5% MVC. Commission errors were responses with the non-prompted hand occurring before the response with the prompted hand (go trials), or any response (no-go trials). Responses below 5% of the individual MVC were manually inspected to identify commission errors.

### fMRI Preprocessing

Functional MRI data were preprocessed using fMRIPrep (v25.2.4) with a study-specific template generated by averaging participants’ T1 images (supplementary material).

### fMRI Statistical Analysis

Only trials that met the response threshold criteria (no “Incorrect” feedback during the task) and were not classified as commission errors post hoc were included in the fMRI analysis. Regressors of interest were modelled in SPM12 for right- and left-go events (response onset, duration 0.0 s) together with their parametric modulation with trial-wise initial vigour, nogo events (cue onset, duration 0.2 s), reward and loss events (duration 1.0 s), all separately for each motivational context. We defined contrasts for the three response types, the effect of motivational context on go responses (high-reward > high-loss context across right- and left-go trials), the vigour parametric modulator, and the linear scaling with outcome magnitude (high loss < low loss < low win < high win). Second-level group analyses were performed using one-way ANOVA models adjusted for age and sex. Statistical significance was defined using a voxel-wise entry threshold of *P* < 0.001 (uncorrected), with a minimum cluster extent of 30 voxels, and cluster-level family-wise error correction at *P* < 0.05.

Region-of-interest (ROI) analyses used activation likelihood estimation clusters reported in a meta-analysis,^49^ as well as spherical ROIs (5 mm radius) centred on peak coordinates in the supplementary motor area (SMA)/preSMA and right inferior frontal gyrus (rIFG) reported in another PD-LID study.^50^ Furthermore, based on previously reported blunted reward responses,^39^ we included the nucleus accumbens, ventral tegmental area (VTA), hippocampus, and amygdala from the WFU PickAtlas toolbox (version 3.0.5).^51^ At the group level, mean initial vigour (averaged across trials for the contralateral hand) was included as a covariate in second-level ANOVA models.

### DAT PET Analysis

DAT availability was quantified using the dopamine transporter-specific binding ratio (SBR) as previously described.^48^ The mean dopamine SBR was calculated from the whole putamen. When relating dopamine SBR to fMRI activation in putamen and nucleus accumbens, SBR was extracted from the fMRI ROIs co-registered to the DAT PET scans and median SBR values were used to reduce segmentation uncertainty near ventricles.

### Statistical Analysis

Statistical analyses were performed in R (v4.5.2).

### Demographic and Clinical Variables

Group differences in continuous variables were evaluated using one-way ANOVA or independent-samples t-tests (parametric) and Kruskal-Wallis test or Mann-Whitney U (non-parametric). Categorical variables were analysed using chi-square tests or Fisher’s exact test. Post hoc analyses were conducted using post hoc Tukey-adjusted comparisons or Dunn’s test with Bonferroni correction.

### Task performance

Behavioural outcomes (reaction time, peak force, and initial vigour) were analysed using linear mixed-effects models. All models included group, age and sex as fixed effects and participant as a random intercept. To handle disease lateralisation (not applicable to HC), two separate models compared each side in PD (most/least affected) against HC. A third model tested lateralisation within the PD groups, including side as a within-subject factor with participant-specific random slope. Statistical significance of fixed effects was assessed using Type III F-tests. Pairwise comparisons of estimated marginal means with Tukey adjustment followed significant effects of group.

Effects of motivational context were assessed using participant-level differences between high-reward and high-loss conditions (high reward – high loss), analysed using linear models adjusted for group, age, and sex. Overall effects were tested by comparing adjusted estimated marginal mean differences against zero using t-tests, and group differences using Type III F-tests.

Error counts were analysed using negative binomial regression models including group, age, and sex as covariates. Tukey-adjusted post hoc comparisons followed significant group effects.

### Functional MRI data

Imaging measures extracted from ROIs with repeated observations (most and least affected sides) were analysed using linear mixed-effects models including side as a within-subject factor. Associations between motor vigour and BOLD activity were analysed using a model that included group, side, and their interaction. Associations between dopamine SBR and BOLD responses were analysed using a similar model including SBR, group, side, and their interactions. The linear mixed-effect models included a random intercept for participant.

Associations between behavioural or clinical measures and nucleus accumbens activity were analysed using linear regression models with activity (averaged across hemispheres), group, and their interaction as predictors. All models included sex and age as covariates unless otherwise specified.

Statistical significance of fixed effects was assessed as described for the behavioural linear mixed-effects models.

### Model Selection

Exploratory model selection was performed using MuMIn (dredge)^52^, ranked by corrected Akaike Information Criterion (AICc). Predictor importance was assessed using summed Akaike weights (Σwi).^53^

## Results

### Participant Characteristics

Groups were comparable in age, sex, and most clinical variables. However, MDI scores were higher in PD-LID compared to HC, but not significantly different between PD groups. PD groups only differed in total ON-medication MDS-UPDRS and motor complication burden (MDS-UPDRS IV) with PD-LID having higher scores than PD-NoLID (Table 1).

### Motor Performance

To determine whether task performance differed across groups, we compared reaction time, peak force, and initial vigour. Reaction time did not show a significant effect of group or side (Fig. 2A). Mean (± SD) reaction time was 602 ± 60 ms in HC, 577 ±67 ms and 594 ±80 ms in PD-NoLID (most/least affected side) and 611 ±109 ms and 603 ±117 ms in PD-LID (most/least affected).

**Figure 2.**
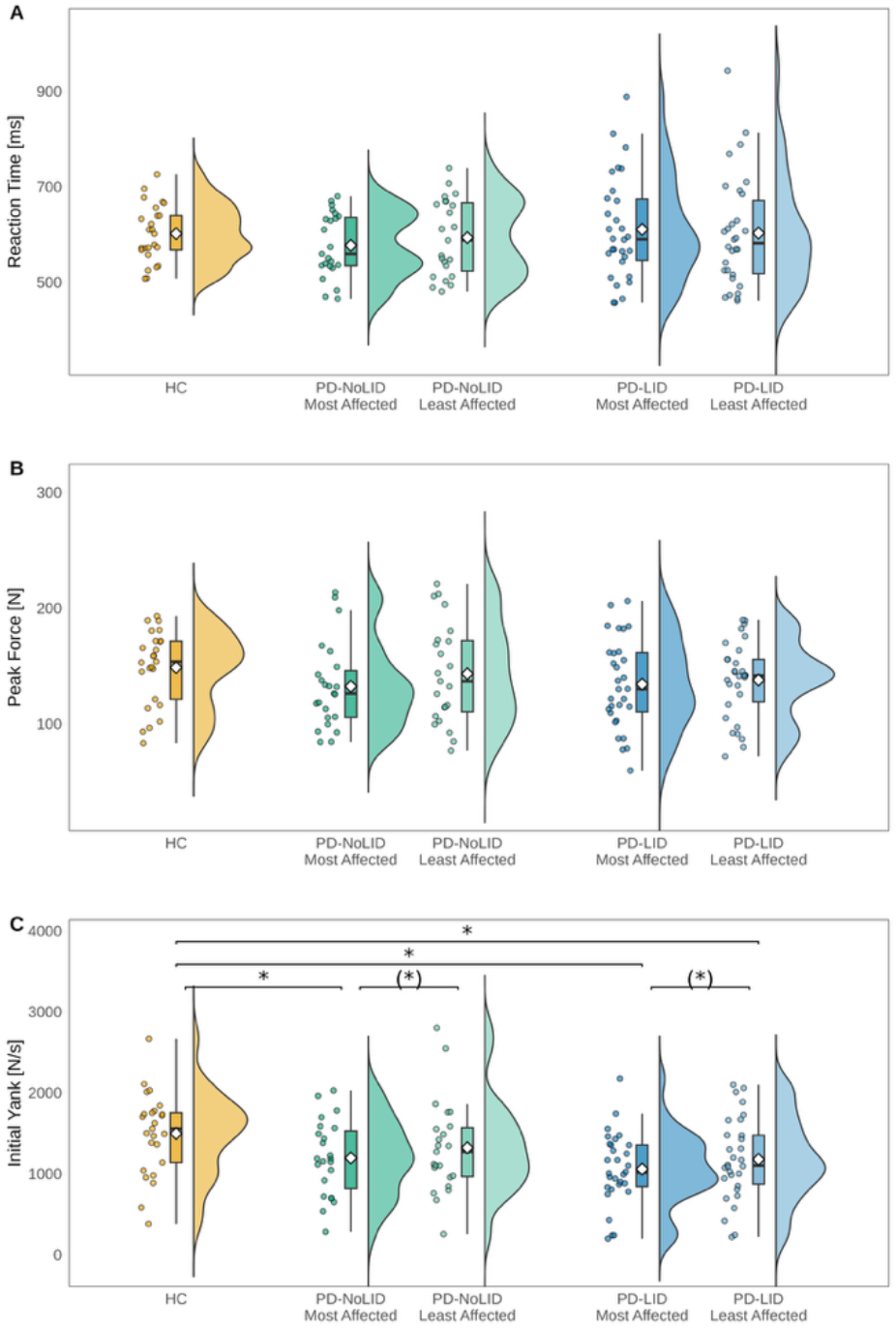
Motor Performance. A: Reaction time, B: Peak force, and C: Initial vigour. For healthy controls, values represent the mean of correct right- and left go-responses. For patients with PD, values are shown separately for the most affected and least affected sides. Individual participant means are displayed as jittered points; boxplots indicate the median and interquartile range; diamonds denote the mean; and half-violin plots represent the distribution density. Horizontal brackets indicate significant effects (*P* < 0.05). An asterisk * indicates a significant pairwise difference based on Tukey-adjusted post hoc comparisons of estimated marginal means, whereas an asterisk in parentheses (*) indicates a significant main effect of side in the ANOVA model for which the corresponding Tukey-adjusted post hoc comparisons separated for each group were not significant. Models included group, sex, and age as fixed effects. Separate models were conducted for comparisons involving the most affected side, the least affected side, and within-PD side comparisons. PD-NoLID: Patients with Parkinson’s disease without levodopa-induced dyskinesia, PD-LID: Patients with Parkinson’s disease with levodopa-induced dyskinesia.

Peak force also did not differ significantly between groups or sides (Fig. 2B). Mean peak force for HC was 149 ± 32 N, for PD-NoLID 133 ±37 N and 143 ±43 N (most/least affected), and for PD-LID 134 ±39 N and 138 ±33 N (most/least affected).

In contrast, initial vigour differed between groups (Fig. 2C). Mean initial vigour was 1498 ± 507 N/s in HC, 1200 ±464 N/s and 1320 ±579 N/s in PD-NoLID (most/least affected), and 1060 ±444 N/s and 1180 ±507 N/s in PD-LID (most/least affected). There was a group effect on the most affected side (F(2,73) = 11.24, ηp² = 0.236, *P* < 0.0001), with reduced values in both PD-NoLID (mean difference vs HC: 333 N/s, 95% CI [53.7, 611.5], t_72.6_ = 2.85, *P* = 0.0154) and PD-LID (522 N/s, 95% CI [256.6, 788.3], t_72.6_ = 4.70, *P* < 0.0001). On the least affected side, a group effect was also present (F(2,73) = 5.73, ηp² = 0.136, *P* = 0.0049), driven by reduced values in PD-LID compared to HC (426 N/s, 95% CI [124, 728.3], t_72.6_ = 3.38, *P* = 0.0033), whereas PD-NoLID did not differ significantly from HC. There were no differences between PD-NoLID and PD-LID. However, initial vigour differed between the most and least affected sides (F(1,51) = 5.67, ηp² = 0.1, *P* = 0.021) with similar side differences in PD-NoLID (127 N/s, 95% CI [−29.9, 284167], t_51_ = 1.62, *P* = 0.11) and PD-LID (120 N/s, 95% CI [−17.1, 258], t_51_ = 1.76, *P* = 0.085). Finally, initial vigour was negatively associated with OFF-medication motor MDS-UPDRS (β = −16.27, 95% CI [−25.62, −6.92], t_49_ = −3.50, adjusted R^2^ = 0.518, *P* = 0.001), indicating that reduced initial vigour was linked to greater motor impairment.

### Errors

To assess motor-control measures related to impulsivity and attention, we tested for group differences in omission and commission errors. Groups did not differ in omission errors, but for commission errors a significant group effect was observed (χ²(2) = 7.75, *P* = 0.021). Post hoc comparisons revealed that HC made fewer commission errors than PD-LID (IRR = 0.415, 95% CI [0.187, 0.922], z = −2.58, *P* = 0.026) (Fig. 3).

**Figure 3.**
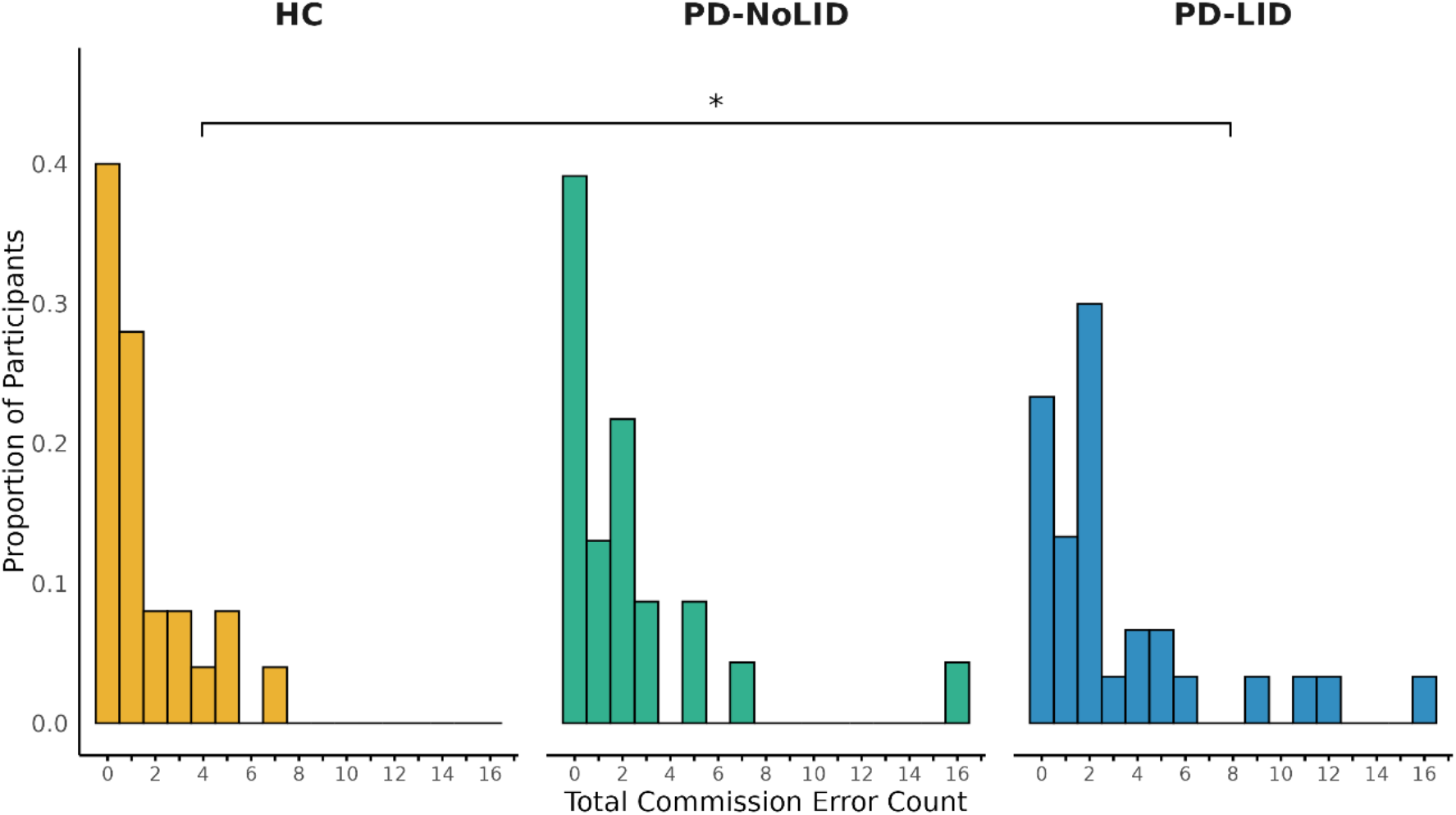
Commission Errors. Total Commission Errors (go- and no-go trials) for HC, PD-NoLID, and PD-LID. Histograms show the proportion of participants at each error count. Horizontal brackets indicate significant pairwise differences based on Tukey-adjusted post hoc comparisons following negative binomial regressions including group, sex, and age as covariates (\**P* < 0.05). HC: Healthy controls, PD-NoLID: Patients with Parkinson’s disease without levodopa-induced dyskinesia, PD-LID: Patients with Parkinson’s disease with levodopa-induced dyskinesia.

### BOLD Responses During Motor Execution

During go responses, activation was observed in the contralateral primary sensorimotor cortex (SM1), ipsilateral primary somatosensory cortex (S1), supplementary motor area (SMA) and dorsal anterior cingulate cortex (dACC), ipsilateral cerebellum lobule IV-VI, bilateral cerebellum lobule VIII, contralateral cerebellum lobule VI/Crus 1, bilateral insula, bilateral dorsal prefrontal cortex, bilateral inferior frontal gyrus (IFG), bilateral putamen, bilateral caudate bilateral thalamus (Supplementary Table 1).

ROI analyses identified significant group effects in the bilateral posterior putamen (Fig. 4). In the right posterior putamen, a significant group effect was observed during both left and right go trials (left go: (F(2,73) = 4.38, η² = 0.11, *P* = 0.016; right go: F(2,73) = 4.11, η²=0.10, *P* = 0.020). Post hoc comparisons revealed reduced BOLD responses in both PD-NoLID and PD-LID compared to HC during left go trials (PD-NoLID: mean difference = 2.54, 95% CI [0.22, 4.86], t_73_ = 2.62, *P* = 0.028; PD-LID: mean difference = 2.27, 95% CI [0.10, 4.43], t_73_ = 2.50, *P* = 0.038), whereas during right go trials, only PD-LID showed significantly reduced activity compared to HC (mean difference = 2.27, 95% CI [0.25, 4.29], t_73_ = 2.68, *P* = 0.024). There were no significant differences between PD-NoLID and PD-LID.

**Figure 4.**
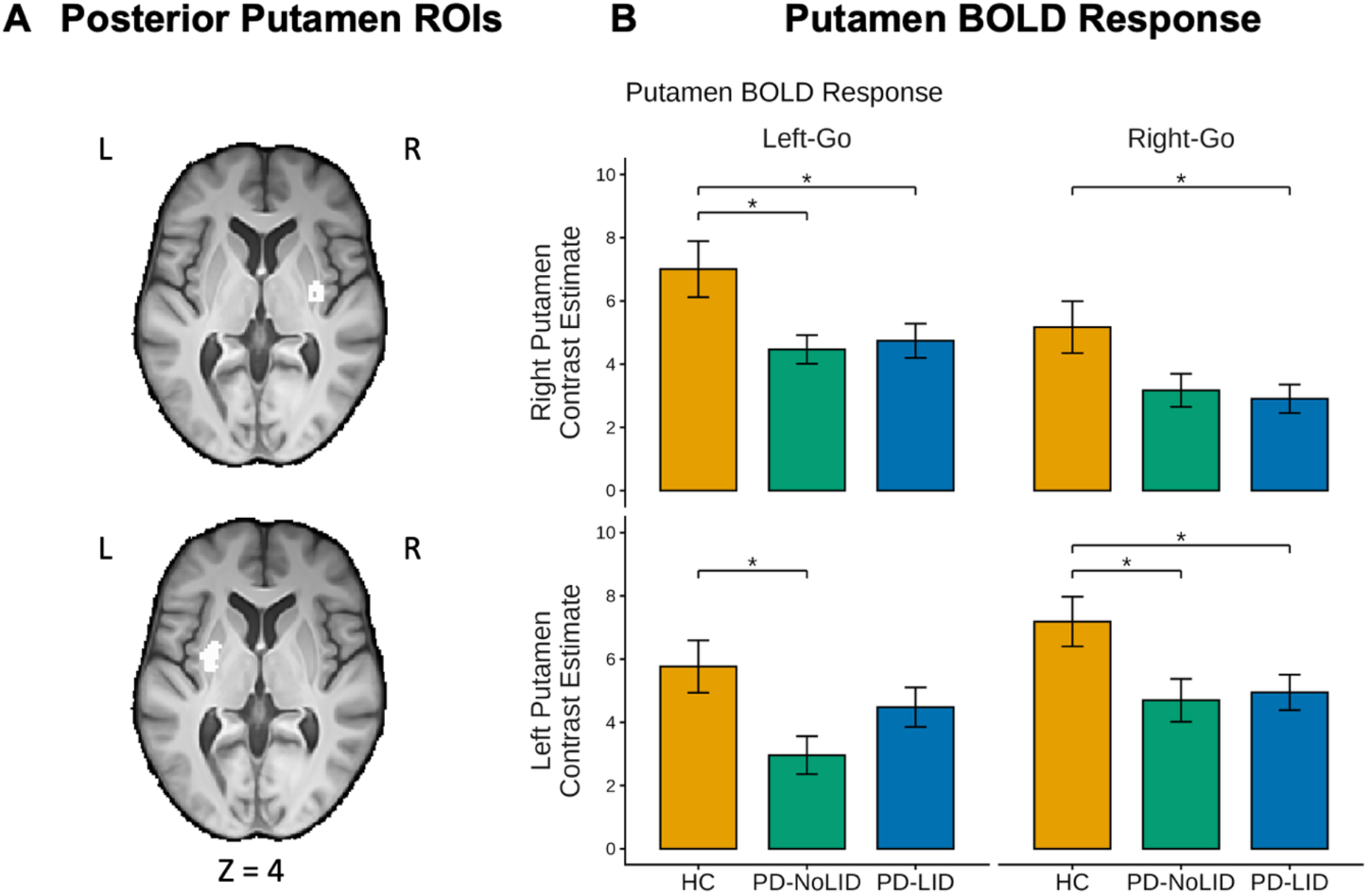
Putaminal Motor Activation. A: Bilateral posterior putamen ROI derived from the Herz et al. (2021) ALE meta-analysis^49^ shown on the study-specific template. B: Mean contrast estimates (± SEM) extracted from left and right putamen ROI during right and left go-responses. Horizontal brackets indicate significant between-group differences based on Tukey-corrected post hoc comparisons from ANOVA models including sex and age as covariates (\**P* < 0.05). HC: Healthy controls, PD-NoLID: Patients with Parkinson’s disease without levodopa-induced dyskinesia, PD-LID: Patients with Parkinson’s disease with levodopa-induced dyskinesia.

Similarly, in the left posterior putamen, significant group effects were observed during both right and left go trials (right go: F(2,73) = 4.07, η² = 0.10, *P* = 0.021; left go: F(2,73) = 3.76, η² = 0.093, *P* = 0.028). Post hoc comparisons showed reduced BOLD responses in both PD-NoLID and PD-LID compared to HC during right go trials (PD-NoLID: mean difference = 2.49, 95% CI [0.12, 4.85], t_73_ = 2.51, *P* = 0.037; PD-LID: mean difference = 2.24, 95% CI [0.030, 4.45], t_73_ = 2.43, *P* = 0.046), whereas during left go trials, only PD-NoLID showed significantly reduced activity compared to HC (mean difference = 2.80, 95% CI [0.36, 5.25], t_73_ = 2.74, *P* = 0.021; Figure 4). No differences were observed between PD-NoLID and PD-LID.

No significant group effects were observed in the other ROIs (premotor cortex, left primary motor cortex, cerebellar, SMA/preSMA or rIFG) nor in a whole-brain analysis.

### BOLD Responses During No-go

No-go trials were associated with activation of the SMA, preSMA and dACC, bilateral lateral parietal cortex, bilateral insula, bilateral IFG, bilateral dorsolateral prefrontal cortex (DLPFC), bilateral putamen, bilateral cerebellum lobule VI/Crus I, bilateral cerebellum lobule VIII/Crus II (Supplementary Table 1). No group differences were observed in the whole-brain analyses.

### Associations between Initial Vigour and Neural Activity

Given the observed behavioural differences in initial vigour, we examined whether these effects were reflected in neural activity during motor responses. However, whole-brain analyses, both probing associations with trial-by-trial variations in vigour (parametric modulation of response with trial-specific initial vigour) and with differences in interindividual average initial vigour (correlating average BOLD activity during responses with average initial vigour), did not reveal any significant effects related to initial vigour or interaction with group. Similarly, a linear mixed-effects model explaining individual average initial vigour using functional activation from the posterior putamen ROI showed no significant associations or interaction with group.

### Dopaminergic Degeneration in the Striatum

To assess whether dopaminergic degeneration differed between the PD groups, we analysed striatal DAT-specific binding ratio in 35 patients with available data using a mixed-effects linear model including group, side, and their interaction, with sex, age, and disease duration as covariates and participant as a random intercept.

A significant effect of group was observed (F(1,30) = 6.00, ηp² = 0.167, *P* = 0.020). Post hoc comparisons revealed that PD-LID had significantly lower dopamine SBR compared to PD-NoLID (mean difference = −0.25, 95% CI [−0.46, −0.041], t_30_ = −2.645, *P* = 0.020). A significant main effect of side was also observed (F(1,33) = 33.14, ηp² = 0.501, *P* < 0.0001) with lower SBR on the most affected side compared to the least affected side (mean difference = −0.21, 95% CI [−0.29, −0.14], t_33_ = −5.76, *P* < 0.0001). No significant group x side interaction was observed.

### Relationship Between Putamen BOLD Activity and Dopaminergic Integrity

We then investigated the association between the two imaging modalities. Posterior putamen BOLD activation during contralateral movements was significantly associated with dopamine SBR in posterior putamen (F(1,52) = 6.86, ηp² = 0.116, *P* = 0.011). Across groups and sides, higher posterior putamen SBR was associated with greater BOLD activation (β = 2.68, 95% CI [0.571, 4.78], t_52.9_ = 2.55, *P* = 0.014). The other factors, as well as the putamen SBR-by-side and putamen SBR-by-group interactions, were not significant (Figure 5).

**Figure 5.**
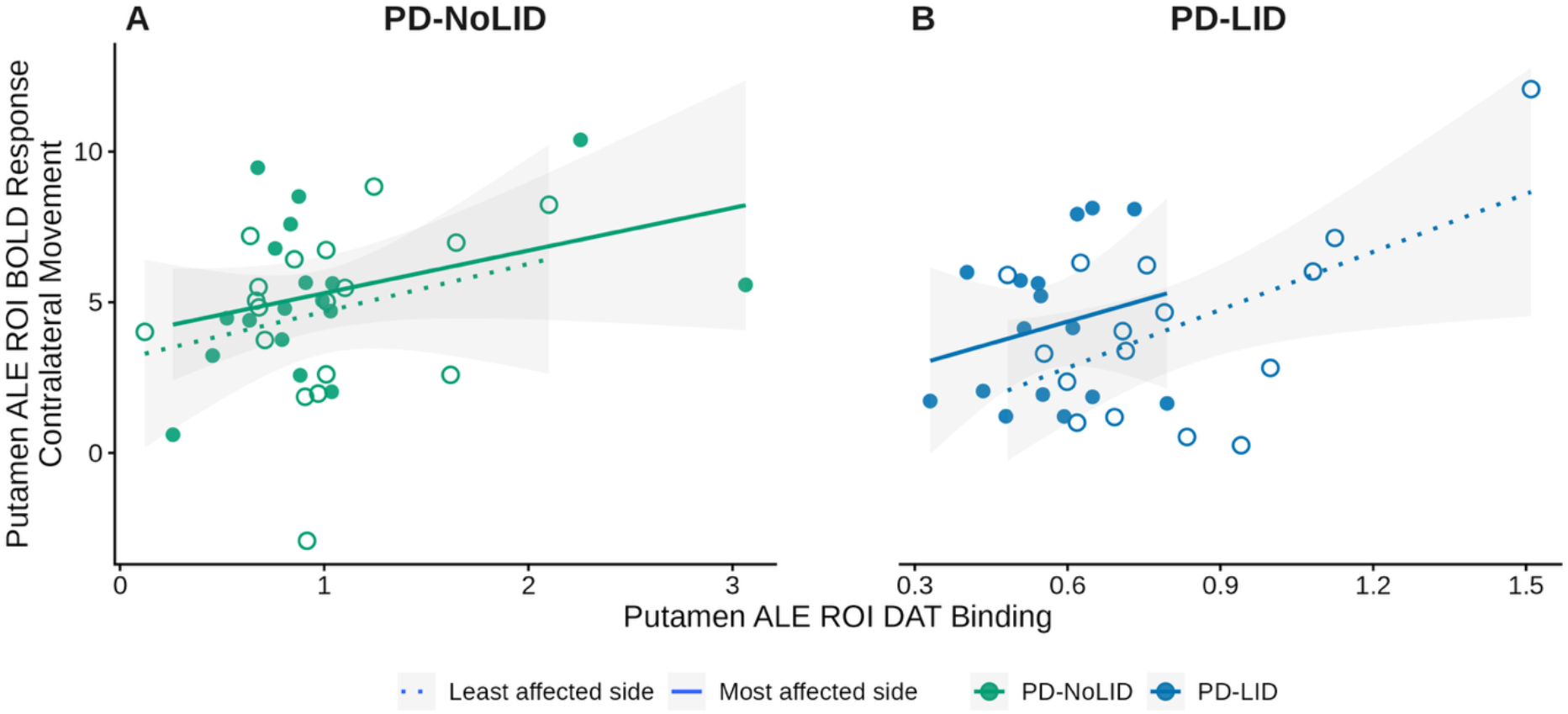
Relationship Between Putamen fMRI Activation and Dopaminergic Integrity. Scatter plots illustrating the relationship between fMRI contrast estimates during contralateral go-responses and the dopamine transporter (DAT) specific binding ratio (SBR) in the posterior putamen. Data are shown separately for PD-NoLID (left) and PD-LID (right) and include the most and least affected sides within each panel. Solid lines indicate group-specific linear regression fits for the most affected side, whereas dotted lines indicate the least affected side. PD-NoLID: Patients with Parkinson’s disease without levodopa-induced dyskinesia, PD-LID: Patients with Parkinson’s disease with levodopa-induced dyskinesia.

### Combined Associations of Motor Vigour and Imaging Measures

Lastly, to complete the link between behaviour, functional activity, and dopaminergic integrity, we performed exploratory model comparisons to identify predictors of initial vigour, including both imaging modalities and additional variables of interest (n = 34). The included predictors were contralateral putamen functional activity during movement, contralateral putaminal dopamine SBR, group, age, disease duration, sex, side, and interactions of group with contralateral putamen BOLD and contralateral putamen SBR.

The best-fitting model included contralateral putamen SBR, age, and sex as predictors (AICc = 1032.5, weight = 0.100). Higher putaminal SBR was associated with greater initial vigour (β = 522.8, 95% CI [235.4, 791.3], t_65.9_ = 3.74, *P* < 0.001). In addition, initial vigour decreased with increasing age (β = −24.4, 95% CI [−38.65, −10.14], t_30.8_ = −3.27, *P* = 0.0026), and was lower in females compared to males (β = −333.6, 95% CI [−576.2, −90.80], t_30.9_ = −2.63, *P* = 0.013). Model inclusion probabilities (i.e. the percentage of the total Akaike weight of models including the given factor^53^) supported the inclusion of age (Σwi = 0.97), sex (Σwi = 0.92), and putaminal SBR (Σwi = 0.87), whereas there was limited support for side (Σwi = 0.52), putamen BOLD activity (Σwi = 0.45), group (Σwi = 0.32), and no support for group x putaminal SBR (Σwi = 0.06) and group x putamen BOLD (Σwi = 0.03).

### Motivation and Reward

#### Effect of Motivation on Motor Output

First, we tested whether the motivational context influenced motor performance. Reaction time was significantly faster in the high-reward than in the high-loss condition (adjusted mean difference = −11.2 ms, 95% CI [−19.5, −2.9], *t*(73) = −2.69, *P* = 0.009).

Peak force showed a significant increase in the high-reward compared to the high-loss condition (adjusted mean difference = 1.56 N, 95% CI [0.47, 2.64], *t*(73) = 2.85, *P* = 0.006).

Motivational context did not significantly influence initial vigour. The effect of motivational context did not differ between groups for any of the behavioural measures.

#### Motivation-Related Modulation of BOLD Responses

After having tested whether the two reward contexts affected behaviour in the three groups, we explored corresponding context effects on the neural response to action cues. The contrast of high-reward versus high-loss motivational context was associated with activation in the occipital cortex and the left fusiform gyrus (Supplementary Table 1). We found no group effects on the neural response to motivational context.

#### Reward-Related Neural Scaling

We then tested the neural responses to reward outcomes by examining the positive linear scaling with all four possible reward outcomes. Across groups, activation increased with outcome magnitudes (high-losses < low-losses < low-wins < high-wins) in the bilateral nucleus accumbens/ventral striatum, SMA/preSMA, ventromedial prefrontal cortex (vmPFC), occipital cortex, left orbitofrontal cortex, left DLPFC, and posterior cingulate (Supplementary Table 1).

Whole-brain analyses did not reveal significant group differences. ROI analyses, however, identified group effects in reward-related regions. In the nucleus accumbens significant group effects were observed in both hemispheres (right: F(2,73) = 3.37, η² = 0.085, *P* = 0.040; left: F(2,73) = 4.19, η² = 0.10, *P* = 0.019, Figure 6). Post hoc comparisons indicated that the linear scaling was significantly reduced in the left nucleus accumbens in PD-LID compared to HC (mean difference = 3.16, 95% CI [0.32, 6.00], t_73_ = 2.66, *P* = 0.026), whereas no other group differences were significant.

**Figure 6.**
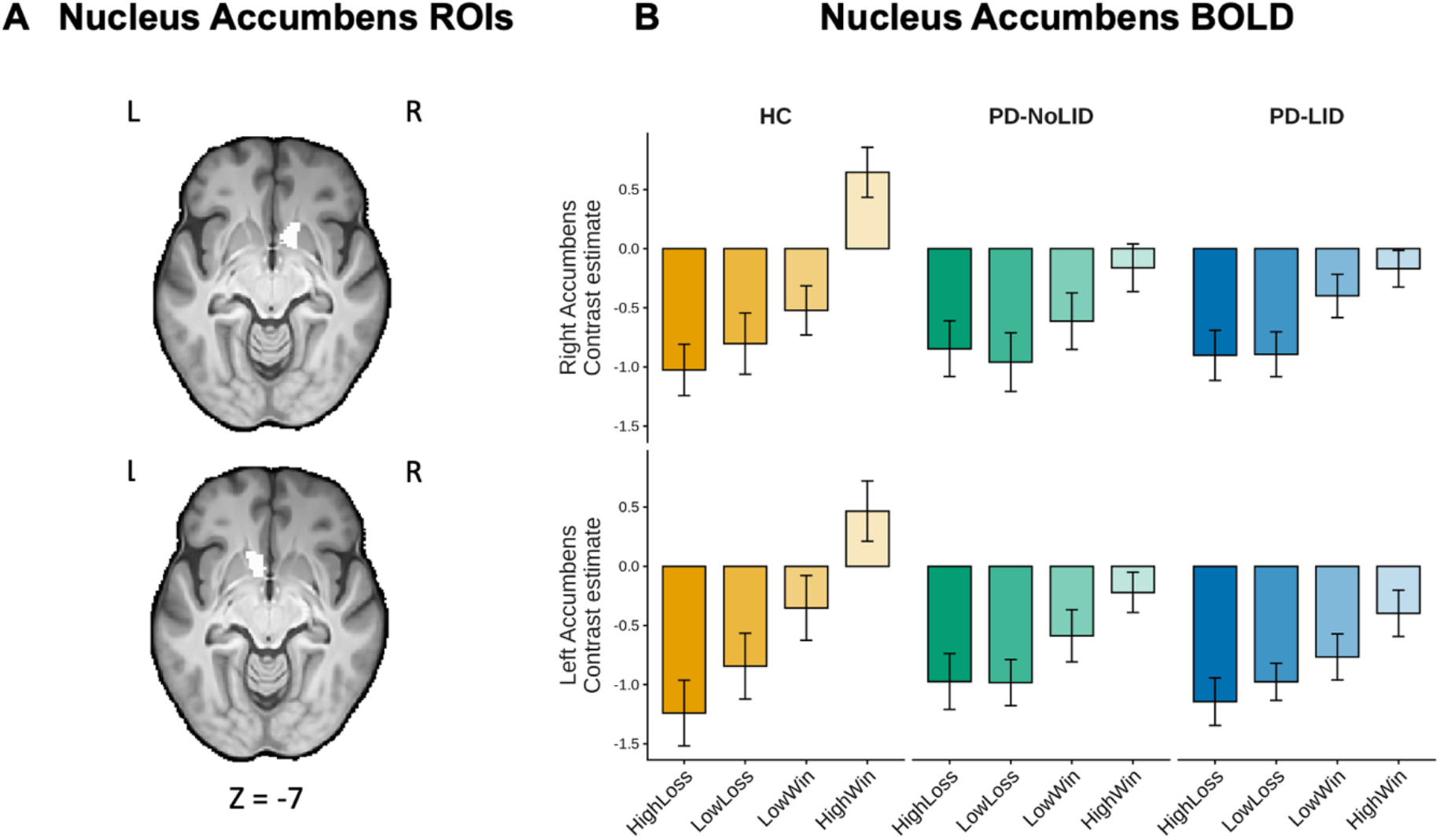
Nucleus Accumbens Activity Scaling with Reward Magnitudes. A: The right (top) and left (bottom) nucleus accumbens ROIs (Pick atlas) shown on the study specific template. B: Mean contrast estimates (± SEM) extracted from the right (top) and left (bottom) nucleus accumbens ROIs for HC, PD-NoLID, and PD-LID for each reward outcome. Contrast estimates indicate relative differences between conditions and do not represent absolute activations or deactivations. HC: Healthy controls, PD-NoLID: Patients with Parkinson’s disease without levodopa-induced dyskinesia, PD-LID: Patients with Parkinson’s disease with levodopa-induced dyskinesia.

In the left hippocampus, a significant group effect was also observed (F(2,73) = 3.57, η² = 0.089, *P* = 0.033). Post hoc comparisons showed a reduced linear scaling in PD-NoLID compared to HC (mean difference = 1.98, 95% CI [0.099, 3.86], t_73_ = 2.52, *P* = 0.037), whereas no other group differences were significant.

No significant group effects were found in the right hippocampus, amygdala, and ventral tegmental area.

#### Association Between Nucleus Accumbens Imaging Measures, Motor Performance, and Non-Motor Symptoms

Following the identification of group differences in reward-related activity, we assessed associations between clinical symptoms (MDI, LARS, BIS-11) and behavioural measures (initial vigour, delta peak force (high-reward - high loss), delta reaction time (high reward - high loss) as outcome variables, and nucleus accumbens activity and dopaminergic integrity as predictors.

We found a significant association between the linear scaling of outcome-related activity in the nucleus accumbens (averaged across hemispheres) and initial vigour (averaged across right- and left-hand trials) (F(1,68) = 7.71, ηp² = 0.10, *P* = 0.0071). The interaction with group was not significant. To quantify the strength of the association across all groups, we fit a separate model without the group-by-BOLD interaction, revealing a robust association with a large effect size (β = 25.19, 95% CI [1.35, 49.03], t_70_ = 2.11, adjusted R² = 0.363, *P* = 0.039).

No significant associations were observed between the linear scaling of the nucleus accumbens and the other behavioural or clinical measures. Also, no associations were found between dopamine SBR in nucleus accumbens and the same clinical measures. Furthermore, we found no significant association between the linear scaling of BOLD responses and the SBR in the nucleus accumbens.

## Discussion

Combining a novel reward-modulated grip-force paradigm with task-based fMRI and dopamine transporter PET, we directly linked nigrostriatal dopaminergic degeneration to both reduced movement-related recruitment of the posterior putamen and impaired motor invigoration. Reduced initial vigour emerged as a sensitive behavioural marker of motor dysfunction and was accompanied by bilateral putaminal hypoactivation during force generation. Importantly, lower putaminal DAT binding was associated with both diminished putaminal activation and reduced motor invigoration, providing the first *in vivo* evidence linking molecular dopaminergic degeneration, striatal functional recruitment, and quantitative impairment of motor invigoration within the same individuals.

Beyond motor circuit abnormalities, we identified evidence for altered motivational processing in PD. Reward-related modulation of nucleus accumbens activity was attenuated in patients and positively associated with motor invigoration across participants. In contrast, putaminal activity was not related to reward-dependent variation in vigour, suggesting a partial dissociation between dorsal striatal mechanisms supporting motor execution and ventral striatal mechanisms contributing to motivational energisation of behaviour. Together, these findings indicate that impaired motor invigoration in PD reflects both a primary consequence of dorsal striatal dopaminergic degeneration and alterations in reward processing.

### Motor Invigoration as a Mechanistic Component of Bradykinesia

Bradykinesia is the cardinal motor feature of PD and comprises several physiological abnormalities, among which impaired motor invigoration appears particularly prominent.^7–9,27^ In the present study, initial movement vigour, quantified as the first peak in the rate of force development, was markedly reduced in PD and showed lateralisation according to symptom severity. Because this measure captures the earliest phase of force generation, it likely reflects deficient scaling of the initial agonist motor command, a long-recognised physiological hallmark of bradykinesia.^12,13,59,60^ In contrast, we found no group differences in reaction time or peak force, consistent with previous reports showing considerable heterogeneity of these measures across tasks and patient cohorts.^8–10,17,61–67^ Together, these findings suggest that motor impairment in PD is not primarily characterised by delayed response initiation or reduced maximal force production, but by a diminished capacity to rapidly mobilise motor output when movement begins.

This interpretation aligns with contemporary models of basal ganglia function that assign dopamine a central role in regulating movement vigour.^24,25,27^ Within these frameworks, dopamine depletion reduces the gain of motor commands, resulting in systematic underscaling of actions. Consequently, bradykinesia may be understood not simply as motor slowing, but as a failure to appropriately invigorate movement. Our findings further indicate that initial vigour constitutes a sensitive and physiologically meaningful marker of this impairment, capturing motor deficits that are not readily detected by conventional measures such as reaction time or peak force.

### Reduced Putamen Recruitment During Motor Invigoration

Movement-related activity in the posterior putamen was significantly reduced bilaterally in PD, despite participants performing unilateral hand contractions. This reduction is consistent with previous imaging studies demonstrating that PD is associated with bilateral dysfunction of motor striatal circuits extending beyond the clinically most affected side.^49,68–72^ The posterior putamen occupies a central position within cortico-basal ganglia motor loops and plays a critical role in the scaling and execution of voluntary movement.^73,74^ Reduced recruitment of this region therefore accords well with previous task-based fMRI studies in Parkinsońs Disease^49^ and mechanistic models proposing that nigrostriatal dopamine depletion diminishes the responsiveness, or gain, of striatal motor circuits. In this framework, insufficient engagement of the posterior putamen limits the capacity of the motor system to generate vigorous actions.

Interestingly, putaminal activity was not directly associated with individual differences in motor vigour. This suggests that the movement-related BOLD response reflects a composite signal related to multiple processes involved in motor planning, execution, and monitoring rather than a specific neural representation of vigour per se. Consequently, reductions in putaminal activation may represent a systems-level signature of impaired motor circuit engagement, while behavioural measures of vigour provide a more direct readout of its functional consequences.

### Dopaminergic Degeneration Links Striatal Dysfunction to Impaired Motor Invigoration

Loss of dopaminergic input to the posterior putamen is a hallmark pathophysiological feature of PD, and reduced movement-related putaminal activation is a consistent finding in functional neuroimaging studies.^49^ Yet, the proposed relationship between nigrostriatal dopaminergic degeneration and impaired functional recruitment of the putamen has not previously been demonstrated in vivo within the same individuals. Previous studies combining DAT PET with resting-state fMRI have reported that striatal dopaminergic deficits are associated with reduced functional connectivity between the striatum and cortical and subcortical regions.^36–38^

Here, lower putaminal DAT binding was associated with reduced movement-related putaminal activation, providing *in vivo* evidence that nigrostriatal dopaminergic degeneration contributes directly to impaired functional recruitment of motor striatal circuits. Furthermore, lower DAT binding was associated with reduced initial vigour. When both DAT binding and movement-related putaminal activity were entered into the same model, dopaminergic integrity remained the dominant predictor of motor vigour. Together, these observations suggest that the behavioural consequences of dopamine loss are more closely reflected by molecular measures of striatal degeneration than by regional BOLD amplitude alone.

These findings integrate several previously independent observations. Experimental studies have demonstrated that dopamine depletion reduces movement vigour,^24,32^ while human imaging studies have linked striatal DAT loss to disease severity and disease progression.^35,38,75^ Functional MRI studies have additionally shown that bradykinesia severity correlates with reduced activation throughout the basal ganglia motor network, including the putamen.^76^ The present results extend this literature by linking dopaminergic degeneration, movement-related striatal recruitment, and quantitative measures of motor invigoration within a single multimodal framework.

Collectively, our findings support a mechanistic model in which degeneration of nigrostriatal dopaminergic input reduces the functional engagement of dorsal striatal motor circuits and thereby constrains motor invigoration. This provides a neurobiological account of bradykinesia that bridges molecular pathology, systems-level dysfunction, and behavioural impairment.

### Reward Processing and Motor Invigoration

Motor invigoration is strongly influenced by motivational state and expected action value.^18^ Accordingly, bradykinesia has increasingly been conceptualised as a disorder of action energisation.^77,78^ Dopaminergic reward prediction signals projecting to the ventral striatum play a central role in assigning value to actions and motivating behavioural output.^79^ Given that mesolimbic dopaminergic pathways also undergo degeneration in PD,^80^ abnormalities in reward processing may contribute to impaired motor invigoration alongside dysfunction of dorsal striatal motor circuits.

The prospect-of-reward manipulation successfully influenced behaviour, with larger prospective wins leading to shorter reaction times and greater peak force production. However, reward context did not significantly alter initial vigour and did not produce differential striatal activation between reward conditions. One possible explanation is that the task balanced incentives to obtain rewards and avoid losses, thereby reducing sensitivity to detect context-dependent effects on motor invigoration. Importantly, these negative findings do not exclude a role of motivational processes in shaping vigour, as revealed by the outcome-related neural responses.

Studies on alterations of reward expectation and reward outcome processing in PD have not produced definite evidence,^81^ but reduced reward responsivity has been reported in PD.^39^ The implementation of two reward contexts allowed us to explore reward expectation and outcome processing in PD and their interaction with kinematic parameters.

During reward-outcome processing, patients with PD exhibited attenuated reward-related scaling of activity in the bilateral nucleus accumbens and left hippocampus, corroborating previous evidence for reduced reward responsivity in PD.^39^ More importantly, individuals exhibiting stronger reward-related modulation of nucleus accumbens activity also demonstrated greater motor invigoration. This association was observed across groups and identifies ventral striatal reward sensitivity as an important determinant of vigorous action.

Experimental studies in animals have established causal links between mesolimbic dopamine signalling and movement vigour.^20–22,82,83^ Task-based neuroimaging studies in humans have further implicated the ventral striatum in value representation and effort-based decision-making.^84,85^ Extending this work, our findings provide evidence that reward-related recruitment of the nucleus accumbens is associated with the capacity to invigorate motor output. Together, these results suggest that impaired motor invigoration in PD arises from at least two partially dissociable mechanisms: degeneration of dorsal striatal motor circuits directly constraining movement generation, and reduced responsiveness of ventral striatal reward circuits limiting the motivational drive to energise action.

### Dyskinesia-Specific Changes

Levodopa-induced dyskinesia (LID) is thought to arise from maladaptive plasticity within cortico-basal ganglia circuits following chronic dopaminergic treatment.^86^ We therefore investigated whether patients with LID exhibit persistent behavioural or neural alterations even in the OFF-medication state.

Patients with LID committed more commission errors than healthy controls, consistent with previous reports of impaired inhibitory control in this subgroup.^87^ In addition, they showed significantly lower putaminal DAT binding than patients without LID after adjustment for demographic and disease-related factors. This finding aligns with evidence that more severe nigrostriatal degeneration increases the risk of developing dyskinesia.^88,89^

In contrast, we found no differences between PD subgroups in motor invigoration, task-related striatal activation, or reward-related neural responses. These results suggest that in the OFF-medication state, the fundamental relationships between dopaminergic degeneration, striatal recruitment, reward processing, and motor invigoration are preserved irrespective of dyskinesia status. Previous neuroimaging studies have reported functional abnormalities in motor and inhibitory networks in LID^50^ but findings across studies have been highly heterogeneous.^90^

Taken together, our findings indicate that patients with LID exhibit more pronounced nigrostriatal dopaminergic degeneration and deficits in inhibitory control, but do not show evidence for a distinct disturbance of the neural mechanisms underlying motor invigoration in the OFF-medication state.

### Conclusions

Using behavioural, functional MRI, and dopamine transporter PET measures, we provide *in vivo* evidence linking nigrostriatal dopaminergic degeneration to reduced striatal recruitment and impaired motor invigoration in Parkinson’s disease. Initial movement vigour emerged as a sensitive marker of motor dysfunction and was closely related to putaminal dopaminergic integrity, supporting the view that bradykinesia reflects a failure to appropriately scale motor output. Patients with PD additionally showed attenuated reward-related responses in ventral striatal circuitry, and reduced reward sensitivity was associated with lower motor invigoration. Together, these findings support a model in which dorsal striatal dopamine loss directly constrains motor invigoration, while ventral striatal hyporesponsivity to reward represents a parallel and partly independent contributor to behavioural dysfunction in Parkinson’s disease.

## Data Availability Statement

Whole-brain SPM results for the main effects of the contrasts of interest within each group are provided in Supplementary Table 2.

The pseudonymized data can only be shared with a formal Data Processing Agreement and formal approval by the Danish Data Protection Agency in line with the requirements of the General Data Protection Regulation.

## Acknowledgements

The authors are grateful to Weijian Liu, Kristoffer Hougaard Madsen, Søren Asp Fuglsang, Ruben Vestergaard, Esben Thade Petersen, and Henrik Lundell for technical support and statistical guidance, and to Laura Sakalauskaitéé, Maryam Noory, Trang Tran, and Lea Maria Aldenhoven for their assistance with data collection and patient recruitment.

