## Supplementary Material for "Striatal Dopaminergic Dysfunction Constrains Motor Invigoration in Parkinson’s Disease"

### Participants

All dopaminergic medication was paused for at least 12 hours prior to the experimental test session. In people with PD receiving amantadine, treatment was gradually withdrawn before the experimental session, with the final dose taken at least 90 h (six elimination half-lives) before testing.

### Clinical Measures

The most and least affected sides were defined based on lateralised motor symptoms (OFF-medication MDS-UPDRS III). In cases of symmetrical scores, classification was based on DAT imaging obtained at diagnosis, or by the reported side of symptom onset when imaging data were unavailable.

### Task

The paradigm was a mixed-event related modified Go/NoGo task that integrated action, reward and emotional cues. A single run consisted of 18 blocks that were separated by an inter-block-interval of 4 seconds with a white screen with fixation cross and subequently 2 seconds displaying a reminder of the action cues (left-pointing triangle cueing a left-hand grip ("Left-Go"), an empty triangle cueing no response ("No-Go") and a right-pointing triangle cueing a right-hand grip ("Right-Go")). Each block (duration ~29 seconds) contained 4 trials and was performed in one of two conditions: either a 'High win' or a 'High loss' prospect. These conditions were visually represented by background images of a desert or an image of a meadow, respectively (the sceneries were taken from stock media website Pixabay, resolution - 1920x1080 px., transparency was increased by 67%). The background was present through the whole duration of a block. The reward contexts were counterbalanced and randomized across a run (9 of each, maximum 3 repetitions of the same reward context in a row).

After an inter-trial-interval fixation period (2.1-2.4 seconds, jittered across trials (uniformly distributed around a mean of 2.25 s across the experiment)), one of the action cues was shown on top of an emotional face (the transparency of the action cue was set to 33% so that the obscured part of the face was still slightly visible) for 200 milliseconds. Participants had either to grip one of the devices ("Left-Go" or "Right-Go") or to withhold the response ("No-Go").

The action cues were randomized and counterbalanced across each run (maximum four repeated types of action in a row). The emotional faces were selected from the FACES database[1] and pre-processed using Adobe Photoshop CC 2019 (background removal and greyscaling). In one run, 72 faces were presented: 24 old, 24 middle-aged, 24 young persons; 12 males and 12 females in each subgroup; 4 happy, 4 sad and 4 neutral faces in each sex category. Faces were selected from the database by picking from the top 20% in the valence recognition score.[1] There were always two male and two female faces in a block.

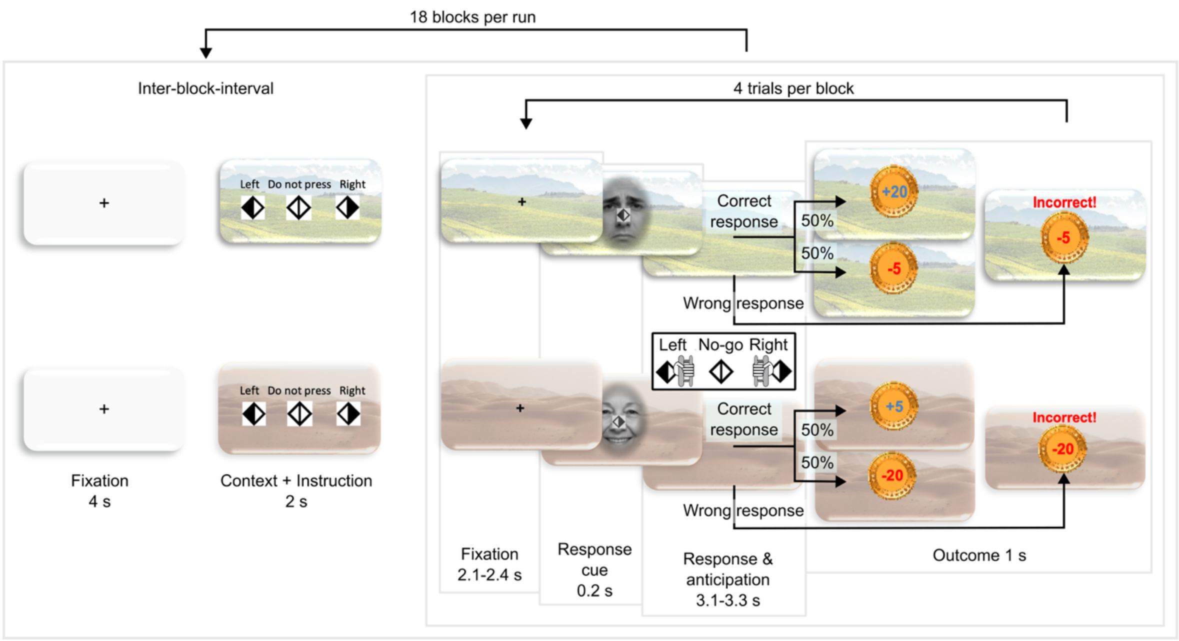


**Supplementary Figure 1. Experimental task design.** Each run consisted of 18 blocks with four trials per block. Blocks were performed in either a high-reward (meadow) or high-loss (desert) motivational context. Participants responded to left-go and right-go cues and withheld responses to no-go cues. Correct responses were followed by a monetary gain or loss with equal probability, whereas incorrect responses resulted in a monetary loss. Timing of the individual task components is shown in the figure. Due to copyright issues, the emotional faces in this figure are not from the FACES database but were created with AI (Microsoft Copilot based on GPT-5).

The face and response cue disappeared from the screen for 3.1-3.3 seconds (jittered across trials (uniformly distributed around a mean of 3.2 s across the experiment)). Within this period, participants had 2 seconds to make their response (all Go responses were permitted during a Go trial, allowing participants to correct their response (e.g., switching from left to right). However, these trials were later excluded from the analysis) followed by 1.1-1.3 seconds that were visually identical, but where responses were no longer taken into account.

This was followed by a reward/loss feedback window of 1 second. If participants had responded correctly within the first 2 seconds after cue offset, they had a 50% chance of winning or losing money. The amount of monetary gain or loss depended on the reward condition. The meadow scenery indicated that upon the correct response, participants had a 50% chance of winning 20 DKK and a 50% chance of losing 5 DKK. The desert scenery indicated the opposite – a 50% chance of winning 5 DKK and a 50% chance of losing 20 DKK. An incorrect response always resulted in a loss (a loss of 5 DKK in the high win prospect condition, and a loss of 20 DKK in the high loss prospect condition). The monetary wins and losses were pre-determined at the onset of the experiment and all outcomes were randomized and counterbalanced (maximally three win or loss outcomes in a row) such that an errorless participation resulted in a reward of 250 DKK.

The paradigm was written with Python 3.7.10 and relied on PsychoPy (version 2021.2.0).

### Behavioural Measures

Grip force was recorded on a normalised scale (0–1 AU ≈ 0–235 N), with a response threshold corresponding to a grip force of approximately 70 N.

A rectified baseline of the df/dt output was computed from the 1s interval preceding cue onset up to the cue. Reaction time was defined as the time at which the df/dt output exceeded the rectified baseline ± 2.5 standard deviations. Reaction times were log-transformed to improve normality and interpreted as proportional changes after back-transformation. Peak force was defined as the maximum force of the prompted hand. Grip-force data were processed in Python (v3.8.5) and MATLAB (vR2020a).

### Details on fMRI Preprocessing (fmriprep)

Results included in this manuscript come from preprocessing performed using fMRIPrep 25.2.4,[2,3] which is based on Nipype 1.10.0.[4,5]

**Preprocessing of B0 inhomogeneity mappings**

A total of 4 fieldmaps were found available within the input BIDS structure for this particular subject. A B0-nonuniformity map (or fieldmap) was estimated based on two (or more) echo-planar imaging (EPI) references with topup.[6]

**Anatomical data preprocessing**

A total of 1 T1-weighted (T1w) images were found within the input BIDS dataset. The T1w image was corrected for intensity non-uniformity (INU) with N4BiasFieldCorrection,[7] distributed with ANTs 2.6.2,[8] and used as T1w-reference throughout the workflow. The T1w-reference was then skull-stripped with a Nipype implementation of the antsBrainExtraction.sh workflow (from ANTs), using OASIS30ANTs as target template. Brain tissue segmentation of cerebrospinal fluid (CSF), white-matter (WM) and grey-matter (GM) was performed on the brain-extracted T1w using fast.[9] Volume-based spatial normalisation to the study-specific template was performed through nonlinear registration with antsRegistration (ANTs 2.6.2), using brain-extracted versions of both T1w reference and the T1w template.

**Functional data preprocessing**

First, a reference volume was generated, using a custom methodology of fMRIPrep, for use in head motion correction. Head-motion parameters with respect to the BOLD reference (transformation matrices, and six corresponding rotation and translation parameters) are estimated before any spatiotemporal filtering using mcflirt.[10] The estimated fieldmap was then aligned with rigid-registration to the target EPI (echo-planar imaging) reference run. The field coefficients were mapped on to the reference EPI using the transform. The BOLD reference was then co-registered to the T1w reference using mri_coreg (FreeSurfer) followed by flirt[11] with the boundary-based registration[12] cost-function. Co-registration was configured with six degrees of freedom. BOLD runs were slice-time corrected to 0.86s (0.5 of slice acquisition range 0s-1.72s) using 3dTshift from AFNI.[13] FD was computed using two formulations following Power (absolute sum of relative motions, Power et al. (2014))[14] and Jenkinson (relative root mean square displacement between affines, Jenkinson et al. (2002)).[10] FD was calculated using the implementation in Nipype (following the definitions by Power et al. 2014). The three global signals are extracted within the CSF, the WM, and the whole-brain masks.

Additionally, a set of physiological regressors were extracted to allow for component-based noise correction (CompCor, Behzadi et al. 2007).[15] Principal components are estimated after high-pass filtering the preprocessed BOLD time-series (using a discrete cosine filter with 128s cut-off) for the anatomical CompCor variant (aCompCor). For aCompCor, three probabilistic masks (CSF, WM and combined CSF+WM) are generated in anatomical space. The implementation differs from that of Behzadi et al. in that instead of eroding the masks by 2 pixels on BOLD space, a mask of pixels that likely contain a volume fraction of GM is subtracted from the aCompCor masks. This mask is obtained by thresholding the corresponding partial volume map at 0.05, and it ensures components are not extracted from voxels containing a minimal fraction of GM. Finally, these masks are resampled into BOLD space and binarized by thresholding at 0.99 (as in the original implementation). Components are also calculated separately within the WM and CSF masks. For each CompCor decomposition, the k components with the largest singular values are retained, such that the retained components' time series are sufficient to explain 50 percent of variance across the nuisance mask (CSF, WM, combined, or temporal). The remaining components are dropped from consideration. The head-motion estimates calculated in the correction step were also placed within the corresponding confounds file. The confound time series derived from head motion estimates and global signals were expanded with the inclusion of temporal derivatives and quadratic terms for each.[16] All resamplings can be performed with a single interpolation step by composing all the pertinent transformations (i.e. head-motion transform matrices, susceptibility distortion correction when available, and co-registrations to anatomical and output spaces). Gridded (volumetric) resamplings were performed using nitransforms, configured with cubic B-spline interpolation.

Many internal operations of fMRIPrep use Nilearn 0.11.1,[17] mostly within the functional processing workflow. For more details of the pipeline, see the section corresponding to workflows in fMRIPrep's documentation.

The above text was modified from the boilerplate text automatically generated by fMRIPrep to exclude irrelevant resampling targets and outputs. The boilerplate text is released under the CC0 license.

### Details on fMRI Statistical Analysis

#### Regressors of no interest

Incorrect trials and their corresponding feedback, as well as the inter-block interval (IBI), were modelled separately as regressors of no interest. Additional nuisance regressors included physiological noise regressors (heart rate and respiration), six rigid-body motion parameters (three translations and three rotations), scrubbing regressors for volumes with framewise displacement > 0.9 mm, and the first five anatomical component-based noise correction (aCompCor) components.

### Methodological considerations

It should be noted that the task did not include both high- and low-reward conditions. The two conditions were instead mirrored in valence; i.e., in terms of expected value there was an equally high motivation to avoid high losses in one condition as to achieve high rewards in the other. This might have balanced out the motivation between the two conditions, rendering the design insensitive to changes in response vigour. Accordingly, there was no difference in the fMRI response to the motor cues between the two contexts, other than in the visual cortex, which likely reflects the differences in visual properties of the background images.

It should be noted that we assumed a linear scaled response to high losses, low losses, low wins to high wins. This does not take into account that a given context only contains two of the outcomes, making comparisons across conditions difficult. However, reward contexts were grouped into short blocks of four trials, producing context changes with high frequency, and the observed response patterns do suggest a close-to-linear scaling across the two contexts.

### Supplementary Table 1

| Right Go | Area | MNI_x | MNI_y | MNI_z |
|---|---|---|---|---|
| Right Go | Left M1 | -37 | -23 | 56 |
| Right Go | Left S1 | -36 | -30 | 62 |
| Right Go | Right S1 | 60 | -18 | 38 |
| Right Go | SMA/preSMA | 0 | -1 | 51 |
| Right Go | dACC | -2 | 5 | 41 |
| Right Go | Right cerebellum IV-VI | 22 | -53 | -24 |
| Right Go | Right cerebellum VIII | 22 | -63 | -52 |
| Right Go | Left cerebellum VIII | -21 | -69 | -52 |
| Right Go | Left cerebellum VI/crus | -33 | -52 | -28 |
| Right Go | Right insula | 39 | 1 | 12 |
| Right Go | Left insula | -37 | -1 | 8 |
| Right Go | Right dorsolateral prefrontal | 45 | 39 | 33 |
| Right Go | Left dorsolateral prefrontal | -44 | 38 | 33 |
| Right Go | rIFG | 57 | 12 | 4 |
| Right Go | lIFG | -59 | 5 | 2 |
| Right Go | Left anterior/middle putamen | -25 | 7 | -1 |
| Right Go | Right caudate | 10 | 4 | 8 |
| Right Go | Left caudate | -11 | 8 | 3 |
| Right Go | Right thalamus | 14 | -14 | 9 |
| Right Go | Left thalamus | -14 | -16 | 7 |
| Right Go | Left parietal oberculum | -51 | -25 | 20 |
| Left Go | Right M1 | 34 | -27 | 62 |
| Left Go | Right S1 | 41 | -30 | 56 |
| Left Go | Left S1 | -42 | -36 | 46 |
| Left Go | SMA/preSMA | 4 | -3 | 51 |
| Left Go | dACC | 3 | 2 | 40 |
| Left Go | Left cerebellum IV-VI | -21 | -54 | -21 |
| Left Go | Left cerebellum VIII | -18 | -63 | -54 |
| Left Go | Right cerebellum VIII | 20 | -63 | -53 |
| Left Go | Right cerebellum VI/crus | 36 | -40 | -31 |
| Left Go | Right insula | 38 | 1 | 11 |
| Left Go | Left insula | -37 | 1 | 7 |
| Left Go | Right dorsolateral prefrontal | 43 | 39 | 33 |
| Left Go | Left dorsolateral prefrontal | -40 | 37 | 28 |
| Left Go | rIFG | 57 | 11 | 5 |
| Left Go | lIFG | -59 | 10 | 0 |
| Left Go | Right putamen | 26 | 2 | 3 |
| Left Go | Left putamen | -27 | 3 | -3 |
| Left Go | Right caudate | 20 | 13 | 0 |

| | | | | |
|---|---|---|---|---|
| **Left Go** | Left caudate | -22 | 8 | -1 |
| **Left Go** | Right thalamus | 15 | -15 | 6 |
| **Left Go** | Left thalamus | -14 | -14 | 9 |
| **Left Go** | Right parietal oberculum | 54 | -16 | 16 |
| **No-go** | SMA/preSMA | 3 | 2 | 71 |
| **No-go** | dACC | -6 | 5 | 45 |
| **No-go** | Right lateral parietal | 57 | -49 | 45 |
| **No-go** | Left lateral parietal | -57 | -47 | 53 |
| **No-go** | Right insula | 37 | 22 | 4 |
| **No-go** | Left insula | -40 | 19 | 3 |
| **No-go** | Right IFG | 53 | 17 | -3 |
| **No-go** | Left IFG | -57 | 5 | 2 |
| **No-go** | Right DLPFC | 42 | 39 | 34 |
| **No-go** | Left DLPFC | -48 | 33 | 30 |
| **No-go** | Right putamen | 29 | 8 | 5 |
| **No-go** | Left putamen | -30 | 10 | 4 |
| **No-go** | Right cerebellum VI/Crus I | 34 | -57 | -29 |
| **No-go** | Left cerebellum VI/Crus I | -40 | -60 | -25 |
| **No-go** | Right cerebellum VIII/Crus II | 31 | -54 | -51 |
| **No-go** | Left cerebellum VIII/Crus II | -34 | -57 | -51 |
| **Motivation** | Occipital cortex | -4 | -79 | 1 |
| **Motivation** | Left fusiform cortex | -28 | -53 | -5 |
| **Reward scaling** | Right nucleus accumbens | 19 | 11 | -10 |
| **Reward scaling** | Left nucleus accumbens | -19 | 8 | -12 |
| **Reward scaling** | SMA/preSMA | 2 | -3 | 68 |
| **Reward scaling** | dACC | 1 | -3 | 38 |
| **Reward scaling** | vmPFC | 2 | 55 | -3 |
| **Reward scaling** | Occipital cortex | 10 | -94 | 24 |
| **Reward scaling** | Left orbitofrontal cortex | -32 | 39 | -15 |
| **Reward scaling** | Left DLPFC | -39 | 46 | 27 |
| **Reward scaling** | Posterior cingulate | -4 | -33 | 49 |

**Supplementary Table 1.** Main effects of task conditions across all participants. Shown are the main effects of Right Go, Left Go, No-Go, Motivation (High Reward Go > High Loss Go), and Reward Scaling (High Loss < Low Loss < Low Win < High Win reward feedback) across all participants. Peak coordinates were transformed from the study-specific template to the MNI152NLin2009cAsym standard space. To improve readability, only selected local maxima are reported. Statistical maps were thresholded at a voxel-wise threshold of $P < 0.001$ (uncorrected) with a cluster extent threshold of >100 voxels. Coordinates are reported in MNI152NLin2009cAsym space.
