## Supplementary Table 2 for "Striatal Dopaminergic Dysfunction Constrains Motor Invigoration in Parkinson’s Disease"

HC-RightGo_OFF

| p | c | p(FWE-corr) | p(FDR-corr) | equivk | p(unc) | p(FWE-corr).1 | p(FDR-corr).1 | T | equivZ | p(unc).1 | MNI_x | MNI_y | MNI_z |
|---|---|---|---|---|---|---|---|---|---|---|---|---|---|
| **0** | 15 | 0 | 0 | 38967 | 0 | 0 | 0 | 18.2 | Inf | 0 | -56.1003 | -23.2035 | 52.2114 |
| | | | | | | 0 | 0 | 15.82 | 7.58 | 0 | -63.2621 | -28.4685 | 47.9932 |
| | | | | | | 0 | 0 | 14.89 | 7.41 | 0 | -43.0479 | -1.77507 | 14.0791 |
| | | | | | | 0 | 0 | 14.46 | 7.32 | 0 | -36.7911 | -22.4055 | 55.8538 |
| | | | | | | 0 | 0 | 14.11 | 7.25 | 0 | -37.683 | -31.4778 | 59.184 |
| | | | | | | 0 | 0 | 13.56 | 7.14 | 0 | -46.8193 | -26.3092 | 61.2465 |
| | | | | | | 0 | 0 | 13.44 | 7.11 | 0 | -50.1326 | -26.2273 | 19.9403 |
| | | | | | | 0 | 0 | 13.28 | 7.07 | 0 | -62.9859 | -18.6036 | 33.4209 |
| | | | | | | 0 | 1 | 12.28 | 6.84 | 0 | -43.2803 | 0.843153 | 2.04268 |
| | | | | | | 0 | 1 | 11.91 | 6.75 | 0 | -40.8618 | -40.1553 | 45.7475 |
| | | | | | | 0 | 1 | 11.87 | 6.74 | 0 | -42.0718 | -16.5556 | 23.5977 |
| | | | | | | 0 | 2 | 11.38 | 6.61 | 0 | -42.3022 | -13.3137 | 62.6665 |
| | | | | | | 0 | 3 | 11.05 | 6.53 | 0 | -45.6843 | -29.4657 | 48.116 |
| | | | | | | 0 | 4 | 10.88 | 6.48 | 0 | -59.5102 | 14.4113 | -2.58137 |
| | | | | | | 0 | 5 | 10.6 | 6.4 | 0 | -56.9174 | -21.6881 | 17.419 |
| | | | | | | 0 | 6 | 10.45 | 6.36 | 0 | -63.5221 | -31.7837 | 29.3527 |
| | | | | | | 0 | 6 | 10.38 | 6.34 | 0 | -56.6502 | 5.38006 | 35.1987 |
| | | | | | | 0 | 6 | 10.36 | 6.33 | 0 | -41.3692 | -44.0879 | 57.4962 |
| | | | | | | 0 | 7 | 10.22 | 6.29 | 0 | -65.5127 | -25.0935 | 23.5437 |
| | | | | | | 0 | 7 | 10.2 | 6.28 | 0 | -32.3008 | -18.9264 | 5.04838 |
| | | | | | | 0 | 9 | 10 | 6.22 | 0 | -40.5114 | -36.5561 | 69.9112 |
| | | | | | | 0 | 11 | 9.76 | 6.15 | 0 | -49.6209 | -43.6186 | 63.4007 |
| | | | | | | 0 | 14 | 9.58 | 6.09 | 0 | -50.829 | 3.66877 | 9.0615 |
| | | | | | | 0 | 15 | 9.54 | 6.08 | 0 | -33.0136 | 8.10478 | 5.57707 |
| | | | | | | 0 | 15 | 9.52 | 6.07 | 0 | -58.4648 | -40.5231 | 27.3595 |
| | | | | | | 0 | 24 | 9.14 | 5.95 | 0 | -54.3416 | -35.297 | 56.0951 |
| | | | | | | 1 | 35 | 8.89 | 5.86 | 0 | -28.092 | -2.1882 | -10.5129 |
| | | | | | | 1 | 48 | 8.67 | 5.78 | 0 | -60.1157 | -44.3739 | 50.767 |
| | | | | | | 1 | 54 | 8.59 | 5.75 | 0 | -28.5577 | 2.07945 | -2.07927 |
| | | | | | | 2 | 0.07 | 8.4 | 5.68 | 0 | -26.3777 | -51.8164 | 67.4722 |
| | | | | | | 2 | 73 | 8.36 | 5.67 | 0 | -26.8659 | -8.57843 | 62.9135 |
| | | | | | | 2 | 75 | 8.33 | 5.66 | 0 | -26.5198 | -15.6769 | 74.1559 |
| | | | | | | 2 | 85 | 8.24 | 5.62 | 0 | -57.3108 | 8.49632 | 14.8493 |
| | | | | | | 2 | 87 | 8.21 | 5.61 | 0 | -57.9777 | 2.47824 | 0.940711 |
| | | | | | | 3 | 92 | 8.17 | 5.6 | 0 | -16.3796 | -2.96184 | 12.1719 |
| | | | | | | 3 | 102 | 8.09 | 5.57 | 0 | -30.1502 | -10.5475 | 10.911 |
| | | | | | | 5 | 156 | 7.8 | 5.45 | 0 | -34.8009 | -19.0765 | 73.2648 |
| | | | | | | 6 | 175 | 7.72 | 5.42 | 0 | -37.148 | -46.2089 | 37.9689 |
| | | | | | | 6 | 175 | 7.71 | 5.42 | 0 | -40.2172 | 16.0568 | -2.23399 |
| | | | | | | 8 | 211 | 7.6 | 5.37 | 0 | -29.7809 | -44.7576 | 75.5856 |
| | | | | | | 17 | 416 | 7.18 | 5.2 | 0 | -62.8686 | 8.43873 | 6.63729 |
| | | | | | | 18 | 432 | 7.14 | 5.18 | 0 | -56.5351 | -9.91558 | 14.6594 |
| | | | | | | 0.03 | 632 | 6.88 | 5.06 | 0 | -49.7995 | 1.51798 | 26.9817 |
| | | 0 | 0 | 5754 | 0 | 0 | 0 | 14.49 | 7.33 | 0 | 17.6203 | -60.1942 | -56.9995 |
| | | | | | | 0 | 6 | 10.47 | 6.36 | 0 | 10.8607 | -76.6533 | -49.4875 |
| | | | | | | 0 | 6 | 10.44 | 6.35 | 0 | 28.6768 | -55.5945 | -52.4987 |
| | | | | | | 0 | 21 | 9.26 | 5.99 | 0 | 22.7596 | -46.2573 | -53.0364 |
| | | | | | | 0 | 24 | 9.13 | 5.94 | 0 | 4.44226 | -69.1165 | -34.5665 |
| | | | | | | 1 | 54 | 8.59 | 5.75 | 0 | 27.6377 | -39.8311 | -49.4105 |

| | | | | | | | | | | | | | |
|---|---|---|---|---|---|---|---|---|---|---|---|---|---|
| | | 0 | 0 | 13692 | 0 | 0 | 1 | 12.41 | 6.88 | 0 | 56.1064 | -16.8692 | 17.6453 |
| | | | | | | 0 | 1 | 12.32 | 6.85 | 0 | 47.2997 | -33.1929 | 51.9884 |
| | | | | | | 0 | 2 | 11.35 | 6.61 | 0 | 63.8861 | -31.3594 | 42.2059 |
| | | | | | | 0 | 3 | 11.02 | 6.52 | 0 | 63.466 | -17.7705 | 38.6489 |
| | | | | | | 0 | 6 | 10.35 | 6.33 | 0 | 60.9651 | -26.8257 | 19.2496 |
| | | | | | | 0 | 7 | 10.15 | 6.27 | 0 | 66.5276 | -14.7334 | 20.1609 |
| | | | | | | 0 | 9 | 9.91 | 6.19 | 0 | 35.5286 | -36.7474 | 46.3794 |
| | | | | | | 2 | 73 | 8.37 | 5.67 | 0 | 60.5848 | -35.0377 | 27.292 |
| | | | | | | 2 | 73 | 8.35 | 5.67 | 0 | 50.7766 | -27.2901 | 27.8034 |
| | | | | | | 2 | 83 | 8.25 | 5.63 | 0 | 66.5714 | -39.9829 | 34.239 |
| | | | | | | 3 | 102 | 8.08 | 5.57 | 0 | 43.1698 | -28.9385 | 22.3357 |
| | | | | | | 4 | 114 | 7.99 | 5.53 | 0 | 57.4926 | -45.6466 | 53.5908 |
| | | | | | | 6 | 161 | 7.77 | 5.44 | 0 | 56.5808 | -17.6332 | 56.1114 |
| | | | | | | 18 | 432 | 7.14 | 5.18 | 0 | 50.4756 | -34.664 | 18.5062 |
| | | | | | | 23 | 517 | 7.02 | 5.13 | 0 | 49.5321 | -39.4391 | 63.2417 |
| | | 0 | 0 | 9900 | 0 | 0 | 2 | 11.57 | 6.66 | 0 | 36.5188 | 2.81414 | 12.2704 |
| | | | | | | 0 | 3 | 11.12 | 6.54 | 0 | 59.4885 | 17.801 | -0.564816 |
| | | | | | | 0 | 7 | 10.14 | 6.26 | 0 | 11.9144 | 6.06668 | 6.01588 |
| | | | | | | 0 | 9 | 10 | 6.22 | 0 | 51.4002 | 12.7652 | 11.3336 |
| | | | | | | 0 | 9 | 9.97 | 6.21 | 0 | 41.1588 | 0.647487 | -1.11676 |
| | | | | | | 0 | 12 | 9.71 | 6.13 | 0 | 19.6334 | 21.8178 | 4.42556 |
| | | | | | | 0 | 15 | 9.5 | 6.06 | 0 | 59.9557 | 12.7782 | 36.0294 |
| | | | | | | 0 | 17 | 9.45 | 6.05 | 0 | 46.2901 | 18.3342 | -3.29707 |
| | | | | | | 0 | 17 | 9.42 | 6.04 | 0 | 62.6858 | 11.8625 | 26.4189 |
| | | | | | | 1 | 41 | 8.76 | 5.81 | 0 | 37.9043 | -11.1746 | 12.771 |
| | | | | | | 2 | 69 | 8.42 | 5.69 | 0 | 26.1855 | 9.30569 | -3.24933 |
| | | | | | | 2 | 0.07 | 8.4 | 5.69 | 0 | 53.8321 | 18.5268 | -8.0506 |
| | | | | | | 2 | 75 | 8.33 | 5.66 | 0 | 23.8766 | 0.942475 | -11.6421 |
| | | | | | | 3 | 107 | 8.05 | 5.55 | 0 | 48.0523 | 0.587399 | 5.96932 |
| | | | | | | 6 | 166 | 7.75 | 5.43 | 0 | 32.1872 | 24.6209 | 9.86368 |
| | | | | | | 9 | 0.25 | 7.5 | 5.33 | 0 | 42.2742 | -11.073 | -5.09164 |
| | | | | | | 0.01 | 275 | 7.44 | 5.31 | 0 | 28.9688 | -2.83908 | 7.43234 |
| | | 0 | 0 | 9526 | 0 | 0 | 2 | 11.43 | 6.63 | 0 | -5.66536 | -6.5936 | 53.1922 |
| | | | | | | 0 | 3 | 10.92 | 6.49 | 0 | -7.61376 | -27.494 | 51.9551 |
| | | | | | | 0 | 6 | 10.44 | 6.35 | 0 | 0.871891 | 8.81825 | 52.8609 |
| | | | | | | 0 | 0.01 | 9.88 | 6.18 | 0 | -3.50944 | 0.652664 | 48.834 |
| | | | | | | 0 | 14 | 9.59 | 6.09 | 0 | 5.5604 | 3.28646 | 64.7082 |
| | | | | | | 0 | 17 | 9.39 | 6.03 | 0 | -6.04278 | 7.85753 | 40.6323 |
| | | | | | | 0 | 17 | 9.39 | 6.03 | 0 | -6.5789 | -16.3701 | 45.6103 |
| | | | | | | 1 | 41 | 8.78 | 5.82 | 0 | 3.02245 | 0.631349 | 38.292 |
| | | | | | | 1 | 56 | 8.56 | 5.74 | 0 | 3.19938 | 9.59457 | 42.7239 |
| | | | | | | 2 | 76 | 8.3 | 5.65 | 0 | -10.8899 | -8.9019 | 71.8847 |
| | | | | | | 3 | 0.11 | 8.02 | 5.54 | 0 | -0.415545 | 17.55 | 32.4764 |
| | | | | | | 4 | 129 | 7.92 | 5.5 | 0 | 3.30654 | -9.36433 | 71.0122 |
| | | | | | | 0.01 | 275 | 7.44 | 5.31 | 0 | 7.81358 | -0.83719 | 55.232 |
| | | | | | | 11 | 278 | 7.42 | 5.3 | 0 | -2.66664 | -17.7986 | 57.2889 |
| | | | | | | 29 | 612 | 6.9 | 5.07 | 0 | 8.63769 | 29.0424 | 35.046 |
| | | | | | | 33 | 688 | 6.83 | 5.04 | 0 | 5.60297 | -19.254 | 48.0487 |
| | | 0 | 0 | 4395 | 0 | 0 | 4 | 10.84 | 6.47 | 0 | 22.5687 | -51.4819 | -23.6588 |
| | | | | | | 0 | 17 | 9.44 | 6.04 | 0 | 39.0157 | -49.9704 | -26.9127 |
| | | | | | | 5 | 148 | 7.83 | 5.47 | 0 | 2.82019 | -62.6916 | -16.2853 |

| | | | | | | | | | | | | | |
|---|---|---|---|---|---|---|---|---|---|---|---|---|---|
| | | 0 | 0 | 563 | 0 | 0 | 9 | 9.93 | 6.2 | 0 | 16.7805 | -10.1483 | 10.8018 |
| | | | | | | 26 | 572 | 6.95 | 5.09 | 0 | 5.9603 | -6.09467 | 9.57422 |
| | | 0 | 0 | 1141 | 0 | 1 | 31 | 8.98 | 5.89 | 0 | -15.4507 | -15.1878 | 8.89344 |
| | | | | | | 1 | 46 | 8.69 | 5.79 | 0 | -15.6485 | -25.924 | 7.79369 |
| | | | | | | 5 | 0.16 | 7.78 | 5.45 | 0 | -2.09653 | -21.2425 | 8.84985 |
| | | | | | | 14 | 366 | 7.27 | 5.23 | 0 | -14.7881 | -15.17 | -1.02928 |
| | | 0 | 0 | 529 | 0 | 2 | 67 | 8.44 | 5.7 | 0 | -20.8261 | -68.3466 | -52.1562 |
| | | 0 | 3 | 138 | 1 | 2 | 75 | 8.32 | 5.66 | 0 | -10.9401 | 4.51065 | 1.67667 |
| | | 0 | 0 | 309 | 0 | 4 | 112 | 8.01 | 5.54 | 0 | -30.7985 | 19.9464 | 8.90048 |
| | | 0 | 1 | 183 | 0 | 5 | 153 | 7.82 | 5.46 | 0 | 32.986 | -9.12297 | 67.1549 |
| | | | | | | 25 | 547 | 6.98 | 5.11 | 0 | 48.6744 | -6.6635 | 61.1635 |
| | | 0 | 5 | 118 | 2 | 7 | 196 | 7.64 | 5.39 | 0 | -40.2515 | -59.5112 | -23.2797 |
| | | 0 | 3 | 135 | 1 | 9 | 238 | 7.53 | 5.34 | 0 | -53.8133 | -66.3094 | 9.14382 |
| | | 0 | 3 | 137 | 1 | 17 | 416 | 7.18 | 5.2 | 0 | -33.5645 | -50.7702 | -28.2784 |
| | | | | | | 25 | 549 | 6.97 | 5.11 | 0 | -35.1039 | -42.8558 | -28.7107 |

PDnoLID-RightGo_OFF

| p | c | p(FWE-corr) | p(FDR-corr) | equivk | p(unc) | p(FWE-corr).1 | p(FDR-corr).1 | T | equivZ | p(unc).1 | MNI_x | MNI_y | MNI_z |
|---|---|---|---|---|---|---|---|---|---|---|---|---|---|
| 0 | 17 | 0 | 0 | 10151 | 0 | 0 | 0 | 23.65 | Inf | 0 | 61.6583 | -16.653 | 24.3155 |
| | | | | | | 0 | 0 | 16.26 | 7.33 | 0 | 53.0922 | -27.4366 | 44.5744 |
| | | | | | | 0 | 1 | 15.21 | 7.16 | 0 | 51.5015 | -20.7904 | 15.8354 |
| | | | | | | 0 | 1 | 14.63 | 7.05 | 0 | 57.0883 | -14.573 | 35.5925 |
| | | | | | | 0 | 7 | 12.05 | 6.53 | 0 | 44.8732 | -32.2175 | 43.516 |
| | | | | | | 0 | 12 | 11.2 | 6.32 | 0 | 62.5855 | -26.5241 | 26.2274 |
| | | | | | | 0 | 14 | 10.94 | 6.26 | 0 | 57.4227 | -28.419 | 53.8963 |
| | | | | | | 0 | 14 | 10.91 | 6.25 | 0 | 33.8481 | -32.5502 | 40.4165 |
| | | | | | | 0 | 14 | 10.89 | 6.24 | 0 | 59.4839 | -39.4422 | 43.1672 |
| | | | | | | 0 | 16 | 10.73 | 6.2 | 0 | 54.5705 | -19.6573 | 53.5205 |
| | | | | | | 0 | 23 | 10.37 | 6.11 | 0 | 47.4007 | -45.5433 | 45.9351 |
| | | | | | | 1 | 34 | 10.02 | 6.01 | 0 | 59.4279 | -9.55439 | 14.4964 |
| | | | | | | 1 | 51 | 9.65 | 5.9 | 0 | 50.2554 | -35.9094 | 54.5395 |
| | | | | | | 1 | 52 | 9.62 | 5.89 | 0 | 54.3239 | -33.7384 | 29.8289 |
| | | | | | | 3 | 96 | 9.11 | 5.74 | 0 | 47.031 | -45.5928 | 59.5194 |
| | | | | | | 0.02 | 455 | 7.81 | 5.29 | 0 | 35.8845 | -39.9921 | 46.0488 |
| | | | | | | 28 | 614 | 7.6 | 5.21 | 0 | 63.7715 | -28.6419 | 37.2012 |
| | | | | | | 0.03 | 642 | 7.57 | 5.2 | 0 | 60.3216 | -43.5279 | 33.9861 |
| | | 0 | 0 | 27556 | 0 | 0 | 0 | 17.27 | 7.49 | 0 | -49.8958 | -39.6467 | 47.2686 |
| | | | | | | 0 | 1 | 14.89 | 7.1 | 0 | -51.4747 | -25.3884 | 21.5137 |
| | | | | | | 0 | 2 | 14.04 | 6.94 | 0 | -55.9831 | 5.07094 | 21.5989 |
| | | | | | | 0 | 2 | 13.76 | 6.89 | 0 | -41.885 | -2.31964 | 3.36709 |
| | | | | | | 0 | 3 | 13.38 | 6.81 | 0 | -49.0185 | -32.2181 | 62.1708 |
| | | | | | | 0 | 4 | 12.88 | 6.71 | 0 | -59.0515 | -28.841 | 45.7042 |
| | | | | | | 0 | 5 | 12.39 | 6.6 | 0 | -41.3777 | -11.4702 | 2.11801 |
| | | | | | | 0 | 7 | 12.15 | 6.55 | 0 | -61.9604 | -21.4393 | 23.6772 |
| | | | | | | 0 | 7 | 12.06 | 6.53 | 0 | -42.7926 | -50.8354 | 54.8493 |
| | | | | | | 0 | 7 | 11.97 | 6.51 | 0 | -34.2986 | -15.7873 | 7.704 |
| | | | | | | 0 | 8 | 11.85 | 6.48 | 0 | -60.5247 | -28.4806 | 31.2112 |
| | | | | | | 0 | 9 | 11.67 | 6.44 | 0 | -34.552 | -1.53485 | 12.5072 |
| | | | | | | 0 | 9 | 11.66 | 6.43 | 0 | -42.0431 | -31.8466 | 49.0798 |
| | | | | | | 0 | 9 | 11.59 | 6.42 | 0 | -41.7228 | -26.028 | 22.8185 |
| | | | | | | 0 | 9 | 11.56 | 6.41 | 0 | -37.3603 | -43.2777 | 42.5352 |
| | | | | | | 0 | 11 | 11.39 | 6.37 | 0 | -51.0422 | -17.8324 | 39.2821 |
| | | | | | | 0 | 11 | 11.38 | 6.37 | 0 | -46.6738 | -0.596543 | 11.8017 |
| | | | | | | 0 | 11 | 11.26 | 6.34 | 0 | -59.3841 | 12.4358 | -2.33943 |
| | | | | | | 0 | 12 | 11.13 | 6.31 | 0 | -37.6359 | -36.3134 | 67.9881 |
| | | | | | | 0 | 14 | 10.91 | 6.25 | 0 | -58.0862 | -14.6114 | 31.1407 |
| | | | | | | 0 | 14 | 10.88 | 6.24 | 0 | -42.6559 | 13.6523 | 6.9833 |
| | | | | | | 0 | 15 | 10.81 | 6.22 | 0 | -36.9559 | -19.7495 | 62.1755 |
| | | | | | | 0 | 16 | 10.79 | 6.22 | 0 | -33.2449 | -30.808 | 57.774 |
| | | | | | | 0 | 0.02 | 10.51 | 6.14 | 0 | -33.3029 | 20.3685 | 7.8243 |
| | | | | | | 1 | 36 | 9.96 | 5.99 | 0 | -50.4133 | 16.662 | 0.314614 |
| | | | | | | 1 | 0.04 | 9.86 | 5.96 | 0 | -59.7145 | 8.48073 | 33.8669 |
| | | | | | | 1 | 47 | 9.72 | 5.92 | 0 | -60.5667 | -47.8705 | 43.2057 |
| | | | | | | 2 | 88 | 9.19 | 5.76 | 0 | -61.1645 | -13.5962 | 17.5264 |
| | | | | | | 4 | 132 | 8.85 | 5.65 | 0 | -56.0627 | -45.6167 | 52.0164 |
| | | | | | | 4 | 149 | 8.75 | 5.62 | 0 | -57.2779 | -18.589 | 53.2186 |
| | | | | | | 5 | 161 | 8.67 | 5.59 | 0 | -34.2149 | -23.283 | 51.5487 |
| | | | | | | 5 | 164 | 8.65 | 5.59 | 0 | -65.171 | -48.1803 | 33.7377 |

| | | | | | | | | | | | |
|---|---|---|---|---|---|---|---|---|---|---|---|
| | | | | 6 | 0.17 | 8.58 | 5.56 | 0 | -42.904 | -21.548 | 69.971 |
| | | | | 8 | 211 | 8.4 | 5.5 | 0 | -38.0579 | -46.5706 | 35.3259 |
| | | | | 14 | 337 | 8.05 | 5.38 | 0 | -47.0545 | -42.5036 | 65.0931 |
| | | | | 17 | 397 | 7.91 | 5.33 | 0 | -37.2518 | -27.3805 | 74.03 |
| | | | | 22 | 495 | 7.76 | 5.27 | 0 | -35.1484 | 6.2051 | -8.84595 |
| | | | | 22 | 495 | 7.75 | 5.27 | 0 | -43.2299 | -14.5904 | 63.6483 |
| 0 | 0 | 6606 | 0 | 0 | 0 | 16.1 | 7.31 | 0 | 46.0893 | 1.37788 | 10.9584 |
| | | | | 0 | 4 | 12.76 | 6.68 | 0 | 60.6611 | 12.9606 | 32.4361 |
| | | | | 0 | 7 | 12.04 | 6.52 | 0 | 57.9158 | 10.3288 | 23.4171 |
| | | | | 0 | 7 | 12.03 | 6.52 | 0 | 37.54 | 7.59855 | -0.142444 |
| | | | | 0 | 7 | 11.92 | 6.5 | 0 | 47.1483 | 15.1759 | 0.795917 |
| | | | | 2 | 63 | 9.45 | 5.84 | 0 | 55.9386 | 10.5417 | 6.3223 |
| | | | | 3 | 96 | 9.11 | 5.74 | 0 | 41.1874 | -5.8214 | 0.0815798 |
| | | | | 7 | 0.19 | 8.48 | 5.53 | 0 | 57.369 | 9.14913 | -2.34771 |
| | | | | 24 | 0.52 | 7.72 | 5.26 | 0 | 32.7771 | 25.4585 | 3.78896 |
| 0 | 0 | 8320 | 0 | 0 | 2 | 14.28 | 6.99 | 0 | 3.9775 | 16.6837 | 45.8982 |
| | | | | 0 | 4 | 12.96 | 6.73 | 0 | -8.98009 | -23.3582 | 51.23 |
| | | | | 0 | 4 | 12.7 | 6.67 | 0 | 0.438237 | 26.8678 | 36.6804 |
| | | | | 0 | 5 | 12.58 | 6.65 | 0 | -0.543683 | -0.372795 | 40.974 |
| | | | | 0 | 15 | 10.81 | 6.22 | 0 | 6.10941 | 17.9496 | 33.6138 |
| | | | | 1 | 34 | 10 | 6 | 0 | 3.66817 | 6.52824 | 65.9477 |
| | | | | 1 | 0.04 | 9.86 | 5.96 | 0 | 1.67513 | 32.6679 | 28.8336 |
| | | | | 1 | 43 | 9.79 | 5.94 | 0 | 0.806027 | -1.00003 | 60.1479 |
| | | | | 5 | 151 | 8.74 | 5.62 | 0 | 6.1568 | -0.0283372 | 75.269 |
| | | | | 5 | 161 | 8.66 | 5.59 | 0 | -8.16724 | 21.6393 | 27.3927 |
| | | | | 5 | 164 | 8.62 | 5.58 | 0 | 6.3582 | 5.46616 | 54.5303 |
| | | | | 12 | 292 | 8.16 | 5.42 | 0 | -8.62993 | -12.7973 | 59.1528 |
| | | | | 19 | 431 | 7.85 | 5.31 | 0 | 4.83889 | 17.8723 | 62.3797 |
| 0 | 0 | 4678 | 0 | 0 | 2 | 13.93 | 6.92 | 0 | 16.7241 | -51.2279 | -24.7995 |
| | | | | 0 | 8 | 11.8 | 6.47 | 0 | 24.9728 | -67.9705 | -24.1623 |
| | | | | 1 | 34 | 10.03 | 6.01 | 0 | 45.1444 | -59.0767 | -28.023 |
| | | | | 1 | 42 | 9.83 | 5.95 | 0 | 28.3315 | -57.4578 | -23.6135 |
| | | | | 1 | 56 | 9.55 | 5.87 | 0 | 36.3705 | -75.4492 | -25.704 |
| | | | | 2 | 91 | 9.16 | 5.75 | 0 | 35.1878 | -38.3312 | -31.2056 |
| | | | | 3 | 109 | 9 | 5.7 | 0 | 18.3155 | -62.7987 | -17.6591 |
| | | | | 5 | 161 | 8.66 | 5.59 | 0 | 19.1396 | -75.8755 | -26.042 |
| | | | | 13 | 317 | 8.1 | 5.4 | 0 | 28.6783 | -57.0967 | -32.4786 |
| 0 | 0 | 4124 | 0 | 0 | 5 | 12.49 | 6.63 | 0 | 13.3464 | -74.7168 | -51.5975 |
| | | | | 0 | 12 | 11.22 | 6.33 | 0 | 13.1087 | -66.2626 | -49.2727 |
| | | | | 1 | 52 | 9.63 | 5.9 | 0 | 2.36825 | -73.6404 | -34.3044 |
| | | | | 1 | 54 | 9.6 | 5.89 | 0 | 41.3499 | -54.0578 | -53.0977 |
| | | | | 2 | 84 | 9.24 | 5.78 | 0 | 33.4977 | -43.1069 | -55.118 |
| | | | | 3 | 114 | 8.96 | 5.69 | 0 | 31.2761 | -59.3198 | -58.5425 |
| | | | | 7 | 193 | 8.47 | 5.53 | 0 | 26.3081 | -53.625 | -50.7807 |
| 0 | 0 | 2012 | 0 | 0 | 0.01 | 11.49 | 6.39 | 0 | -42.2299 | -60.9708 | -31.3586 |
| | | | | 0 | 14 | 10.93 | 6.25 | 0 | -24.4743 | -64.0952 | -23.2345 |
| | | | | 0 | 25 | 10.32 | 6.09 | 0 | -32.1019 | -57.6173 | -24.1287 |
| | | | | 3 | 119 | 8.93 | 5.68 | 0 | -22.2405 | -73.166 | -21.0504 |
| | | | | 13 | 328 | 8.07 | 5.39 | 0 | -31.3957 | -47.1821 | -30.7326 |
| 0 | 0 | 335 | 0 | 0 | 11 | 11.31 | 6.35 | 0 | 40.115 | 54.5969 | 15.4274 |
| 0 | 0 | 643 | 0 | 0 | 12 | 11.2 | 6.32 | 0 | -12.3452 | 10.9967 | -0.683603 |
| | | | | 0 | 13 | 11.08 | 6.29 | 0 | -22.8466 | 8.23823 | -0.682769 |

| | | | | | | | | | | | | | |
|---|---|---|---|---|---|---|---|---|---|---|---|---|---|
| | | | | | | 5 | 159 | 8.68 | 5.6 | 0 | -20.8631 | 5.40467 | -12.1166 |
| | | 0 | 0 | 1382 | 0 | 0 | 16 | 10.74 | 6.2 | 0 | -16.1765 | -66.1534 | -46.4321 |
| | | | | | | 0 | 0.02 | 10.53 | 6.15 | 0 | -36.8491 | -57.6239 | -51.0212 |
| | | | | | | 6 | 167 | 8.6 | 5.57 | 0 | -30.5389 | -66.8433 | -56.542 |
| | | | | | | 6 | 181 | 8.52 | 5.54 | 0 | -21.6374 | -72.287 | -51.7521 |
| | | 0 | 0 | 111 | 0 | 0 | 28 | 10.23 | 6.07 | 0 | 27.3725 | 55.5007 | 32.8546 |
| | | 0 | 0 | 271 | 0 | 1 | 42 | 9.81 | 5.95 | 0 | -11.3999 | -15.783 | 4.9888 |
| | | 0 | 0 | 140 | 0 | 2 | 85 | 9.23 | 5.78 | 0 | 42.8641 | 39.8503 | 32.8743 |
| | | 0 | 0 | 948 | 0 | 2 | 85 | 9.22 | 5.77 | 0 | 1.41981 | -55.964 | -15.4981 |
| | | | | | | 5 | 164 | 8.64 | 5.59 | 0 | 1.83226 | -75.9588 | -12.9547 |
| | | | | | | 6 | 171 | 8.57 | 5.56 | 0 | 3.8809 | -64.1671 | -14.1145 |
| | | | | | | 11 | 0.29 | 8.16 | 5.42 | 0 | 4.74571 | -65.0587 | -27.6046 |
| | | | | | | 32 | 665 | 7.54 | 5.19 | 0 | 0.563848 | -54.6958 | -7.03265 |
| | | 0 | 0 | 978 | 0 | 2 | 88 | 9.19 | 5.76 | 0 | -29.5461 | 52.6906 | 19.8961 |
| | | | | | | 3 | 97 | 9.1 | 5.73 | 0 | -36.6791 | 40.4678 | 29.4262 |
| | | | | | | 5 | 164 | 8.63 | 5.58 | 0 | -43.3949 | 43.9376 | 21.8552 |
| | | | | | | 15 | 367 | 7.98 | 5.36 | 0 | -45.5953 | 34.5731 | 31.1117 |
| | | 0 | 0 | 224 | 0 | 3 | 105 | 9.04 | 5.71 | 0 | 20.8848 | 14.2423 | -0.331678 |
| | | 0 | 0 | 191 | 0 | 5 | 164 | 8.63 | 5.58 | 0 | -60.1317 | -68.6248 | -6.42133 |
| | | | | | | 0.01 | 266 | 8.23 | 5.44 | 0 | -57.0069 | -62.9127 | 3.74815 |

LID-RightGo_OFF

| p | c | p(FWE-corr) | p(FDR-corr) | equivk | p(unc) | p(FWE-corr).1 | p(FDR-corr).1 | T | equivZ | p(unc).1 | MNI_x | MNI_y | MNI_z |
|---|---|---|---|---|---|---|---|---|---|---|---|---|---|
| **0** | 15 | 0 | 0 | 15389 | 0 | 0 | 0 | 17.59 | Inf | 0 | 62.2192 | -17.0574 | 29.7272 |
| | | | | | | 0 | 0 | 13.57 | 7.47 | 0 | 61.6962 | -33.7917 | 45.4356 |
| | | | | | | 0 | 0 | 12.05 | 7.09 | 0 | 51.8529 | -26.4197 | 44.724 |
| | | | | | | 0 | 1 | 10.63 | 6.68 | 0 | 68.1037 | -29.2859 | 29.5176 |
| | | | | | | 0 | 1 | 10.22 | 6.55 | 0 | 52.5189 | -16.1112 | 23.7623 |
| | | | | | | 0 | 1 | 10.22 | 6.55 | 0 | 56.2669 | -28.5899 | 55.1351 |
| | | | | | | 0 | 1 | 10.21 | 6.54 | 0 | 51.3403 | -41.3935 | 52.9798 |
| | | | | | | 0 | 1 | 10.17 | 6.53 | 0 | 38.5734 | -32.042 | 40.2955 |
| | | | | | | 0 | 5 | 9.27 | 6.22 | 0 | 46.6514 | -48.6818 | 56.3722 |
| | | | | | | 1 | 19 | 8.57 | 5.96 | 0 | 37.6361 | -37.3973 | 51.7542 |
| | | | | | | 1 | 32 | 8.28 | 5.84 | 0 | 65.2709 | -15.6989 | 19.1847 |
| | | | | | | 2 | 52 | 8.02 | 5.73 | 0 | 38.4311 | -28.9416 | 17.0636 |
| | | | | | | 3 | 83 | 7.77 | 5.63 | 0 | 51.9643 | -30.8316 | 28.7635 |
| | | | | | | 11 | 259 | 7.15 | 5.35 | 0 | 38.2314 | -29.1522 | 54.2082 |
| | | 0 | 0 | 60900 | 0 | 0 | 0 | 15.6 | Inf | 0 | -60.2471 | 5.07527 | 4.01125 |
| | | | | | | 0 | 0 | 15.09 | 7.8 | 0 | -51.3622 | -24.6494 | 41.4637 |
| | | | | | | 0 | 0 | 15.01 | 7.79 | 0 | -52.7179 | -25.6704 | 19.1758 |
| | | | | | | 0 | 0 | 13.97 | 7.57 | 0 | -36.7303 | -26.8368 | 60.559 |
| | | | | | | 0 | 0 | 13.89 | 7.55 | 0 | -58.9186 | -23.6473 | 50.0543 |
| | | | | | | 0 | 0 | 13.24 | 7.39 | 0 | -66.6302 | -23.5559 | 21.1028 |
| | | | | | | 0 | 0 | 12.8 | 7.29 | 0 | -58.082 | -35.0458 | 47.356 |
| | | | | | | 0 | 0 | 12.35 | 7.17 | 0 | -23.7942 | -38.5053 | 69.742 |
| | | | | | | 0 | 0 | 12.21 | 7.13 | 0 | -35.6929 | -0.522113 | 10.1383 |
| | | | | | | 0 | 0 | 12.11 | 7.1 | 0 | -44.2768 | -33.0176 | 65.58 |
| | | | | | | 0 | 0 | 11.97 | 7.07 | 0 | -52.3363 | -24.3123 | 61.2277 |
| | | | | | | 0 | 0 | 11.94 | 7.06 | 0 | -40.7054 | -35.5219 | 40.5724 |
| | | | | | | 0 | 0 | 11.83 | 7.03 | 0 | 2.91187 | 3.47676 | 68.1509 |
| | | | | | | 0 | 0 | 11.61 | 6.97 | 0 | -44.5739 | -21.4708 | 66.7006 |
| | | | | | | 0 | 0 | 11.47 | 6.93 | 0 | -55.9026 | 7.63857 | 24.1047 |
| | | | | | | 0 | 0 | 11.45 | 6.92 | 0 | -42.7473 | -48.9524 | 53.9906 |
| | | | | | | 0 | 0 | 11.1 | 6.82 | 0 | -64.2542 | -22.3534 | 32.4042 |
| | | | | | | 0 | 1 | 10.85 | 6.75 | 0 | -35.143 | -17.8667 | 7.78909 |
| | | | | | | 0 | 1 | 10.79 | 6.73 | 0 | -7.85068 | -4.66936 | 50.8177 |
| | | | | | | 0 | 1 | 10.77 | 6.72 | 0 | -35.367 | -18.9337 | 65.6144 |
| | | | | | | 0 | 1 | 10.75 | 6.71 | 0 | -61.6394 | -31.2776 | 22.3797 |
| | | | | | | 0 | 1 | 10.61 | 6.67 | 0 | -39.4337 | 8.29817 | 3.56093 |
| | | | | | | 0 | 1 | 10.6 | 6.67 | 0 | -2.13217 | 4.55079 | 39.4509 |
| | | | | | | 0 | 1 | 10.59 | 6.66 | 0 | -34.6157 | -21.5507 | 19.2163 |
| | | | | | | 0 | 1 | 10.42 | 6.61 | 0 | -44.3061 | -40.9226 | 57.7399 |
| | | | | | | 0 | 1 | 10.31 | 6.58 | 0 | 0.5467 | -13.2768 | 66.8548 |
| | | | | | | 0 | 1 | 10.24 | 6.55 | 0 | -1.22133 | -11.591 | 53.5757 |
| | | | | | | 0 | 1 | 10.09 | 6.5 | 0 | -39.2371 | -1.1663 | -6.37149 |
| | | | | | | 0 | 2 | 10.05 | 6.49 | 0 | -7.63815 | -25.2202 | 46.864 |
| | | | | | | 0 | 2 | 10.04 | 6.48 | 0 | -59.2428 | -40.8399 | 32.2473 |
| | | | | | | 0 | 2 | 9.87 | 6.43 | 0 | -38.8616 | -45.1846 | 39.8743 |
| | | | | | | 0 | 2 | 9.8 | 6.41 | 0 | -47.6757 | 3.67277 | 8.02629 |
| | | | | | | 0 | 2 | 9.78 | 6.4 | 0 | -35.1609 | -40.3299 | 64.9424 |
| | | | | | | 0 | 3 | 9.63 | 6.35 | 0 | 8.33828 | 16.0117 | 37.26 |
| | | | | | | 0 | 4 | 9.44 | 6.28 | 0 | -42.1082 | -13.6795 | -6.29924 |
| | | | | | | 0 | 4 | 9.38 | 6.26 | 0 | -25.8231 | -49.712 | 68.0322 |

| | | | | | | | | | | | | |
|---|---|---|---|---|---|---|---|---|---|---|---|---|
| | | | | | 0 | 4 | 9.37 | 6.25 | 0 | 5.42077 | -2.86834 | 48.106 |
| | | | | | 0 | 6 | 9.19 | 6.19 | 0 | -33.8419 | 16.1423 | 6.35212 |
| | | | | | 0 | 12 | 8.84 | 6.06 | 0 | -29.7005 | 4.2891 | -5.79289 |
| | | | | | 0 | 12 | 8.83 | 6.06 | 0 | -25.9932 | 2.66328 | 4.11399 |
| | | | | | 0 | 17 | 8.63 | 5.98 | 0 | -11.6266 | -30.3457 | 65.7105 |
| | | | | | 1 | 0.02 | 8.54 | 5.95 | 0 | -26.8439 | -5.70698 | 2.37552 |
| | | | | | 1 | 23 | 8.46 | 5.91 | 0 | -52.3996 | 12.2895 | -0.880209 |
| | | | | | 1 | 24 | 8.44 | 5.91 | 0 | -10.3538 | -37.0953 | 75.471 |
| | | | | | 1 | 32 | 8.29 | 5.84 | 0 | -32.4925 | -14.3426 | 56.0266 |
| | | | | | 1 | 0.04 | 8.16 | 5.79 | 0 | -16.6015 | 12.1251 | -9.53236 |
| | | | | | 1 | 44 | 8.11 | 5.77 | 0 | -9.46112 | -46.1344 | 63.547 |
| | | | | | 1 | 44 | 8.11 | 5.77 | 0 | -31.2793 | -10.7089 | 74.1428 |
| | | | | | 4 | 112 | 7.61 | 5.56 | 0 | -28.3743 | 23.9074 | 8.53058 |
| | | | | | 4 | 115 | 7.59 | 5.55 | 0 | -18.7819 | -13.6655 | 71.297 |
| | | | | | 5 | 131 | 7.52 | 5.52 | 0 | 2.35339 | 21.5107 | 42.8307 |
| | | | | | 7 | 182 | 7.34 | 5.44 | 0 | -45.7067 | 2.92402 | -20.0278 |
| | | | | | 7 | 186 | 7.33 | 5.43 | 0 | -11.3782 | -52.6263 | 75.8646 |
| | | | | | 12 | 286 | 7.1 | 5.33 | 0 | -33.7011 | -50.907 | 65.4861 |
| | | | | | 16 | 364 | 6.98 | 5.27 | 0 | -26.7405 | -4.34289 | 12.1393 |
| | 0 | 0 | 28536 | 0 | 0 | 0 | 13.91 | 7.55 | 0 | 14.3109 | -55.7112 | -20.1377 |
| | | | | | 0 | 0 | 13.55 | 7.47 | 0 | 3.65732 | -55.7184 | -7.7205 |
| | | | | | 0 | 0 | 13.09 | 7.36 | 0 | 2.45554 | -64.6934 | -20.8825 |
| | | | | | 0 | 0 | 12.31 | 7.16 | 0 | 43.9819 | -58.0157 | -27.9151 |
| | | | | | 0 | 0 | 12.2 | 7.13 | 0 | 32.0441 | -54.1959 | -25.7132 |
| | | | | | 0 | 0 | 12.06 | 7.09 | 0 | -38.6204 | -50.6311 | -28.1914 |
| | | | | | 0 | 0 | 11.5 | 6.94 | 0 | 5.56674 | -67.8919 | -35.916 |
| | | | | | 0 | 0 | 11.44 | 6.92 | 0 | 31.9943 | -64.882 | -23.2596 |
| | | | | | 0 | 0 | 11.36 | 6.89 | 0 | 25.5605 | -57.1782 | -53.207 |
| | | | | | 0 | 0 | 11.27 | 6.87 | 0 | 9.06845 | -65.6313 | -54.1986 |
| | | | | | 0 | 1 | 10.93 | 6.77 | 0 | 17.745 | -60.5305 | -45.5823 |
| | | | | | 0 | 1 | 10.62 | 6.67 | 0 | 5.75292 | -72.2667 | -47.765 |
| | | | | | 0 | 1 | 10.46 | 6.62 | 0 | -21.737 | -69.7619 | -22.8547 |
| | | | | | 0 | 1 | 10.36 | 6.59 | 0 | -28.3046 | -53.0228 | -30.7835 |
| | | | | | 0 | 1 | 10.15 | 6.52 | 0 | 17.246 | -76.6963 | -48.076 |
| | | | | | 0 | 2 | 9.87 | 6.43 | 0 | 32.9468 | -46.6043 | -48.8112 |
| | | | | | 0 | 2 | 9.76 | 6.39 | 0 | -40.8545 | -58.7095 | -30.0976 |
| | | | | | 0 | 3 | 9.68 | 6.37 | 0 | -21.1489 | -47.579 | -27.2363 |
| | | | | | 0 | 3 | 9.6 | 6.34 | 0 | 39.8094 | -52.9693 | -52.0107 |
| | | | | | 0 | 6 | 9.19 | 6.19 | 0 | 4.54826 | -46.0864 | -8.14738 |
| | | | | | 1 | 18 | 8.61 | 5.97 | 0 | 5.61525 | -47.3603 | -19.114 |
| | | | | | 1 | 34 | 8.25 | 5.83 | 0 | 10.3958 | -36.463 | -20.3896 |
| | | | | | 1 | 45 | 8.09 | 5.76 | 0 | -33.8873 | -65.5843 | -24.6797 |
| | | | | | 2 | 55 | 7.99 | 5.72 | 0 | -16.2389 | -61.1583 | -16.8443 |
| | | | | | 2 | 69 | 7.87 | 5.67 | 0 | 30.6747 | -36.0359 | -38.6442 |
| | | | | | 2 | 69 | 7.87 | 5.67 | 0 | -16.0372 | -46.8445 | -18.9423 |
| | | | | | 3 | 72 | 7.84 | 5.66 | 0 | 22.3329 | -60.6607 | -31.1451 |
| | | | | | 8 | 195 | 7.3 | 5.42 | 0 | 46.139 | -53.8393 | -43.9144 |
| | | | | | 14 | 318 | 7.05 | 5.3 | 0 | -51.2196 | -61.2343 | -33.3601 |
| | | | | | 14 | 329 | 7.03 | 5.29 | 0 | 51.4532 | -57.9977 | -33.2318 |
| | | | | | 19 | 432 | 6.89 | 5.23 | 0 | 45.4748 | -73.8677 | -28.6211 |
| | | | | | 21 | 477 | 6.84 | 5.2 | 0 | -5.56122 | -48.6468 | -8.75705 |
| | | | | | 29 | 618 | 6.7 | 5.13 | 0 | 26.3402 | -32.0046 | -29.7394 |

| | | | | | | | | | | | | | |
|---|---|---|---|---|---|---|---|---|---|---|---|---|---|
| | | 0 | 0 | 7209 | 0 | 0 | 0 | 12.71 | 7.26 | 0 | 36.3671 | 1.66183 | 8.53143 |
| | | | | | | 0 | 0 | 12.18 | 7.12 | 0 | 53.4714 | 14.6061 | -1.9286 |
| | | | | | | 0 | 0 | 11.45 | 6.92 | 0 | 40.7458 | 5.54712 | -2.77231 |
| | | | | | | 0 | 1 | 10.82 | 6.73 | 0 | 27.6306 | 5.11972 | -0.975902 |
| | | | | | | 0 | 6 | 9.24 | 6.21 | 0 | 37.261 | 14.9632 | 1.91671 |
| | | | | | | 0 | 8 | 9.02 | 6.13 | 0 | 54.8083 | 8.35963 | 10.0038 |
| | | | | | | 1 | 27 | 8.37 | 5.88 | 0 | 55.5654 | 6.33145 | 27.306 |
| | | | | | | 2 | 62 | 7.93 | 5.69 | 0 | 29.66 | 26.3771 | 9.65427 |
| | | | | | | 4 | 116 | 7.59 | 5.55 | 0 | 23.8253 | 7.23246 | -10.3177 |
| | | | | | | 5 | 129 | 7.53 | 5.52 | 0 | 60.0019 | 13.5608 | 20.9769 |
| | | 0 | 0 | 643 | 0 | 0 | 1 | 10.72 | 6.7 | 0 | -13.696 | -14.8899 | 6.22288 |
| | | | | | | 0 | 13 | 8.79 | 6.04 | 0 | -14.9736 | -6.4356 | 16.8922 |
| | | 0 | 0 | 1769 | 0 | 0 | 2 | 9.96 | 6.46 | 0 | -36.9429 | -59.9345 | -52.1346 |
| | | | | | | 0 | 17 | 8.63 | 5.98 | 0 | -20.7206 | -69.176 | -50.7808 |
| | | | | | | 5 | 134 | 7.51 | 5.51 | 0 | -32.2307 | -48.8204 | -48.2434 |
| | | 0 | 0 | 154 | 0 | 0 | 3 | 9.64 | 6.35 | 0 | 0.0806738 | -8.26991 | 12.2023 |
| | | 0 | 0 | 282 | 0 | 0 | 11 | 8.87 | 6.07 | 0 | -59.0649 | -67.4375 | -7.71403 |
| | | 0 | 0 | 754 | 0 | 0 | 17 | 8.63 | 5.98 | 0 | -47.8836 | 40.9982 | 25.3319 |
| | | 0 | 0 | 310 | 0 | 1 | 17 | 8.61 | 5.97 | 0 | -12.7766 | 8.41813 | 5.52498 |
| | | 0 | 0 | 254 | 0 | 1 | 19 | 8.56 | 5.95 | 0 | -14.8079 | -24.6388 | -11.9283 |
| | | 0 | 0 | 197 | 0 | 3 | 84 | 7.76 | 5.62 | 0 | 14.4209 | 1.91296 | 15.0433 |
| | | 0 | 0 | 164 | 0 | 4 | 101 | 7.67 | 5.58 | 0 | 41.5455 | 38.459 | 29.4159 |
| | | 0 | 2 | 111 | 1 | 5 | 121 | 7.56 | 5.54 | 0 | 18.2117 | -22.404 | -11.3177 |
| | | | | | | 26 | 0.57 | 6.75 | 5.16 | 0 | 18.7244 | -12.9305 | -12.5955 |
| | | 0 | 0 | 296 | 0 | 6 | 147 | 7.45 | 5.49 | 0 | 42.4582 | 47.7308 | 6.0071 |

HC-LeftGo_OFF

| p | c | p(FWE-corr) | p(FDR-corr) | equivk | p(unc) | p(FWE-corr).1 | p(FDR-corr).1 | T | equivZ | p(unc).1 | MNI_x | MNI_y | MNI_z |
|---|---|---|---|---|---|---|---|---|---|---|---|---|---|
| **0** | 14 | 0 | 0 | 40475 | 0 | 0 | 0 | 14.57 | 7.35 | 0 | 45.2178 | -31.1876 | 51.8571 |
| | | | | | | 0 | 0 | 14.14 | 7.26 | 0 | 40.0227 | -21.0314 | 53.4546 |
| | | | | | | 0 | 1 | 12.91 | 6.99 | 0 | 50.27 | -17.0038 | 61.1522 |
| | | | | | | 0 | 1 | 12.38 | 6.87 | 0 | 44.6369 | -19.8779 | 22.5689 |
| | | | | | | 0 | 1 | 12.2 | 6.82 | 0 | 41.5504 | -23.8857 | 64.2223 |
| | | | | | | 0 | 1 | 12.09 | 6.8 | 0 | 58.019 | -22.9273 | 45.315 |
| | | | | | | 0 | 1 | 11.88 | 6.74 | 0 | 63.7287 | -30.2673 | 42.3488 |
| | | | | | | 0 | 1 | 11.59 | 6.67 | 0 | 41.1588 | 0.647487 | -1.11676 |
| | | | | | | 0 | 2 | 11.07 | 6.53 | 0 | 39.091 | 2.10827 | 12.3654 |
| | | | | | | 0 | 3 | 10.9 | 6.49 | 0 | 57.4926 | -45.6466 | 53.5908 |
| | | | | | | 0 | 4 | 10.72 | 6.43 | 0 | 67.6863 | -14.65 | 23.6473 |
| | | | | | | 0 | 5 | 10.53 | 6.38 | 0 | 32.4237 | -18.6612 | 10.3587 |
| | | | | | | 0 | 5 | 10.49 | 6.37 | 0 | 49.4341 | -28.0778 | 64.3158 |
| | | | | | | 0 | 6 | 10.38 | 6.33 | 0 | 62.5262 | 14.3999 | 5.19309 |
| | | | | | | 0 | 6 | 10.3 | 6.31 | 0 | 49.9708 | -6.40636 | 58.9789 |
| | | | | | | 0 | 9 | 10.02 | 6.23 | 0 | 33.8761 | -10.307 | 66.8488 |
| | | | | | | 0 | 9 | 9.99 | 6.22 | 0 | 59.9605 | -23.6867 | 19.598 |
| | | | | | | 0 | 9 | 9.96 | 6.21 | 0 | 51.3465 | 11.7455 | 11.3282 |
| | | | | | | 0 | 0.01 | 9.84 | 6.17 | 0 | 64.2303 | -42.3418 | 32.9109 |
| | | | | | | 0 | 13 | 9.7 | 6.13 | 0 | 29.078 | -5.93253 | 4.85383 |
| | | | | | | 0 | 16 | 9.47 | 6.05 | 0 | 20.0048 | 7.30036 | 3.29572 |
| | | | | | | 0 | 21 | 9.21 | 5.97 | 0 | 52.6206 | -17.3564 | 16.3592 |
| | | | | | | 1 | 34 | 8.92 | 5.87 | 0 | 23.4156 | 19.9553 | 4.35987 |
| | | | | | | 1 | 35 | 8.88 | 5.86 | 0 | 34.7542 | -21.1581 | 74.8312 |
| | | | | | | 1 | 45 | 8.69 | 5.79 | 0 | 26.3858 | -56.8098 | 65.3931 |
| | | | | | | 1 | 52 | 8.55 | 5.74 | 0 | 41.2639 | -8.24192 | 3.5605 |
| | | | | | | 2 | 68 | 8.36 | 5.67 | 0 | 29.5565 | -1.12746 | -7.23998 |
| | | | | | | 2 | 71 | 8.31 | 5.65 | 0 | 47.2788 | 10.5648 | 3.49091 |
| | | | | | | 2 | 85 | 8.17 | 5.6 | 0 | 50.1858 | -42.4694 | 35.0015 |
| | | | | | | 3 | 96 | 8.07 | 5.56 | 0 | 64.9928 | 0.180978 | 7.40293 |
| | | | | | | 3 | 105 | 8.01 | 5.54 | 0 | 50.1538 | -0.346667 | 4.77609 |
| | | | | | | 4 | 132 | 7.85 | 5.47 | 0 | 28.4951 | -44.0365 | 68.7295 |
| | | | | | | 5 | 142 | 7.8 | 5.45 | 0 | 60.4134 | -31.021 | 27.4258 |
| | | | | | | 5 | 144 | 7.79 | 5.45 | 0 | 36.6451 | -47.9514 | 70.1574 |
| | | | | | | 5 | 149 | 7.76 | 5.44 | 0 | 65.0129 | -9.58785 | 11.6084 |
| | | | | | | 9 | 225 | 7.5 | 5.33 | 0 | 49.3114 | -38.2159 | 62.2554 |
| | | | | | | 11 | 294 | 7.34 | 5.27 | 0 | 34.8791 | 16.9393 | 7.12594 |
| | | | | | | 11 | 294 | 7.34 | 5.26 | 0 | 23.8933 | 7.14851 | -9.14202 |
| | | | | | | 12 | 295 | 7.33 | 5.26 | 0 | 25.7224 | -41.1394 | 78.4657 |
| | | | | | | 15 | 367 | 7.19 | 5.2 | 0 | 22.1067 | -7.43262 | 79.5268 |
| | | | | | | 16 | 375 | 7.17 | 5.19 | 0 | 9.85288 | 3.92598 | 7.20346 |
| | | | | | | 17 | 0.4 | 7.12 | 5.17 | 0 | 38.0759 | 7.70146 | -14.3025 |
| | | | | | | 24 | 539 | 6.95 | 5.09 | 0 | 30.5586 | -28.4456 | 15.7711 |
| | | | | | | 34 | 707 | 6.77 | 5.02 | 0 | 35.7744 | -42.9338 | 61.8166 |
| | | | | | | 34 | 707 | 6.77 | 5.02 | 0 | 59.2351 | -37.4556 | 55.1686 |
| | | 0 | 0 | 4574 | 0 | 0 | 0 | 14.45 | 7.32 | 0 | -24.9611 | -57.9785 | -54.5116 |
| | | | | | | 8 | 216 | 7.53 | 5.34 | 0 | -0.765009 | -73.8273 | -37.8327 |
| | | 0 | 0 | 7485 | 0 | 0 | 1 | 12.56 | 6.91 | 0 | -38.1263 | -0.578737 | 7.85808 |
| | | | | | | 0 | 15 | 9.52 | 6.07 | 0 | -30.3127 | -8.32622 | 8.74027 |
| | | | | | | 0 | 15 | 9.52 | 6.07 | 0 | -57.4339 | 10.2728 | -1.27915 |

| | | | | | | | | | | | | | |
|---|---|---|---|---|---|---|---|---|---|---|---|---|---|
| | | | | | | 1 | 34 | 8.94 | 5.88 | 0 | -44.001 | -9.96986 | 8.42293 |
| | | | | | | 1 | 35 | 8.87 | 5.85 | 0 | -56.7897 | 6.77587 | 27.7128 |
| | | | | | | 1 | 45 | 8.7 | 5.79 | 0 | -50.5865 | 9.30912 | 6.26652 |
| | | | | | | 1 | 49 | 8.61 | 5.76 | 0 | -49.0196 | 3.29789 | 14.7716 |
| | | | | | | 1 | 56 | 8.49 | 5.72 | 0 | -22.8177 | 7.15301 | -0.700131 |
| | | | | | | 1 | 59 | 8.45 | 5.7 | 0 | -53.8284 | -3.7828 | 2.79648 |
| | | | | | | 2 | 74 | 8.28 | 5.64 | 0 | -40.8804 | -13.3992 | -10.1322 |
| | | | | | | 3 | 101 | 8.03 | 5.55 | 0 | -40.0397 | -14.3398 | -0.317627 |
| | | | | | | 3 | 109 | 7.98 | 5.52 | 0 | -39.0561 | 15.9084 | -3.41476 |
| | | | | | | 23 | 0.52 | 6.97 | 5.1 | 0 | -27.0102 | -2.17123 | -10.5454 |
| | | 0 | 0 | 14133 | 0 | 0 | 1 | 12.4 | 6.87 | 0 | -55.3627 | -24.9894 | 49.1522 |
| | | | | | | 0 | 1 | 11.82 | 6.73 | 0 | -63.2621 | -28.4685 | 47.9932 |
| | | | | | | 0 | 4 | 10.63 | 6.41 | 0 | -59.322 | -21.4374 | 36.2847 |
| | | | | | | 0 | 6 | 10.29 | 6.31 | 0 | -63.4307 | -31.3854 | 31.8897 |
| | | | | | | 0 | 6 | 10.24 | 6.3 | 0 | -64.7307 | -37.7893 | 22.7585 |
| | | | | | | 0 | 8 | 10.08 | 6.25 | 0 | -55.6211 | -22.4945 | 17.4381 |
| | | | | | | 0 | 15 | 9.57 | 6.09 | 0 | -46.7524 | -19.8182 | 23.0503 |
| | | | | | | 0 | 15 | 9.55 | 6.08 | 0 | -43.3897 | -36.2195 | 47.5996 |
| | | | | | | 0 | 15 | 9.55 | 6.08 | 0 | -57.2424 | -48.4128 | 29.1365 |
| | | | | | | 0 | 0.02 | 9.26 | 5.99 | 0 | -50.3361 | -34.3814 | 37.4464 |
| | | | | | | 1 | 34 | 8.93 | 5.87 | 0 | -55.528 | -41.1236 | 58.7854 |
| | | | | | | 1 | 34 | 8.91 | 5.87 | 0 | -38.5687 | -44.1608 | 45.1036 |
| | | | | | | 2 | 81 | 8.2 | 5.61 | 0 | -43.1425 | -49.3495 | 61.5187 |
| | | | | | | 4 | 124 | 7.89 | 5.49 | 0 | -54.5552 | -31.981 | 60.2814 |
| | | | | | | 18 | 417 | 7.1 | 5.16 | 0 | -55.6349 | -52.0398 | 55.0018 |
| | | | | | | 46 | 0.93 | 6.6 | 4.94 | 0 | -41.1406 | -42.8534 | 69.3713 |
| | | 0 | 0 | 983 | 0 | 0 | 1 | 11.82 | 6.73 | 0 | 17.0062 | -17.0099 | 5.80007 |
| | | | | | | 22 | 495 | 7 | 5.12 | 0 | 17.8351 | -10.1294 | -3.63096 |
| | | 0 | 0 | 4597 | 0 | 0 | 16 | 9.44 | 6.05 | 0 | 4.59017 | 1.44024 | 53.0822 |
| | | | | | | 1 | 0.05 | 8.59 | 5.75 | 0 | 2.64726 | -1.03531 | 71.7304 |
| | | | | | | 1 | 56 | 8.49 | 5.72 | 0 | 3.83015 | 2.96944 | 40.1823 |
| | | | | | | 5 | 153 | 7.74 | 5.43 | 0 | 11.3764 | 2.91711 | 73.9971 |
| | | | | | | 6 | 156 | 7.73 | 5.42 | 0 | 6.92379 | -17.0204 | 47.3161 |
| | | | | | | 17 | 389 | 7.14 | 5.18 | 0 | -5.85328 | 11.0291 | 42.048 |
| | | | | | | 34 | 707 | 6.77 | 5.02 | 0 | 2.72702 | 13.8713 | 37.8415 |
| | | 0 | 0 | 2091 | 0 | 1 | 46 | 8.67 | 5.78 | 0 | -19.9942 | -51.3527 | -22.5323 |
| | | | | | | 2 | 71 | 8.31 | 5.65 | 0 | -34.5512 | -38.2202 | -29.3533 |
| | | | | | | 4 | 129 | 7.86 | 5.48 | 0 | -32.5787 | -50.8397 | -28.2577 |
| | | 0 | 11 | 103 | 5 | 1 | 49 | 8.62 | 5.76 | 0 | -20.24 | -43.3203 | -53.0845 |
| | | 0 | 0 | 998 | 0 | 1 | 0.05 | 8.59 | 5.75 | 0 | 22.5792 | -62.0348 | -53.2634 |
| | | | | | | 5 | 149 | 7.76 | 5.44 | 0 | 12.6292 | -61.2594 | -53.186 |
| | | | | | | 9 | 238 | 7.46 | 5.32 | 0 | 14.699 | -70.2805 | -47.9025 |
| | | 0 | 7 | 122 | 3 | 1 | 0.05 | 8.58 | 5.75 | 0 | -33.1916 | -39.8152 | -47.0235 |
| | | 0 | 2 | 173 | 1 | 2 | 69 | 8.34 | 5.66 | 0 | -19.5438 | -0.120583 | 12.1732 |
| | | 0 | 0 | 255 | 0 | 2 | 81 | 8.2 | 5.61 | 0 | 28.1591 | -48.1 | -50.8622 |
| | | | | | | 21 | 468 | 7.03 | 5.13 | 0 | 28.5936 | -39.1086 | -51.692 |
| | | 0 | 3 | 150 | 1 | 3 | 93 | 8.09 | 5.57 | 0 | -54.9451 | -65.0291 | 7.90042 |
| | | 0 | 0 | 251 | 0 | 4 | 134 | 7.84 | 5.47 | 0 | -30.7768 | 18.9238 | 8.90506 |

PDnoLID-LeftGo_OFF

| p | c | p(FWE-corr) | p(FDR-corr) | equivk | p(unc) | p(FWE-corr).1 | p(FDR-corr).1 | T | equivZ | p(unc).1 | MNI_x | MNI_y | MNI_z |
|---|---|---|---|---|---|---|---|---|---|---|---|---|---|
| **0** | 15 | 0 | 0 | 22163 | 0 | 0 | 0 | 18.4 | 7.65 | 0 | 61.6583 | -16.653 | 24.3155 |
| | | | | | | 0 | 0 | 17.38 | 7.5 | 0 | 51.1217 | -17.2098 | 54.6689 |
| | | | | | | 0 | 1 | 14.95 | 7.11 | 0 | 46.8415 | -18.6616 | 19.2685 |
| | | | | | | 0 | 1 | 14.7 | 7.07 | 0 | 37.6166 | -21.0644 | 66.8537 |
| | | | | | | 0 | 1 | 13.73 | 6.88 | 0 | 57.39 | -15.6554 | 38.0963 |
| | | | | | | 0 | 2 | 13.07 | 6.75 | 0 | 55.5893 | -34.1084 | 28.5858 |
| | | | | | | 0 | 3 | 12.68 | 6.67 | 0 | 40.507 | -30.7379 | 53.2035 |
| | | | | | | 0 | 3 | 12.61 | 6.65 | 0 | 33.1978 | -28.0536 | 57.8156 |
| | | | | | | 0 | 4 | 12.14 | 6.55 | 0 | 54.0785 | -28.5214 | 50.7402 |
| | | | | | | 0 | 5 | 12 | 6.51 | 0 | 46.3338 | -27.1222 | 62.4243 |
| | | | | | | 0 | 9 | 11.26 | 6.34 | 0 | 59.3388 | -8.75362 | 15.6079 |
| | | | | | | 0 | 17 | 10.78 | 6.22 | 0 | 31.6459 | -29.8371 | 45.8695 |
| | | | | | | 0 | 18 | 10.62 | 6.17 | 0 | 43.2514 | -23.354 | 46.2445 |
| | | | | | | 0 | 25 | 10.28 | 6.08 | 0 | 59.421 | -38.1962 | 42.9893 |
| | | | | | | 1 | 26 | 10.2 | 6.06 | 0 | 34.8108 | -38.1176 | 47.4082 |
| | | | | | | 1 | 37 | 9.88 | 5.97 | 0 | 27.7674 | -19.4148 | 64.8421 |
| | | | | | | 1 | 38 | 9.85 | 5.96 | 0 | 44.3357 | -12.9346 | 25.4998 |
| | | | | | | 1 | 56 | 9.51 | 5.86 | 0 | 21.0899 | -49.7581 | 73.3666 |
| | | | | | | 2 | 81 | 9.21 | 5.77 | 0 | 43.2163 | -27.8895 | 23.53 |
| | | | | | | 3 | 97 | 9.03 | 5.71 | 0 | 33.271 | -39.2364 | 62.1097 |
| | | | | | | 6 | 166 | 8.59 | 5.57 | 0 | 38.8249 | -10.0887 | 69.5302 |
| | | | | | | 17 | 401 | 7.93 | 5.33 | 0 | 62.4821 | -43.9388 | 28.7825 |
| | | | | | | 19 | 432 | 7.87 | 5.31 | 0 | 24.5664 | -15.4624 | 75.4967 |
| | | 0 | 0 | 4778 | 0 | 0 | 0 | 16.58 | 7.38 | 0 | -28.1866 | -53.4445 | -55.5296 |
| | | | | | | 0 | 1 | 14.7 | 7.07 | 0 | -18.9112 | -62.6109 | -52.4695 |
| | | | | | | 0 | 4 | 12.21 | 6.56 | 0 | -10.0804 | -66.0642 | -41.5553 |
| | | | | | | 0 | 19 | 10.56 | 6.16 | 0 | -28.6815 | -64.9106 | -51.0435 |
| | | | | | | 1 | 32 | 10.03 | 6.01 | 0 | -2.11763 | -70.401 | -43.8503 |
| | | | | | | 11 | 278 | 8.21 | 5.44 | 0 | -1.02179 | -69.1622 | -34.3104 |
| | | 0 | 0 | 6895 | 0 | 0 | 1 | 15.44 | 7.2 | 0 | -7.08696 | -58.2001 | -11.4587 |
| | | | | | | 0 | 1 | 14.17 | 6.97 | 0 | -22.3433 | -51.24 | -24.9323 |
| | | | | | | 0 | 2 | 13.33 | 6.8 | 0 | -31.7921 | -46.3228 | -34.2263 |
| | | | | | | 0 | 9 | 11.25 | 6.33 | 0 | -32.1199 | -57.4305 | -25.1776 |
| | | | | | | 0 | 17 | 10.8 | 6.22 | 0 | -1.01639 | -66.8692 | -19.7306 |
| | | | | | | 1 | 38 | 9.84 | 5.96 | 0 | -25.5469 | -64.0959 | -23.1288 |
| | | | | | | 2 | 85 | 9.16 | 5.75 | 0 | 1.75031 | -74.8283 | -12.7717 |
| | | | | | | 3 | 0.09 | 9.11 | 5.74 | 0 | -42.3121 | -59.7555 | -32.5039 |
| | | 0 | 0 | 10418 | 0 | 0 | 1 | 15.36 | 7.18 | 0 | -46.6437 | -32.1295 | 43.1388 |
| | | | | | | 0 | 2 | 13.45 | 6.83 | 0 | -60.8788 | -24.6456 | 33.5487 |
| | | | | | | 0 | 4 | 12.26 | 6.57 | 0 | -53.7621 | -27.5181 | 20.0639 |
| | | | | | | 0 | 4 | 12.16 | 6.55 | 0 | -62.8725 | -20.0608 | 24.5612 |
| | | | | | | 0 | 6 | 11.76 | 6.46 | 0 | -65.2472 | -48.2045 | 32.5692 |
| | | | | | | 0 | 7 | 11.59 | 6.42 | 0 | -60.366 | -27.7352 | 45.6303 |
| | | | | | | 0 | 25 | 10.26 | 6.08 | 0 | -38.8284 | -44.9689 | 45.121 |
| | | | | | | 1 | 34 | 9.97 | 6 | 0 | -53.4732 | -35.1438 | 59.4301 |
| | | | | | | 1 | 51 | 9.62 | 5.89 | 0 | -63.0591 | -34.7617 | 27.5015 |
| | | | | | | 1 | 53 | 9.58 | 5.88 | 0 | -59.3391 | -50.4377 | 41.7497 |
| | | | | | | 3 | 97 | 9.03 | 5.71 | 0 | -62.0948 | -40.9476 | 38.7096 |
| | | | | | | 4 | 0.13 | 8.78 | 5.63 | 0 | -43.611 | -51.8324 | 56.2095 |

| | | | | | | | | | | | | |
|---|---|---|---|---|---|---|---|---|---|---|---|---|
| | | | | | 5 | 137 | 8.75 | 5.62 | 0 | -45.7922 | -41.986 | 56.873 |
| | | | | | 8 | 207 | 8.43 | 5.51 | 0 | -55.0162 | -45.7 | 51.9794 |
| | | | | | 15 | 0.37 | 8 | 5.36 | 0 | -55.8741 | -51.7579 | 31.1064 |
| | 0 | 0 | 6514 | 0 | 0 | 1 | 13.99 | 6.93 | 0 | 46.1555 | 0.248513 | 10.9054 |
| | | | | | 0 | 4 | 12.34 | 6.59 | 0 | 45.826 | 16.1804 | 0.679822 |
| | | | | | 0 | 4 | 12.16 | 6.55 | 0 | 37.979 | 0.904808 | 17.0722 |
| | | | | | 0 | 4 | 12.04 | 6.52 | 0 | 60.7755 | 11.8643 | 32.4495 |
| | | | | | 0 | 25 | 10.31 | 6.09 | 0 | 34.8559 | 19.196 | 5.8635 |
| | | | | | 0 | 26 | 10.24 | 6.07 | 0 | 38.7087 | -5.83156 | -2.28085 |
| | | | | | 1 | 27 | 10.17 | 6.05 | 0 | 53.6172 | 8.11877 | 8.83431 |
| | | | | | 1 | 58 | 9.48 | 5.85 | 0 | 28.396 | -0.150348 | -8.32906 |
| | | | | | 2 | 69 | 9.34 | 5.81 | 0 | 34.5402 | -16.322 | 10.5628 |
| | | | | | 2 | 85 | 9.16 | 5.75 | 0 | 38.324 | 2.74471 | -10.7012 |
| | | | | | 3 | 95 | 9.06 | 5.72 | 0 | 60.5646 | 6.01623 | 2.70317 |
| | | | | | 12 | 314 | 8.13 | 5.41 | 0 | 35.5231 | -16.7683 | -4.88736 |
| | 0 | 0 | 5181 | 0 | 0 | 1 | 13.59 | 6.86 | 0 | -40.5954 | 3.25382 | 0.478001 |
| | | | | | 0 | 2 | 13.25 | 6.79 | 0 | -55.0385 | 6.05656 | 20.471 |
| | | | | | 0 | 7 | 11.58 | 6.42 | 0 | -45.5817 | -2.91786 | 10.7417 |
| | | | | | 0 | 17 | 10.75 | 6.21 | 0 | -36.9237 | 1.69088 | 11.3555 |
| | | | | | 0 | 18 | 10.66 | 6.19 | 0 | -28.2843 | 22.8315 | 8.56115 |
| | | | | | 0 | 18 | 10.63 | 6.18 | 0 | -41.3581 | 15.2401 | 4.2647 |
| | | | | | 0 | 0.02 | 10.51 | 6.14 | 0 | -40.2359 | -10.1593 | -1.62564 |
| | | | | | 0 | 25 | 10.28 | 6.08 | 0 | -58.355 | 11.2535 | -1.3048 |
| | | | | | 1 | 54 | 9.56 | 5.87 | 0 | -61.1441 | 11.3049 | 12.7184 |
| | | | | | 3 | 97 | 9.02 | 5.71 | 0 | -57.6543 | 6.34009 | 36.4874 |
| | | | | | 0.03 | 637 | 7.58 | 5.21 | 0 | -62.7538 | 8.52838 | 25.5324 |
| | 0 | 0 | 7398 | 0 | 0 | 2 | 12.92 | 6.72 | 0 | 5.83936 | -8.31207 | 42.3365 |
| | | | | | 0 | 3 | 12.55 | 6.64 | 0 | 1.2917 | 22.4335 | 34.1192 |
| | | | | | 0 | 7 | 11.5 | 6.4 | 0 | 0.514962 | -0.460949 | 39.4721 |
| | | | | | 0 | 9 | 11.28 | 6.34 | 0 | 2.70482 | -5.39503 | 62.2855 |
| | | | | | 0 | 17 | 10.74 | 6.21 | 0 | 3.56885 | -25.522 | 48.4657 |
| | | | | | 0 | 17 | 10.73 | 6.2 | 0 | 6.18986 | 20.7002 | 42.7186 |
| | | | | | 0 | 18 | 10.69 | 6.19 | 0 | 3.79555 | 10.695 | 44.0256 |
| | | | | | 0 | 22 | 10.44 | 6.12 | 0 | -11.1746 | 13.6785 | 34.9805 |
| | | | | | 0 | 25 | 10.26 | 6.08 | 0 | 10.6158 | -7.49316 | 53.3326 |
| | | | | | 1 | 36 | 9.91 | 5.98 | 0 | -0.221852 | -1.42815 | 50.1485 |
| | | | | | 1 | 48 | 9.67 | 5.91 | 0 | 7.58936 | 5.49681 | 54.6411 |
| | | | | | 3 | 109 | 8.92 | 5.67 | 0 | -8.15436 | 21.5197 | 28.4752 |
| | | | | | 7 | 195 | 8.47 | 5.53 | 0 | 2.98226 | 5.43796 | 67.0535 |
| | | | | | 0.01 | 255 | 8.27 | 5.46 | 0 | 12.3278 | -29.1425 | 43.2667 |
| | | | | | 15 | 377 | 7.99 | 5.36 | 0 | -7.69892 | -26.2354 | 45.482 |
| | 0 | 0 | 296 | 0 | 0 | 4 | 12.1 | 6.54 | 0 | 34.5793 | -43.9923 | -55.2677 |
| | 0 | 0 | 466 | 0 | 0 | 18 | 10.68 | 6.19 | 0 | 13.2365 | -73.4906 | -52.8227 |
| | | | | | 3 | 94 | 9.07 | 5.73 | 0 | 23.7424 | -77.8202 | -54.1627 |
| | | | | | 18 | 417 | 7.89 | 5.32 | 0 | 18.3589 | -72.5536 | -43.5732 |
| | 0 | 0 | 179 | 0 | 0 | 22 | 10.43 | 6.12 | 0 | 9.90001 | 2.73774 | 9.58028 |
| | 0 | 0 | 794 | 0 | 1 | 29 | 10.12 | 6.04 | 0 | 32.3073 | -58.1461 | -56.4334 |
| | | | | | 2 | 87 | 9.14 | 5.75 | 0 | 39.1945 | -54.1813 | -49.3779 |
| | | | | | 5 | 144 | 8.7 | 5.61 | 0 | 36.4931 | -66.8329 | -57.2007 |
| | | | | | 19 | 435 | 7.85 | 5.31 | 0 | 27.541 | -66.4563 | -57.0493 |
| | 0 | 0 | 107 | 0 | 1 | 53 | 9.58 | 5.88 | 0 | 7.79345 | -53.7001 | 77.2657 |

| | | | | | | | | | | | | | |
|---|---|---|---|---|---|---|---|---|---|---|---|---|---|
| | | 0 | 0 | 133 | 0 | 1 | 57 | 9.49 | 5.85 | 0 | 40.2611 | 42.8012 | 5.36337 |
| | | 0 | 0 | 202 | 0 | 2 | 0.06 | 9.44 | 5.84 | 0 | 47.6337 | -65.9053 | -25.6775 |
| | | 0 | 0 | 194 | 0 | 3 | 109 | 8.92 | 5.68 | 0 | 19.6461 | 14.1559 | -0.323971 |

LID-LeftGo_OFF

| p | c | p(FWE-corr) | p(FDR-corr) | equivk | p(unc) | p(FWE-corr).1 | p(FDR-corr).1 | T | equivZ | p(unc).1 | MNI_x | MNI_y | MNI_z |
|---|---|---|---|---|---|---|---|---|---|---|---|---|---|
| **0** | 22 | 0 | 0 | 79237 | 0 | 0 | 0 | 20.93 | Inf | 0 | 39.3935 | -29.2857 | 56.674 |
| | | | | | | 0 | 0 | 15.76 | Inf | 0 | 36.4457 | 0.705755 | 8.56248 |
| | | | | | | 0 | 0 | 14.55 | 7.69 | 0 | 50.587 | -32.0501 | 55.3426 |
| | | | | | | 0 | 0 | 14.42 | 7.66 | 0 | 44.6327 | -21.0739 | 48.79 |
| | | | | | | 0 | 0 | 13.99 | 7.57 | 0 | 37.4275 | -32.7925 | 41.9756 |
| | | | | | | 0 | 0 | 13.97 | 7.57 | 0 | 52.1572 | -28.6849 | 44.3676 |
| | | | | | | 0 | 0 | 13.58 | 7.48 | 0 | 64.6007 | -19.8711 | 28.5268 |
| | | | | | | 0 | 0 | 13.19 | 7.38 | 0 | 39.1853 | -31.5857 | 65.3974 |
| | | | | | | 0 | 0 | 12.99 | 7.33 | 0 | 46.3433 | -23.9684 | 62.5171 |
| | | | | | | 0 | 0 | 12.87 | 7.3 | 0 | 48.1425 | -16.6174 | 24.8873 |
| | | | | | | 0 | 0 | 12.51 | 7.21 | 0 | -35.8099 | -0.401274 | 7.79635 |
| | | | | | | 0 | 0 | 12.41 | 7.19 | 0 | 51.4506 | 13.2347 | -0.440957 |
| | | | | | | 0 | 0 | 12.41 | 7.19 | 0 | 54.9221 | -15.0123 | 15.7662 |
| | | | | | | 0 | 0 | 12.14 | 7.11 | 0 | -52.4536 | -22.2777 | 44.075 |
| | | | | | | 0 | 0 | 12 | 7.08 | 0 | 34.9401 | 13.8576 | 0.943144 |
| | | | | | | 0 | 0 | 11.79 | 7.02 | 0 | -67.6573 | -23.4801 | 23.4275 |
| | | | | | | 0 | 0 | 11.72 | 7 | 0 | -61.6646 | 7.29434 | 2.99564 |
| | | | | | | 0 | 0 | 11.56 | 6.95 | 0 | 59.7605 | 14.7424 | -1.64961 |
| | | | | | | 0 | 0 | 11.51 | 6.94 | 0 | 60.4396 | -32.7488 | 45.6754 |
| | | | | | | 0 | 0 | 11.35 | 6.89 | 0 | -56.3639 | -25.8499 | 16.8232 |
| | | | | | | 0 | 0 | 11.31 | 6.88 | 0 | 53.711 | -16.0344 | 42.5492 |
| | | | | | | 0 | 0 | 11.29 | 6.88 | 0 | -64.2542 | -22.3534 | 32.4042 |
| | | | | | | 0 | 1 | 10.89 | 6.75 | 0 | 42.05 | 4.5239 | -2.77843 |
| | | | | | | 0 | 1 | 10.82 | 6.73 | 0 | -56.0648 | 7.30517 | 20.5844 |
| | | | | | | 0 | 1 | 10.75 | 6.71 | 0 | 52.9814 | -24.0607 | 19.6805 |
| | | | | | | 0 | 1 | 10.68 | 6.69 | 0 | 29.674 | -51.6469 | 67.9837 |
| | | | | | | 0 | 2 | 10.28 | 6.56 | 0 | -55.1485 | -35.8105 | 51.4414 |
| | | | | | | 0 | 2 | 10.25 | 6.55 | 0 | 46.1237 | -37.9264 | 67.3782 |
| | | | | | | 0 | 2 | 10.2 | 6.54 | 0 | 6.92477 | 4.35102 | 1.19574 |
| | | | | | | 0 | 2 | 10.11 | 6.51 | 0 | 54.8378 | 7.06526 | 8.81386 |
| | | | | | | 0 | 2 | 10.06 | 6.49 | 0 | 32.9324 | -19.2903 | 70.3502 |
| | | | | | | 0 | 3 | 9.99 | 6.47 | 0 | 15.3436 | 3.99391 | 14.988 |
| | | | | | | 0 | 3 | 9.98 | 6.47 | 0 | -62.383 | -33.6555 | 25.4673 |
| | | | | | | 0 | 3 | 9.95 | 6.45 | 0 | 51.9539 | -28.3474 | 30.1718 |
| | | | | | | 0 | 3 | 9.94 | 6.45 | 0 | -38.1951 | -0.14487 | -4.01148 |
| | | | | | | 0 | 3 | 9.91 | 6.44 | 0 | 36.7976 | -15.2355 | 12.8005 |
| | | | | | | 0 | 3 | 9.8 | 6.41 | 0 | -33.7955 | 16.0383 | 7.52629 |
| | | | | | | 0 | 4 | 9.75 | 6.39 | 0 | 54.7922 | -13.0485 | 53.0294 |
| | | | | | | 0 | 4 | 9.68 | 6.36 | 0 | -60.5247 | -31.8489 | 39.2033 |
| | | | | | | 0 | 4 | 9.63 | 6.35 | 0 | 59.8164 | 9.77853 | 30.3344 |
| | | | | | | 0 | 5 | 9.57 | 6.33 | 0 | 43.5058 | -12.9064 | 66.8077 |
| | | | | | | 0 | 6 | 9.4 | 6.27 | 0 | 21.9371 | -56.644 | 65.3156 |
| | | | | | | 0 | 7 | 9.27 | 6.22 | 0 | -46.6621 | -41.7568 | 56.1297 |
| | | | | | | 0 | 8 | 9.24 | 6.21 | 0 | -37.2222 | -23.8658 | 18.5395 |
| | | | | | | 0 | 9 | 9.17 | 6.18 | 0 | 0.0341527 | -8.01524 | 9.78421 |
| | | | | | | 0 | 9 | 9.17 | 6.18 | 0 | -40.905 | -35.8024 | 43.1147 |
| | | | | | | 0 | 15 | 8.83 | 6.06 | 0 | -39.7227 | -12.3894 | -8.82114 |
| | | | | | | 0 | 16 | 8.77 | 6.03 | 0 | -47.8891 | -1.46047 | 7.3114 |
| | | | | | | 0 | 16 | 8.77 | 6.03 | 0 | 39.8713 | -10.906 | 59.1995 |
| | | | | | | 1 | 24 | 8.56 | 5.95 | 0 | 25.823 | -14.825 | 72.1027 |

| | | | | | | | | | | | | |
|---|---|---|---|---|---|---|---|---|---|---|---|---|
| | | | | | 1 | 24 | 8.54 | 5.94 | 0 | 44.5157 | -34.933 | 23.1165 |
| | | | | | 1 | 26 | 8.5 | 5.93 | 0 | -12.7397 | 19.7696 | 5.85895 |
| | | | | | 1 | 31 | 8.39 | 5.88 | 0 | -45.5103 | -20.9387 | 15.5712 |
| | | | | | 1 | 33 | 8.36 | 5.87 | 0 | -54.2386 | -40.9365 | 41.5829 |
| | | | | | 1 | 36 | 8.31 | 5.85 | 0 | -6.84014 | -0.826848 | 6.44045 |
| | | | | | 1 | 37 | 8.3 | 5.85 | 0 | 33.677 | -18.4791 | 2.30116 |
| | | | | | 1 | 48 | 8.16 | 5.79 | 0 | 12.3416 | 14.7927 | 3.60091 |
| | | | | | 1 | 49 | 8.15 | 5.79 | 0 | -56.6604 | 4.5072 | 32.6132 |
| | | | | | 1 | 0.05 | 8.12 | 5.78 | 0 | -14.9297 | -5.44534 | 16.8953 |
| | | | | | 2 | 55 | 8.07 | 5.75 | 0 | -13.8056 | 12.0564 | -3.15221 |
| | | | | | 2 | 58 | 8.03 | 5.74 | 0 | 16.817 | -33.3936 | 65.3313 |
| | | | | | 2 | 68 | 7.94 | 5.7 | 0 | -14.3832 | -12.0312 | 10.3521 |
| | | | | | 2 | 0.07 | 7.92 | 5.69 | 0 | -9.72188 | 5.51841 | 2.90165 |
| | | | | | 2 | 0.07 | 7.92 | 5.69 | 0 | 16.4262 | -59.7554 | 71.8682 |
| | | | | | 3 | 82 | 7.83 | 5.65 | 0 | 33.0037 | 2.37906 | -8.45882 |
| | | | | | 3 | 85 | 7.81 | 5.64 | 0 | 19.3401 | 7.16522 | -14.0015 |
| | | | | | 3 | 85 | 7.81 | 5.64 | 0 | 49.3197 | -44.6175 | 52.5738 |
| | | | | | 4 | 125 | 7.6 | 5.55 | 0 | -42.4622 | -49.513 | 55.4809 |
| | | | | | 5 | 138 | 7.53 | 5.52 | 0 | -40.9835 | -47.8052 | 38.9358 |
| | | | | | 5 | 138 | 7.52 | 5.52 | 0 | -27.2441 | 3.80675 | 5.44141 |
| | | | | | 5 | 138 | 7.52 | 5.52 | 0 | 15.7093 | 21.898 | -5.10203 |
| | | | | | 5 | 149 | 7.48 | 5.5 | 0 | -50.2594 | 11.2214 | -0.778069 |
| | | | | | 6 | 157 | 7.45 | 5.48 | 0 | 39.3196 | 7.73504 | -13.2869 |
| | | | | | 6 | 174 | 7.39 | 5.46 | 0 | 13.1198 | -3.77214 | 12.8097 |
| | | | | | 7 | 0.19 | 7.35 | 5.44 | 0 | -36.6881 | -15.5193 | 1.93025 |
| | | | | | 9 | 224 | 7.26 | 5.4 | 0 | -19.1183 | 2.03243 | 0.548815 |
| | | | | | 9 | 239 | 7.23 | 5.39 | 0 | 25.0937 | 10.3295 | -5.63122 |
| | | | | | 9 | 241 | 7.22 | 5.38 | 0 | 27.7023 | -24.2375 | 76.2557 |
| | | | | | 11 | 277 | 7.13 | 5.34 | 0 | -29.6802 | 3.22421 | -4.54352 |
| | | | | | 11 | 278 | 7.12 | 5.34 | 0 | 10.9812 | -51.9204 | 72.9633 |
| | | | | | 14 | 325 | 7.03 | 5.29 | 0 | -23.5117 | 16.4418 | -4.48253 |
| | | | | | 14 | 333 | 7.02 | 5.29 | 0 | 52.685 | -4.84433 | 9.77768 |
| | | | | | 16 | 359 | 6.98 | 5.27 | 0 | 41.8175 | -7.46651 | -9.73644 |
| | | | | | 0.03 | 0.63 | 6.69 | 5.13 | 0 | -42.7876 | -44.4192 | 63.5173 |
| | 0 | 0 | 14293 | 0 | 0 | 0 | 12.68 | 7.25 | 0 | -20.891 | -53.922 | -20.1149 |
| | | | | | 0 | 0 | 12.34 | 7.17 | 0 | -31.5354 | -51.6051 | -25.4227 |
| | | | | | 0 | 0 | 11.54 | 6.95 | 0 | -1.23642 | -60.8675 | -21.729 |
| | | | | | 0 | 0 | 11.34 | 6.89 | 0 | -19.8302 | -68.4166 | -21.0704 |
| | | | | | 0 | 1 | 10.63 | 6.68 | 0 | -6.73274 | -63.5115 | -11.8972 |
| | | | | | 0 | 5 | 9.51 | 6.31 | 0 | -27.686 | -63.0282 | -23.0942 |
| | | | | | 0 | 5 | 9.49 | 6.3 | 0 | -40.5341 | -60.1986 | -25.2374 |
| | | | | | 0 | 7 | 9.26 | 6.22 | 0 | -36.3581 | -51.9465 | -31.5119 |
| | | | | | 0 | 12 | 8.94 | 6.1 | 0 | -4.836 | -47.9597 | -15.4168 |
| | | | | | 1 | 24 | 8.56 | 5.95 | 0 | -30.8565 | -60.108 | -34.2257 |
| | | | | | 1 | 36 | 8.31 | 5.85 | 0 | -52.3003 | -62.5216 | -32.2567 |
| | | | | | 3 | 85 | 7.81 | 5.64 | 0 | 9.4875 | -43.1731 | -19.7051 |
| | | | | | 4 | 0.12 | 7.62 | 5.56 | 0 | -0.616387 | -53.1531 | -0.754887 |
| | | | | | 0.01 | 249 | 7.2 | 5.37 | 0 | 18.7199 | -34.7687 | -22.729 |
| | | | | | 12 | 285 | 7.11 | 5.33 | 0 | -47.2689 | -51.4464 | -21.19 |
| | | | | | 0.03 | 635 | 6.68 | 5.12 | 0 | 10.8986 | -57.0415 | -11.5242 |
| | | | | | 44 | 888 | 6.5 | 5.03 | 0 | -33.3399 | -70.5298 | -25.7468 |
| | 0 | 0 | 12415 | 0 | 0 | 0 | 12.5 | 7.21 | 0 | 1.71441 | -10.3326 | 51.0895 |

| | | | | | | | | | | | | | |
|---|---|---|---|---|---|---|---|---|---|---|---|---|---|
| | | | | | | 0 | 0 | 11.96 | 7.07 | 0 | 6.11126 | 16.7928 | 34.7497 |
| | | | | | | 0 | 0 | 11.72 | 7 | 0 | 2.79529 | 1.76791 | 39.8744 |
| | | | | | | 0 | 0 | 11.52 | 6.94 | 0 | 2.91187 | 3.47676 | 68.1509 |
| | | | | | | 0 | 3 | 10.03 | 6.48 | 0 | -6.41313 | -3.64966 | 40.6004 |
| | | | | | | 0 | 4 | 9.69 | 6.37 | 0 | 6.32786 | -2.74482 | 49.7436 |
| | | | | | | 0 | 5 | 9.49 | 6.3 | 0 | 10.5676 | -15.7811 | 42.2813 |
| | | | | | | 0 | 5 | 9.49 | 6.3 | 0 | 9.78914 | 8.00842 | 36.1598 |
| | | | | | | 0 | 11 | 9.01 | 6.12 | 0 | 1.42877 | -16.8848 | 66.6198 |
| | | | | | | 1 | 29 | 8.44 | 5.91 | 0 | -3.84236 | 18.445 | 28.7093 |
| | | | | | | 2 | 55 | 8.07 | 5.76 | 0 | -4.93852 | -12.554 | 35.4154 |
| | | | | | | 2 | 59 | 8.02 | 5.73 | 0 | 6.25461 | -37.6507 | 57.4841 |
| | | | | | | 2 | 73 | 7.9 | 5.68 | 0 | -3.57916 | -8.26487 | 69.0717 |
| | | | | | | 2 | 78 | 7.86 | 5.67 | 0 | 19.4037 | 8.91579 | 67.0227 |
| | | | | | | 2 | 79 | 7.85 | 5.66 | 0 | 11.8911 | -23.941 | 50.2229 |
| | | | | | | 3 | 105 | 7.69 | 5.59 | 0 | -9.25636 | -5.74921 | 51.7332 |
| | | | | | | 13 | 301 | 7.08 | 5.32 | 0 | 2.74826 | 12.3284 | 66.9535 |
| | | | | | | 17 | 382 | 6.94 | 5.25 | 0 | 4.78499 | -15.0285 | 75.5627 |
| | | | | | | 23 | 498 | 6.8 | 5.18 | 0 | -15.2943 | -9.68432 | 60.6071 |
| | | | | | | 27 | 0.57 | 6.74 | 5.15 | 0 | 5.50193 | 28.0588 | 27.4681 |
| | | 0 | 0 | 4910 | 0 | 0 | 0 | 12.09 | 7.1 | 0 | 30.8653 | -62.4185 | -25.4572 |
| | | | | | | 0 | 0 | 11.65 | 6.98 | 0 | 41.8412 | -57.0736 | -27.8045 |
| | | | | | | 1 | 41 | 8.24 | 5.82 | 0 | 21.7595 | -64.7797 | -20.2698 |
| | | | | | | 11 | 267 | 7.15 | 5.35 | 0 | 45.4377 | -48.9693 | -21.0094 |
| | | 0 | 0 | 5150 | 0 | 0 | 0 | 12.03 | 7.08 | 0 | -29.2535 | -55.5665 | -56.6732 |
| | | | | | | 0 | 2 | 10.32 | 6.58 | 0 | -20.6012 | -70.2643 | -53.0392 |
| | | | | | | 0 | 3 | 9.86 | 6.43 | 0 | -33.7441 | -57.6628 | -48.7088 |
| | | | | | | 0 | 3 | 9.82 | 6.41 | 0 | -13.5954 | -59.7617 | -54.9722 |
| | | | | | | 0 | 0.02 | 8.65 | 5.99 | 0 | -10.817 | -73.6229 | -50.2117 |
| | | 0 | 0 | 983 | 0 | 0 | 2 | 10.23 | 6.55 | 0 | 18.2721 | -23.3547 | -12.5048 |
| | | | | | | 0 | 6 | 9.38 | 6.26 | 0 | 15.4956 | -5.88447 | -9.98842 |
| | | | | | | 1 | 37 | 8.3 | 5.85 | 0 | 13.6385 | -37.0798 | 0.351454 |
| | | 0 | 0 | 512 | 0 | 0 | 6 | 9.38 | 6.26 | 0 | 15.8009 | -11.9846 | 5.87965 |
| | | | | | | 0 | 9 | 9.14 | 6.17 | 0 | 16.5395 | -23.6246 | 6.92543 |
| | | 0 | 0 | 720 | 0 | 0 | 7 | 9.26 | 6.22 | 0 | 1.16677 | -65.6625 | -32.0152 |
| | | | | | | 9 | 245 | 7.21 | 5.38 | 0 | -0.772161 | -73.7557 | -36.6379 |
| | | 0 | 0 | 1423 | 0 | 0 | 12 | 8.97 | 6.11 | 0 | 10.8337 | -62.9633 | -41.8864 |
| | | | | | | 0 | 16 | 8.8 | 6.04 | 0 | 21.8409 | -77.7972 | -51.5488 |
| | | | | | | 2 | 58 | 8.03 | 5.74 | 0 | 23.3851 | -57.4585 | -54.1742 |
| | | 0 | 0 | 583 | 0 | 0 | 16 | 8.77 | 6.03 | 0 | 47.1695 | 47.3219 | 4.14036 |
| | | 0 | 0 | 321 | 0 | 0 | 19 | 8.68 | 6 | 0 | -15.5962 | -31.7605 | 73.3966 |
| | | 0 | 1 | 132 | 0 | 1 | 22 | 8.59 | 5.96 | 0 | -45.0444 | 41.8483 | 4.52499 |
| | | 0 | 1 | 118 | 0 | 1 | 29 | 8.45 | 5.91 | 0 | -51.7911 | -55.5593 | 12.0672 |
| | | 0 | 0 | 889 | 0 | 1 | 0.03 | 8.41 | 5.89 | 0 | 39.2172 | 39.7358 | 24.0179 |
| | | | | | | 1 | 53 | 8.1 | 5.77 | 0 | 43.103 | 49.7115 | 17.3195 |
| | | 0 | 0 | 152 | 0 | 1 | 31 | 8.39 | 5.89 | 0 | -19.2459 | -57.2082 | 71.24 |
| | | 0 | 0 | 219 | 0 | 2 | 59 | 8.02 | 5.73 | 0 | -42.2821 | -12.819 | 66.7078 |
| | | 0 | 0 | 569 | 0 | 3 | 91 | 7.77 | 5.63 | 0 | -46.72 | 41.1301 | 26.5938 |
| | | | | | | 3 | 99 | 7.72 | 5.61 | 0 | -41.531 | 44.9291 | 17.0072 |
| | | | | | | 4 | 131 | 7.57 | 5.54 | 0 | -43.2214 | 30.7942 | 29.2637 |
| | | 0 | 0 | 215 | 0 | 3 | 97 | 7.73 | 5.61 | 0 | 26.9913 | -43.0914 | -52.8518 |
| | | | | | | 13 | 314 | 7.05 | 5.3 | 0 | 31.6784 | -39.2019 | -42.8512 |
| | | 0 | 0 | 195 | 0 | 4 | 128 | 7.58 | 5.54 | 0 | -59.8514 | -68.9178 | -0.51016 |

| | | | | | | | | | | | |
|---|---|---|---|---|---|---|---|---|---|---|---|
| 0 | 0 | 144 | 0 | 5 | 138 | 7.52 | 5.52 | 0 | -11.3434 | -27.4777 | 44.556 |
| | | | | 0.01 | 249 | 7.2 | 5.37 | 0 | -6.99299 | -32.0148 | 54.5402 |
| 0 | 1 | 130 | 0 | 7 | 0.2 | 7.32 | 5.43 | 0 | 41.1763 | -51.8187 | -45.9635 |
| 0 | 2 | 102 | 1 | 13 | 312 | 7.06 | 5.31 | 0 | 32.2323 | -52.9256 | -55.1502 |

HC-NoGo_OFF

| p | c | p(FWE-corr) | p(FDR-corr) | equivk | p(unc) | p(FWE-corr).1 | p(FDR-corr).1 | T | equivZ | p(unc).1 | MNI_x | MNI_y | MNI_z |
|---|---|---|---|---|---|---|---|---|---|---|---|---|---|
| **0** | 21 | 0 | 0 | 16023 | 0 | 1 | 0.01 | 8.68 | 5.78 | 0 | -54.3198 | -55.0911 | 55.0225 |
| | | | | | | 6 | 14 | 7.76 | 5.44 | 0 | -59.07 | -45.5162 | 53.1454 |
| | | | | | | 29 | 33 | 6.9 | 5.07 | 0 | -57.3734 | -58.9489 | 44.935 |
| | | | | | | 38 | 39 | 6.77 | 5.01 | 0 | -45.1708 | -58.6062 | 60.0069 |
| | | | | | | 177 | 127 | 5.94 | 4.62 | 0 | -59.3753 | -48.2237 | 34.7391 |
| | | | | | | 208 | 127 | 5.85 | 4.57 | 0 | -36.7347 | -48.0207 | 40.2673 |
| | | | | | | 404 | 165 | 5.45 | 4.35 | 0 | -60.7687 | -34.5931 | 33.8864 |
| | | | | | | 477 | 182 | 5.33 | 4.29 | 0 | -47.1298 | -39.5034 | 53.145 |
| | | | | | | 654 | 266 | 5.08 | 4.15 | 0 | -42.5178 | -70.1648 | 56.0453 |
| | | | | | | 671 | 267 | 5.06 | 4.13 | 0 | -65.3564 | -28.4906 | 49.0092 |
| | | | | | | 842 | 331 | 4.8 | 3.98 | 0 | -65.0111 | -18.0939 | 34.7642 |
| | | | | | | 934 | 376 | 4.6 | 3.86 | 0 | -59.6186 | -26.1093 | 35.7339 |
| | | | | | | 946 | 376 | 4.56 | 3.83 | 0 | -46.9866 | -44.1929 | 42.5564 |
| | | | | | | 992 | 524 | 4.3 | 3.67 | 0 | -47.8885 | -46.3494 | 30.105 |
| | | | | | | 996 | 547 | 4.22 | 3.62 | 0 | -38.9216 | -45.8163 | 46.5411 |
| | | | | | | 1 | 735 | 3.88 | 3.38 | 0 | -58.2434 | -18.7444 | 54.2975 |
| | | 0 | 0 | 6956 | 0 | 2 | 0.01 | 8.37 | 5.67 | 0 | -20.7859 | -1.13531 | 11.0075 |
| | | | | | | 115 | 94 | 6.18 | 4.74 | 0 | -35.0411 | -1.14324 | -5.50543 |
| | | | | | | 164 | 127 | 5.99 | 4.64 | 0 | -43.2725 | 5.33599 | 0.594992 |
| | | | | | | 317 | 137 | 5.6 | 4.44 | 0 | -40.3916 | 18.7403 | 1.57521 |
| | | | | | | 393 | 165 | 5.47 | 4.36 | 0 | -44.5837 | 7.32156 | 11.7098 |
| | | | | | | 673 | 267 | 5.06 | 4.13 | 0 | -32.1678 | 20.0068 | 10.3769 |
| | | | | | | 789 | 0.3 | 4.89 | 4.03 | 0 | -54.2119 | 9.50615 | 0.421744 |
| | | | | | | 941 | 376 | 4.58 | 3.84 | 0 | -43.036 | -3.01399 | 14.0986 |
| | | | | | | 982 | 467 | 4.4 | 3.73 | 0 | -43.2275 | -13.559 | -7.49119 |
| | | | | | | 995 | 547 | 4.25 | 3.63 | 0 | -33.7165 | -12.9546 | -3.24166 |
| | | | | | | 995 | 547 | 4.24 | 3.62 | 0 | -25.5908 | 4.91491 | 16.9073 |
| | | | | | | 1 | 0.77 | 3.84 | 3.36 | 0 | -29.7853 | 10.6519 | 4.33217 |
| | | | | | | 1 | 788 | 3.82 | 3.34 | 0 | -41.1901 | -3.73 | -12.0112 |
| | | | | | | 1 | 901 | 3.59 | 3.18 | 1 | -31.1262 | -15.8793 | 10.7152 |
| | | 0 | 0 | 13610 | 0 | 12 | 0.02 | 7.39 | 5.29 | 0 | 56.4854 | -46.5346 | 52.4402 |
| | | | | | | 25 | 31 | 6.99 | 5.12 | 0 | 51.1468 | -33.7316 | 38.6956 |
| | | | | | | 87 | 78 | 6.33 | 4.81 | 0 | 66.3422 | -37.9365 | 41.5768 |
| | | | | | | 214 | 127 | 5.84 | 4.56 | 0 | 36.5694 | -38.5974 | 45.0667 |
| | | | | | | 275 | 128 | 5.69 | 4.48 | 0 | 61.2373 | -34.0813 | 53.8442 |
| | | | | | | 0.74 | 282 | 4.96 | 4.08 | 0 | 67.9226 | -28.232 | 39.2775 |
| | | | | | | 891 | 347 | 4.71 | 3.92 | 0 | 69.6955 | -17.7348 | 24.6935 |
| | | | | | | 923 | 372 | 4.63 | 3.88 | 0 | 64.9379 | -50.0135 | 38.6415 |
| | | | | | | 927 | 376 | 4.62 | 3.87 | 0 | 35.9976 | -30.5835 | 37.5485 |
| | | | | | | 939 | 376 | 4.58 | 3.85 | 0 | 58.8718 | -25.2628 | 48.0126 |
| | | | | | | 985 | 485 | 4.37 | 3.71 | 0 | 41.4727 | -47.3936 | 47.3332 |
| | | | | | | 0.99 | 515 | 4.32 | 3.68 | 0 | 64.5666 | -18.4553 | 40.9396 |
| | | | | | | 995 | 547 | 4.24 | 3.63 | 0 | 66.6963 | -38.8925 | 24.6355 |
| | | | | | | 997 | 557 | 4.19 | 3.59 | 0 | 54.7913 | -54.5452 | 29.6041 |
| | | | | | | 1 | 621 | 4.05 | 3.5 | 0 | 50.8173 | -40.5826 | 64.1874 |
| | | | | | | 1 | 848 | 3.68 | 3.24 | 1 | 46.1311 | -66.7258 | 54.1158 |
| | | 0 | 0 | 2904 | 0 | 89 | 78 | 6.32 | 4.8 | 0 | 0.276325 | 7.88506 | 53.9831 |
| | | | | | | 419 | 165 | 5.42 | 4.34 | 0 | -5.98703 | 1.10905 | 48.6979 |

| | | | | | | | | | | | |
|---|---|---|---|---|---|---|---|---|---|---|---|
| | | | | 0.95 | 381 | 4.55 | 3.82 | 0 | -6.28942 | 12.7251 | 46.0318 |
| | | | | 1 | 0.72 | 3.9 | 3.4 | 0 | 4.69349 | 13.9511 | 38.9197 |
| | | | | 1 | 848 | 3.67 | 3.23 | 1 | 7.12314 | 5.74753 | 42.2933 |
| 0.01 | 5 | 1103 | 0 | 247 | 128 | 5.75 | 4.52 | 0 | 41.1063 | -50.9658 | -25.7626 |
| | | | | 993 | 525 | 4.29 | 3.66 | 0 | 26.309 | -51.3144 | -28.5348 |
| | | | | 1 | 635 | 4 | 3.46 | 0 | 46.6826 | -53.4297 | -34.159 |
| | | | | 1 | 683 | 3.95 | 3.43 | 0 | 40.6816 | -42.1182 | -15.768 |
| | | | | 1 | 823 | 3.74 | 3.29 | 1 | 49.9325 | -42.0133 | -14.2123 |
| 0 | 0 | 2202 | 0 | 276 | 128 | 5.69 | 4.48 | 0 | -32.2777 | -41.3534 | -35.4299 |
| | | | | 786 | 0.3 | 4.89 | 4.03 | 0 | -35.1825 | -51.9064 | -31.5663 |
| | | | | 847 | 331 | 4.79 | 3.97 | 0 | -40.4382 | -59.3435 | -24.2319 |
| | | | | 1 | 621 | 4.04 | 3.5 | 0 | -46.8262 | -61.3045 | -31.3979 |
| | | | | 1 | 848 | 3.69 | 3.25 | 1 | -36.8639 | -39.7276 | -44.4227 |
| 0 | 0 | 6065 | 0 | 404 | 165 | 5.45 | 4.35 | 0 | 38.3866 | 20.6706 | 5.66092 |
| | | | | 412 | 165 | 5.44 | 4.35 | 0 | 36.293 | 3.71102 | 9.72267 |
| | | | | 532 | 206 | 5.25 | 4.25 | 0 | 45.4202 | 17.3527 | -1.96958 |
| | | | | 543 | 206 | 5.24 | 4.24 | 0 | 49.3479 | 14.4311 | 9.15959 |
| | | | | 718 | 282 | 4.99 | 4.09 | 0 | 40.6786 | -5.54984 | -8.52893 |
| | | | | 0.74 | 282 | 4.96 | 4.08 | 0 | 50.6717 | 14.512 | -7.71001 |
| | | | | 775 | 0.3 | 4.91 | 4.04 | 0 | 28.3876 | 27.2428 | 8.18049 |
| | | | | 946 | 376 | 4.56 | 3.83 | 0 | 37.5474 | 13.1287 | 0.90057 |
| | | | | 996 | 547 | 4.22 | 3.62 | 0 | 47.441 | -0.97793 | 16.6192 |
| | | | | 999 | 621 | 4.07 | 3.51 | 0 | 37.8867 | 2.87316 | -15.2623 |
| | | | | 1 | 635 | 4 | 3.47 | 0 | 31.3168 | 22.9317 | -9.20639 |
| | | | | 1 | 0.72 | 3.91 | 3.4 | 0 | 39.4666 | 30.6055 | -1.03786 |
| | | | | 1 | 823 | 3.78 | 3.31 | 0 | 31.6615 | 15.2079 | -14.1441 |
| | | | | 1 | 0.99 | 3.49 | 3.1 | 1 | 33.1061 | 0.245471 | -7.42079 |
| 201 | 101 | 508 | 11 | 0.45 | 176 | 5.38 | 4.31 | 0 | 9.23553 | 2.89368 | 74.8406 |
| | | | | 701 | 281 | 5.02 | 4.11 | 0 | 5.48117 | -7.59227 | 72.416 |
| | | | | 1 | 823 | 3.74 | 3.29 | 1 | 19.5482 | -7.85524 | 65.8398 |
| 866 | 582 | 189 | 96 | 737 | 282 | 4.97 | 4.08 | 0 | -28.0494 | 20.3184 | -11.4611 |
| 1 | 1 | 1619 | 0 | 741 | 282 | 4.96 | 4.08 | 0 | -21.3769 | -63.8961 | -56.0881 |
| | | | | 915 | 364 | 4.65 | 3.89 | 0 | -31.3763 | -53.2056 | -46.0658 |
| | | | | 971 | 421 | 4.46 | 3.77 | 0 | -37.7778 | -52.083 | -53.2232 |
| | | | | 997 | 547 | 4.21 | 3.61 | 0 | -9.19223 | -62.0962 | -52.5996 |
| 422 | 222 | 366 | 26 | 832 | 325 | 4.82 | 3.99 | 0 | 15.4467 | 5.98043 | 7.06146 |
| | | | | 886 | 346 | 4.72 | 3.93 | 0 | 23.8938 | 5.41118 | 1.72737 |
| 936 | 589 | 152 | 132 | 879 | 346 | 4.73 | 3.94 | 0 | 53.6272 | -30.4349 | 11.0756 |
| 945 | 0.59 | 146 | 139 | 882 | 346 | 4.72 | 3.93 | 0 | 31.5157 | -16.6559 | 6.99609 |
| 811 | 0.52 | 212 | 0.08 | 906 | 358 | 4.67 | 3.9 | 0 | 51.348 | 6.19164 | -34.7223 |
| 751 | 493 | 235 | 66 | 995 | 547 | 4.24 | 3.63 | 0 | -7.49232 | -15.3998 | 45.2438 |
| 922 | 584 | 161 | 122 | 996 | 547 | 4.21 | 3.61 | 0 | -20.7582 | 48.5173 | -14.8395 |
| 911 | 584 | 167 | 115 | 997 | 547 | 4.21 | 3.61 | 0 | -59.9746 | -69.0923 | 3.126 |
| 889 | 584 | 178 | 105 | 999 | 598 | 4.12 | 3.55 | 0 | -41.4207 | 38.2124 | 9.42507 |
| 925 | 584 | 159 | 124 | 999 | 598 | 4.12 | 3.55 | 0 | 22.4511 | -60.8071 | -53.186 |
| | | | | 1 | 824 | 3.74 | 3.28 | 1 | 30.6566 | -57.5038 | -55.1457 |
| 767 | 493 | 229 | 0.07 | 1 | 621 | 4.05 | 3.5 | 0 | 54.4538 | -43.1457 | 13.7946 |
| 981 | 766 | 113 | 189 | 1 | 764 | 3.85 | 3.37 | 0 | 1.14261 | -72.7159 | -40.3112 |

PDnoLID-NoGo_OFF

| p | c | p(FWE-corr) | p(FDR-corr) | equivk | p(unc) | p(FWE-corr).1 | p(FDR-corr).1 | T | equivZ | p(unc).1 | MNI_x | MNI_y | MNI_z |
|---|---|---|---|---|---|---|---|---|---|---|---|---|---|
| **0** | 19 | 0 | 0 | 13657 | 0 | 1 | 3 | 9.54 | 5.87 | 0 | 60.0767 | -38.0497 | 38.0454 |
| | | | | | | 17 | 8 | 7.92 | 5.33 | 0 | 65.3381 | -43.5304 | 30.2974 |
| | | | | | | 62 | 0.02 | 7.13 | 5.03 | 0 | 45.3867 | -44.5393 | 48.7103 |
| | | | | | | 138 | 34 | 6.64 | 4.82 | 0 | 54.3212 | -35.4204 | 52.5979 |
| | | | | | | 166 | 35 | 6.53 | 4.77 | 0 | 54.6558 | -47.9096 | 43.9391 |
| | | | | | | 257 | 49 | 6.25 | 4.65 | 0 | 63.0147 | -52.2523 | 41.091 |
| | | | | | | 278 | 49 | 6.2 | 4.62 | 0 | 50.6667 | -46.0553 | 56.8452 |
| | | | | | | 378 | 62 | 5.98 | 4.52 | 0 | 48.5277 | -40.5575 | 63.2221 |
| | | | | | | 385 | 62 | 5.97 | 4.51 | 0 | 53.1275 | -26.0462 | 55.1253 |
| | | | | | | 641 | 106 | 5.55 | 4.31 | 0 | 62.6977 | -17.5418 | 22.015 |
| | | | | | | 0.8 | 149 | 5.3 | 4.18 | 0 | 53.1162 | -46.0263 | 33.2415 |
| | | | | | | 981 | 263 | 4.79 | 3.9 | 0 | 41.1071 | -46.5576 | 69.0948 |
| | | | | | | 989 | 285 | 4.72 | 3.85 | 0 | 56.3565 | -20.4917 | 38.3682 |
| | | | | | | 999 | 376 | 4.45 | 3.69 | 0 | 46.9463 | -23.3649 | 43.0945 |
| | | 0 | 0 | 14934 | 0 | 2 | 3 | 9.36 | 5.81 | 0 | -59.3848 | -44.3683 | 44.5915 |
| | | | | | | 7 | 5 | 8.42 | 5.51 | 0 | -65.171 | -48.1803 | 33.7377 |
| | | | | | | 41 | 15 | 7.38 | 5.13 | 0 | -44.6944 | -44.4938 | 44.7353 |
| | | | | | | 115 | 0.03 | 6.75 | 4.87 | 0 | -60.7657 | -26.3117 | 33.5768 |
| | | | | | | 162 | 35 | 6.54 | 4.78 | 0 | -51.2447 | -52.2728 | 49.8025 |
| | | | | | | 164 | 35 | 6.54 | 4.78 | 0 | -51.0776 | -49.1725 | 31.2031 |
| | | | | | | 199 | 41 | 6.41 | 4.72 | 0 | -56.6429 | -32.6639 | 51.4947 |
| | | | | | | 248 | 49 | 6.27 | 4.66 | 0 | -46.1174 | -58.4797 | 57.6513 |
| | | | | | | 278 | 49 | 6.2 | 4.62 | 0 | -48.8293 | -31.649 | 43.403 |
| | | | | | | 307 | 54 | 6.13 | 4.59 | 0 | -54.2367 | -35.0228 | 60.298 |
| | | | | | | 671 | 111 | 5.51 | 4.28 | 0 | -53.5523 | -60.2373 | 50.7205 |
| | | | | | | 939 | 215 | 4.99 | 4.01 | 0 | -36.5026 | -45.4665 | 32.9906 |
| | | | | | | 973 | 248 | 4.84 | 3.92 | 0 | -43.4704 | -39.0778 | 28.1456 |
| | | | | | | 998 | 0.36 | 4.52 | 3.74 | 0 | -54.7051 | -36.4779 | 30.2087 |
| | | 0 | 0 | 4095 | 0 | 4 | 5 | 8.89 | 5.67 | 0 | -44.8664 | 15.7927 | 1.76682 |
| | | | | | | 69 | 21 | 7.06 | 5 | 0 | -32.1133 | 25.7328 | 4.55612 |
| | | | | | | 924 | 207 | 5.03 | 4.03 | 0 | -41.7889 | -2.90528 | 11.7048 |
| | | | | | | 992 | 298 | 4.68 | 3.83 | 0 | -58.4929 | 14.4827 | -2.62844 |
| | | | | | | 1 | 584 | 4.06 | 3.45 | 0 | -30.3954 | 25.6817 | -9.35449 |
| | | | | | | 1 | 0.75 | 3.81 | 3.28 | 1 | -47.9905 | 0.64486 | 3.68143 |
| | | 0 | 0 | 10174 | 0 | 6 | 5 | 8.58 | 5.56 | 0 | 3.55902 | 2.98961 | 65.8008 |
| | | | | | | 8 | 5 | 8.37 | 5.49 | 0 | 1.19089 | 6.87396 | 57.9164 |
| | | | | | | 37 | 14 | 7.45 | 5.15 | 0 | 7.73887 | 25.9388 | 47.1419 |
| | | | | | | 55 | 19 | 7.2 | 5.06 | 0 | 2.13958 | 15.6042 | 63.5497 |
| | | | | | | 381 | 62 | 5.98 | 4.52 | 0 | -1.25649 | 11.1668 | 49.9901 |
| | | | | | | 465 | 74 | 5.83 | 4.45 | 0 | 2.79808 | 16.518 | 44.5903 |
| | | | | | | 963 | 234 | 4.9 | 3.95 | 0 | 9.94924 | -9.12771 | 75.9865 |
| | | | | | | 967 | 239 | 4.88 | 3.94 | 0 | 7.60077 | 6.95672 | 36.4439 |
| | | | | | | 986 | 273 | 4.75 | 3.87 | 0 | -9.2795 | 20.4846 | 27.4744 |
| | | | | | | 994 | 311 | 4.65 | 3.81 | 0 | 6.6961 | -3.71431 | 40.3748 |
| | | | | | | 1 | 451 | 4.3 | 3.6 | 0 | -2.53184 | 20.9596 | 37.6164 |
| | | | | | | 1 | 535 | 4.14 | 3.5 | 0 | 5.23067 | 30.3009 | 30.0604 |
| | | | | | | 1 | 0.62 | 3.99 | 3.41 | 0 | 9.5106 | 20.8299 | 30.3317 |
| | | 0 | 0 | 5982 | 0 | 11 | 6 | 8.16 | 5.42 | 0 | 33.326 | 22.3893 | 5.70344 |

| | | | | | | | | | | | | | |
|---|---|---|---|---|---|---|---|---|---|---|---|---|---|
| | | | | | | 347 | 61 | 6.04 | 4.55 | 0 | 46.0325 | 15.1947 | 0.842053 |
| | | | | | | 417 | 67 | 5.91 | 4.49 | 0 | 48.435 | 23.3934 | 14.5533 |
| | | | | | | 552 | 91 | 5.69 | 4.38 | 0 | 36.7895 | 27.0083 | -6.02027 |
| | | | | | | 579 | 96 | 5.64 | 4.35 | 0 | 57.0829 | 11.9273 | 1.70123 |
| | | | | | | 947 | 218 | 4.96 | 3.99 | 0 | 43.9086 | 2.43122 | 12.3796 |
| | | | | | | 999 | 366 | 4.47 | 3.71 | 0 | 37.7881 | 8.60592 | 2.49563 |
| | | | | | | 1 | 0.43 | 4.35 | 3.63 | 0 | 50.3861 | 9.58201 | 26.4727 |
| | | | | | | 1 | 451 | 4.3 | 3.6 | 0 | 47.5179 | 23.025 | -11.1941 |
| | | | | | | 1 | 586 | 4.06 | 3.45 | 0 | 46.848 | 30.9609 | 4.981 |
| | | 0 | 0 | 2694 | 0 | 93 | 26 | 6.88 | 4.92 | 0 | -46.5582 | 32.8558 | 28.7549 |
| | | | | | | 543 | 91 | 5.7 | 4.38 | 0 | -44.6218 | 41.248 | 24.3647 |
| | | | | | | 787 | 148 | 5.32 | 4.19 | 0 | -36.0116 | 49.4929 | 13.6894 |
| | | | | | | 954 | 226 | 4.93 | 3.98 | 0 | -41.8434 | 41.5751 | 33.9295 |
| | | | | | | 998 | 358 | 4.53 | 3.74 | 0 | -28.3587 | 46.6141 | 20.1976 |
| | | | | | | 999 | 366 | 4.47 | 3.7 | 0 | -41.5189 | 24.6982 | 42.9359 |
| | | 0 | 0 | 1884 | 0 | 138 | 34 | 6.64 | 4.82 | 0 | -36.6327 | -54.1653 | -53.5191 |
| | | | | | | 623 | 105 | 5.58 | 4.32 | 0 | -31.8912 | -64.6526 | -49.6171 |
| | | | | | | 999 | 366 | 4.47 | 3.7 | 0 | -26.6941 | -69.4696 | -44.1846 |
| | | 561 | 0.12 | 232 | 25 | 204 | 41 | 6.4 | 4.71 | 0 | 24.6099 | 55.7628 | 30.7795 |
| | | 142 | 29 | 407 | 5 | 456 | 74 | 5.84 | 4.45 | 0 | -0.555715 | -23.855 | 45.3294 |
| | | | | | | 996 | 328 | 4.6 | 3.78 | 0 | 7.78648 | -28.3518 | 49.0041 |
| | | | | | | 1 | 517 | 4.18 | 3.53 | 0 | 3.76965 | -16.5787 | 42.9912 |
| | | 664 | 148 | 206 | 33 | 777 | 147 | 5.34 | 4.2 | 0 | -11.9583 | 6.08865 | 71.333 |
| | | 0.46 | 98 | 260 | 19 | 804 | 149 | 5.29 | 4.17 | 0 | 17.1555 | -72.6172 | -43.5203 |
| | | 34 | 7 | 586 | 1 | 808 | 149 | 5.28 | 4.17 | 0 | 52.7166 | 3.94674 | 50.8763 |
| | | | | | | 1 | 451 | 4.3 | 3.6 | 0 | 58.2439 | 15.238 | 36.4975 |
| | | 12 | 3 | 728 | 0 | 815 | 0.15 | 5.27 | 4.16 | 0 | -39.4316 | -55.0712 | -25.6209 |
| | | 246 | 49 | 340 | 9 | 933 | 0.21 | 5.01 | 4.02 | 0 | 47.5512 | 0.825306 | -32.6769 |
| | | 971 | 0.36 | 109 | 107 | 946 | 218 | 4.96 | 3.99 | 0 | 34.1402 | -41.512 | -49.2775 |
| | | 961 | 0.36 | 116 | 98 | 997 | 342 | 4.57 | 3.76 | 0 | 62.8109 | -24.7968 | -9.76608 |
| | | 925 | 328 | 133 | 78 | 999 | 365 | 4.49 | 3.72 | 0 | -47.612 | 7.46932 | 21.312 |
| | | 941 | 337 | 126 | 86 | 1 | 535 | 4.14 | 3.5 | 0 | 16.2297 | 11.5416 | 68.2665 |
| | | 973 | 0.36 | 108 | 109 | 1 | 535 | 4.14 | 3.5 | 0 | 41.7768 | 39.8611 | 34.2201 |

LID-NoGo_OFF

| p | c | p(FWE-corr) | p(FDR-corr) | equivk | p(unc) | p(FWE-corr).1 | p(FDR-corr).1 | T | equivZ | p(unc).1 | MNI_x | MNI_y | MNI_z |
|---|---|---|---|---|---|---|---|---|---|---|---|---|---|
| **0** | 28 | 0 | 0 | 16224 | 0 | 0 | 1 | 9.31 | 6.23 | 0 | 52.4006 | 13.4764 | -1.73292 |
| | | | | | | 0 | 1 | 8.98 | 6.11 | 0 | 31.3187 | 2.26287 | 0.0600546 |
| | | | | | | 1 | 1 | 8.53 | 5.94 | 0 | 38.4166 | 19.3456 | 1.86428 |
| | | | | | | 7 | 4 | 7.4 | 5.47 | 0 | 53.1563 | 11.2556 | 19.9862 |
| | | | | | | 33 | 9 | 6.67 | 5.12 | 0 | 40.7439 | 6.62676 | -2.76584 |
| | | | | | | 58 | 12 | 6.41 | 4.99 | 0 | 39.1613 | 2.17408 | 13.6347 |
| | | | | | | 211 | 0.03 | 5.78 | 4.65 | 0 | 26.0865 | 28.4913 | 3.4105 |
| | | | | | | 426 | 0.05 | 5.4 | 4.43 | 0 | 46.4453 | -0.0643954 | -4.94258 |
| | | | | | | 0.46 | 54 | 5.35 | 4.4 | 0 | 38.5332 | 29.722 | -3.34129 |
| | | | | | | 513 | 61 | 5.28 | 4.36 | 0 | 47.13 | 30.4625 | 6.40936 |
| | | | | | | 601 | 72 | 5.16 | 4.29 | 0 | 34.7235 | 24.9271 | -10.7015 |
| | | | | | | 734 | 91 | 4.99 | 4.19 | 0 | 28.4916 | 9.10903 | 6.62157 |
| | | | | | | 852 | 0.12 | 4.82 | 4.08 | 0 | 40.0984 | -1.95566 | -15.4568 |
| | | | | | | 993 | 253 | 4.37 | 3.78 | 0 | 33.3207 | -7.49682 | -1.19503 |
| | | | | | | 997 | 284 | 4.29 | 3.73 | 0 | 25.4299 | -3.76867 | 16.6451 |
| | | | | | | 1 | 544 | 3.85 | 3.42 | 0 | 41.3391 | -2.73795 | 2.56211 |
| | | | | | | 1 | 871 | 3.52 | 3.18 | 1 | 36.267 | 21.8884 | 15.526 |
| | | 0 | 0 | 19872 | 0 | 0 | 1 | 8.82 | 6.05 | 0 | 9.12489 | 9.21907 | 72.6934 |
| | | | | | | 1 | 1 | 8.42 | 5.9 | 0 | -0.441976 | -9.9792 | 67.3244 |
| | | | | | | 3 | 3 | 7.76 | 5.62 | 0 | 0.878389 | 2.01497 | 70.309 |
| | | | | | | 3 | 3 | 7.75 | 5.62 | 0 | 7.89938 | 17.1154 | 40.9948 |
| | | | | | | 41 | 0.01 | 6.57 | 5.07 | 0 | 0.893013 | 13.3548 | 65.8707 |
| | | | | | | 82 | 15 | 6.25 | 4.9 | 0 | -3.17791 | 13.1022 | 40.2863 |
| | | | | | | 86 | 15 | 6.22 | 4.89 | 0 | 1.36704 | 2.83174 | 43.9207 |
| | | | | | | 139 | 22 | 5.99 | 4.77 | 0 | 19.4686 | 7.70563 | 64.9823 |
| | | | | | | 177 | 27 | 5.87 | 4.7 | 0 | 9.35662 | -16.8989 | 41.8099 |
| | | | | | | 209 | 0.03 | 5.79 | 4.65 | 0 | 0.847816 | -8.49967 | 47.0089 |
| | | | | | | 0.24 | 32 | 5.71 | 4.61 | 0 | -7.8406 | -18.6545 | 69.7459 |
| | | | | | | 509 | 0.06 | 5.28 | 4.36 | 0 | -15.6259 | 1.59408 | 76.5389 |
| | | | | | | 0.61 | 72 | 5.15 | 4.28 | 0 | 8.44785 | 29.1884 | 27.1446 |
| | | | | | | 672 | 81 | 5.07 | 4.24 | 0 | -40.1332 | -11.6687 | 66.9431 |
| | | | | | | 702 | 86 | 5.04 | 4.21 | 0 | 5.86457 | -15.0292 | 76.5144 |
| | | | | | | 752 | 95 | 4.97 | 4.17 | 0 | -19.4279 | 9.35649 | 68.7514 |
| | | | | | | 865 | 125 | 4.8 | 4.07 | 0 | -1.53493 | 9.31336 | 28.1229 |
| | | | | | | 0.92 | 151 | 4.69 | 4 | 0 | -8.1365 | 25.1227 | 24.8563 |
| | | | | | | 927 | 153 | 4.67 | 3.98 | 0 | -8.80711 | 24.891 | 42.8438 |
| | | | | | | 929 | 154 | 4.67 | 3.98 | 0 | 7.35235 | -24.7211 | 70.3443 |
| | | | | | | 948 | 169 | 4.62 | 3.95 | 0 | -18.9274 | -13.8578 | 73.5902 |
| | | | | | | 0.96 | 177 | 4.58 | 3.92 | 0 | -11.6413 | 36.9276 | 26.3288 |
| | | | | | | 966 | 182 | 4.55 | 3.91 | 0 | -11.2107 | -10.0943 | 46.6939 |
| | | | | | | 994 | 257 | 4.36 | 3.77 | 0 | -30.1628 | -9.46491 | 70.7035 |
| | | | | | | 999 | 338 | 4.19 | 3.66 | 0 | -4.90865 | -24.471 | 74.3811 |
| | | | | | | 1 | 475 | 3.95 | 3.49 | 0 | -11.642 | -24.6306 | 59.6552 |
| | | | | | | 1 | 873 | 3.52 | 3.17 | 1 | 26.5718 | -2.8095 | 73.8112 |
| | | 0 | 0 | 12223 | 0 | 1 | 1 | 8.44 | 5.91 | 0 | -36.3165 | -53.0119 | -31.5373 |
| | | | | | | 1 | 1 | 8.37 | 5.88 | 0 | -34.1987 | -61.4308 | -52.0032 |
| | | | | | | 9 | 4 | 7.29 | 5.41 | 0 | -41.6161 | -62.3903 | -25.322 |
| | | | | | | 117 | 0.02 | 6.07 | 4.81 | 0 | -24.3874 | -68.5825 | -56.2027 |
| | | | | | | 125 | 21 | 6.04 | 4.8 | 0 | -31.7197 | -62.7931 | -22.8249 |
| | | | | | | 0.16 | 25 | 5.92 | 4.73 | 0 | -59.0649 | -67.4375 | -7.71403 |

| | | | | | | | | | | | | | |
|---|---|---|---|---|---|---|---|---|---|---|---|---|---|
| | | | | | | 191 | 29 | 5.83 | 4.68 | 0 | -21.388 | -56.8867 | -55.9329 |
| | | | | | | 657 | 78 | 5.09 | 4.25 | 0 | -17.7103 | -54.9497 | -19.1243 |
| | | | | | | 719 | 89 | 5.01 | 4.2 | 0 | -16.9665 | -74.4464 | -53.684 |
| | | | | | | 987 | 224 | 4.43 | 3.83 | 0 | -47.9669 | -53.5589 | -32.3519 |
| | | | | | | 998 | 0.3 | 4.26 | 3.71 | 0 | -44.0077 | -50.3192 | -21.1066 |
| | | | | | | 1 | 371 | 4.13 | 3.62 | 0 | -30.6737 | -42.0104 | -33.5323 |
| | | | | | | 1 | 471 | 3.96 | 3.5 | 0 | -51.6113 | -65.282 | -18.9284 |
| | | | | | | 1 | 557 | 3.83 | 3.4 | 0 | -36.7917 | -50.9547 | -44.5449 |
| | | | | | | 1 | 686 | 3.7 | 3.31 | 0 | -21.8119 | -63.7499 | -27.6592 |
| | | | | | | 1 | 695 | 3.69 | 3.3 | 0 | -43.8907 | -41.4076 | -22.6157 |
| | | | | | | 1 | 875 | 3.52 | 3.17 | 1 | -11.189 | -63.1877 | -51.226 |
| | | 0 | 0 | 40013 | 0 | 1 | 2 | 8.24 | 5.82 | 0 | -53.978 | -37.1994 | 53.8717 |
| | | | | | | 8 | 4 | 7.34 | 5.44 | 0 | -57.3792 | -58.811 | 46.1551 |
| | | | | | | 9 | 4 | 7.29 | 5.41 | 0 | -37.5477 | 5.72468 | 3.18966 |
| | | | | | | 16 | 7 | 7.02 | 5.29 | 0 | -40.4174 | 19.031 | 4.26356 |
| | | | | | | 17 | 7 | 6.98 | 5.27 | 0 | -58.0117 | 3.89312 | 4.14124 |
| | | | | | | 0.02 | 7 | 6.9 | 5.23 | 0 | -41.7363 | -4.09737 | 14.069 |
| | | | | | | 24 | 8 | 6.82 | 5.19 | 0 | -51.6659 | -23.7075 | 20.6747 |
| | | | | | | 25 | 8 | 6.81 | 5.19 | 0 | -51.53 | -49.2339 | 50.9215 |
| | | | | | | 36 | 9 | 6.63 | 5.1 | 0 | -30.9256 | 24.9391 | 9.87367 |
| | | | | | | 39 | 0.01 | 6.6 | 5.08 | 0 | -55.2896 | 4.6004 | 17.9725 |
| | | | | | | 53 | 12 | 6.45 | 5.01 | 0 | -44.5248 | 0.800508 | -18.9079 |
| | | | | | | 0.07 | 14 | 6.32 | 4.94 | 0 | -38.8561 | -17.8144 | 3.4452 |
| | | | | | | 129 | 21 | 6.03 | 4.79 | 0 | -50.2113 | 13.5288 | -1.87775 |
| | | | | | | 137 | 22 | 6 | 4.77 | 0 | -37.2222 | -23.8658 | 18.5395 |
| | | | | | | 189 | 29 | 5.84 | 4.68 | 0 | -68.0208 | -34.4134 | 18.1369 |
| | | | | | | 225 | 31 | 5.75 | 4.63 | 0 | -66.7686 | -32.4782 | 30.4602 |
| | | | | | | 265 | 34 | 5.66 | 4.58 | 0 | -60.5975 | -46.709 | 43.279 |
| | | | | | | 271 | 35 | 5.65 | 4.58 | 0 | -53.2464 | -50.7087 | 32.1784 |
| | | | | | | 271 | 35 | 5.65 | 4.58 | 0 | -39.7287 | -10.4621 | -6.23635 |
| | | | | | | 313 | 0.04 | 5.57 | 4.53 | 0 | -36.2396 | -40.4834 | 42.283 |
| | | | | | | 319 | 41 | 5.56 | 4.53 | 0 | -43.3135 | -15.505 | 11.7627 |
| | | | | | | 338 | 41 | 5.53 | 4.51 | 0 | -62.2244 | -22.763 | 31.3737 |
| | | | | | | 356 | 43 | 5.5 | 4.49 | 0 | -44.3655 | -57.9782 | 64.5843 |
| | | | | | | 387 | 47 | 5.45 | 4.46 | 0 | -49.7366 | 14.0375 | -10.8716 |
| | | | | | | 411 | 49 | 5.42 | 4.44 | 0 | -61.2232 | -22.6588 | 17.5585 |
| | | | | | | 0.46 | 54 | 5.35 | 4.4 | 0 | -40.0737 | -47.5857 | 41.7405 |
| | | | | | | 587 | 71 | 5.18 | 4.3 | 0 | -30.5171 | -6.88766 | 3.85592 |
| | | | | | | 601 | 72 | 5.16 | 4.29 | 0 | -29.9378 | 5.56089 | -1.99965 |
| | | | | | | 619 | 73 | 5.14 | 4.28 | 0 | -55.744 | 8.12794 | 28.8271 |
| | | | | | | 775 | 101 | 4.94 | 4.15 | 0 | -46.1553 | 1.41458 | 24.3355 |
| | | | | | | 787 | 103 | 4.92 | 4.14 | 0 | -55.4221 | 6.21227 | 39.344 |
| | | | | | | 898 | 139 | 4.74 | 4.03 | 0 | -49.2428 | -40.195 | 16.3058 |
| | | | | | | 952 | 172 | 4.6 | 3.94 | 0 | -42.4044 | -44.0291 | 66.9939 |
| | | | | | | 1 | 383 | 4.1 | 3.6 | 0 | -31.7007 | 4.36587 | -21.8624 |
| | | | | | | 1 | 448 | 4 | 3.53 | 0 | -25.5267 | -3.32156 | 12.1165 |
| | | | | | | 1 | 463 | 3.98 | 3.51 | 0 | -30.6805 | -3.90146 | -20.8217 |
| | | | | | | 1 | 479 | 3.94 | 3.48 | 0 | -31.3094 | 22.1232 | -6.74511 |
| | | | | | | 1 | 537 | 3.86 | 3.43 | 0 | -36.4935 | -32.1059 | 16.8159 |
| | | | | | | 1 | 807 | 3.57 | 3.21 | 1 | -35.795 | 29.2839 | -3.23866 |
| | | 0 | 0 | 29679 | 0 | 2 | 2 | 8.02 | 5.73 | 0 | 61.4568 | -19.8593 | 33.3182 |
| | | | | | | 3 | 2 | 7.85 | 5.66 | 0 | 52.2836 | -24.5024 | 38.4584 |

| | | | | | | | | | | | | | |
|---|---|---|---|---|---|---|---|---|---|---|---|---|---|
| | | | | | | 5 | 4 | 7.51 | 5.51 | 0 | 43.2722 | -35.4477 | 41.4002 |
| | | | | | | 6 | 4 | 7.43 | 5.48 | 0 | 48.8172 | -49.6071 | 54.7855 |
| | | | | | | 33 | 9 | 6.67 | 5.12 | 0 | 58.1054 | -49.1577 | 42.5268 |
| | | | | | | 33 | 9 | 6.67 | 5.12 | 0 | 50.4403 | -32.8195 | 56.7175 |
| | | | | | | 54 | 12 | 6.44 | 5 | 0 | 67.3514 | -32.335 | 27.1622 |
| | | | | | | 65 | 13 | 6.36 | 4.96 | 0 | 39.3969 | -49.666 | 41.5623 |
| | | | | | | 109 | 19 | 6.11 | 4.83 | 0 | 69.0899 | -23.6145 | 28.2534 |
| | | | | | | 147 | 23 | 5.97 | 4.75 | 0 | 66.6685 | -41.1969 | 28.0822 |
| | | | | | | 159 | 25 | 5.92 | 4.73 | 0 | 54.5904 | -32.3179 | 39.787 |
| | | | | | | 203 | 0.03 | 5.8 | 4.66 | 0 | 62.1876 | -52.2932 | 9.16551 |
| | | | | | | 217 | 0.03 | 5.77 | 4.64 | 0 | 62.8487 | -32.7068 | 45.643 |
| | | | | | | 224 | 31 | 5.75 | 4.63 | 0 | 63.8226 | -29.6183 | 35.8732 |
| | | | | | | 291 | 37 | 5.61 | 4.56 | 0 | 52.1933 | -37.7563 | 49.3677 |
| | | | | | | 347 | 42 | 5.51 | 4.5 | 0 | 59.6431 | -26.2002 | -2.50549 |
| | | | | | | 476 | 56 | 5.32 | 4.39 | 0 | 55.5137 | -43.3955 | 12.0681 |
| | | | | | | 536 | 64 | 5.25 | 4.34 | 0 | 51.3835 | -15.3982 | 16.0523 |
| | | | | | | 0.61 | 72 | 5.15 | 4.28 | 0 | 57.2963 | -27.272 | 54.0294 |
| | | | | | | 628 | 74 | 5.13 | 4.27 | 0 | 47.5322 | -38.8634 | 66.5865 |
| | | | | | | 659 | 78 | 5.09 | 4.25 | 0 | 34.8636 | -22.001 | 16.1266 |
| | | | | | | 728 | 0.09 | 5 | 4.19 | 0 | 52.0695 | -31.4397 | 0.281968 |
| | | | | | | 802 | 105 | 4.9 | 4.13 | 0 | 43.4525 | -55.0897 | 64.1762 |
| | | | | | | 0.83 | 113 | 4.86 | 4.1 | 0 | 38.3403 | -33.1005 | 32.1875 |
| | | | | | | 923 | 152 | 4.68 | 3.99 | 0 | 51.2932 | -48.5014 | 31.3266 |
| | | | | | | 997 | 279 | 4.3 | 3.74 | 0 | 68.1232 | -19.3396 | -11.8658 |
| | | | | | | 1 | 408 | 4.06 | 3.57 | 0 | 58.1796 | -63.6109 | -9.96696 |
| | | | | | | 1 | 592 | 3.79 | 3.37 | 0 | 37.8527 | -16.0143 | 2.34719 |
| | | | | | | 1 | 678 | 3.7 | 3.31 | 0 | 31.0202 | -33.2913 | 22.2999 |
| | | | | | | 1 | 712 | 3.67 | 3.29 | 1 | 58.8635 | -33.3082 | 18.1228 |
| | | | | | | 1 | 816 | 3.56 | 3.2 | 1 | 65.2422 | -53.9999 | 20.3936 |
| | | | | | | 1 | 882 | 3.5 | 3.16 | 1 | 43.5594 | -62.0671 | 59.2791 |
| | | | | | | 1 | 901 | 3.49 | 3.15 | 1 | 48.5468 | -19.8538 | 54.8615 |
| | | 0 | 0 | 2011 | 0 | 2 | 2 | 7.9 | 5.68 | 0 | -29.8734 | -11.6835 | -9.4512 |
| | | | | | | 625 | 74 | 5.13 | 4.27 | 0 | -13.8963 | -22.9245 | -19.266 |
| | | | | | | 843 | 118 | 4.84 | 4.09 | 0 | -17.9394 | -4.67472 | -12.5677 |
| | | | | | | 982 | 211 | 4.47 | 3.85 | 0 | -9.89522 | -18.3342 | -26.8221 |
| | | | | | | 999 | 353 | 4.16 | 3.64 | 0 | -8.37353 | -6.26777 | -14.9224 |
| | | 0 | 0 | 2385 | 0 | 4 | 3 | 7.65 | 5.57 | 0 | 17.3135 | -4.02575 | -19.3713 |
| | | | | | | 122 | 21 | 6.06 | 4.8 | 0 | 20.0297 | -16.1201 | -13.6109 |
| | | | | | | 0.44 | 52 | 5.37 | 4.42 | 0 | 8.06781 | -18.1851 | -16.9634 |
| | | 0 | 0 | 7427 | 0 | 9 | 4 | 7.27 | 5.4 | 0 | -45.9836 | 44.9929 | 18.1098 |
| | | | | | | 17 | 7 | 6.99 | 5.27 | 0 | -47.6866 | 30.379 | 30.2843 |
| | | | | | | 59 | 12 | 6.4 | 4.98 | 0 | -45.6656 | 24.1766 | 39.8845 |
| | | | | | | 83 | 15 | 6.24 | 4.9 | 0 | -38.9886 | 53.5389 | 21.0938 |
| | | | | | | 226 | 31 | 5.75 | 4.63 | 0 | -47.205 | 42.5854 | 6.04439 |
| | | | | | | 252 | 33 | 5.69 | 4.6 | 0 | -30.0172 | 58.4775 | 28.9995 |
| | | | | | | 387 | 47 | 5.45 | 4.46 | 0 | -45.9353 | 34.8107 | 21.2984 |
| | | | | | | 1 | 371 | 4.13 | 3.62 | 0 | -33.2711 | 38.261 | 38.1828 |
| | | | | | | 1 | 475 | 3.95 | 3.49 | 0 | -45.6097 | 52.1037 | 0.610078 |
| | | | | | | 1 | 732 | 3.64 | 3.27 | 1 | -37.5285 | 35.2697 | 13.4544 |
| | | 0 | 0 | 12137 | 0 | 13 | 6 | 7.09 | 5.32 | 0 | 41.7444 | -44.5507 | -26.7704 |
| | | | | | | 0.02 | 7 | 6.9 | 5.23 | 0 | 44.6621 | -52.7682 | -26.7563 |
| | | | | | | 27 | 8 | 6.76 | 5.16 | 0 | 38.0778 | -54.7414 | -56.0835 |

| | | | | | | | | | | | |
|---|---|---|---|---|---|---|---|---|---|---|---|
| | | | | 29 | 9 | 6.73 | 5.15 | 0 | 32.0031 | -46.7282 | -47.2927 |
| | | | | 34 | 9 | 6.66 | 5.11 | 0 | 19.9478 | -58.6262 | -51.923 |
| | | | | 54 | 12 | 6.44 | 5 | 0 | 26.9229 | -60.2866 | -37.0906 |
| | | | | 139 | 22 | 5.99 | 4.77 | 0 | 30.6663 | -69.691 | -27.3963 |
| | | | | 155 | 24 | 5.94 | 4.74 | 0 | 36.2918 | -62.0571 | -46.3203 |
| | | | | 0.25 | 33 | 5.69 | 4.6 | 0 | 41.1687 | -67.8258 | -55.9414 |
| | | | | 325 | 41 | 5.55 | 4.52 | 0 | 17.1431 | -65.906 | -44.3287 |
| | | | | 344 | 42 | 5.52 | 4.5 | 0 | 29.6372 | -54.3519 | -52.6226 |
| | | | | 362 | 44 | 5.49 | 4.49 | 0 | 32.4338 | -57.2535 | -30.0562 |
| | | | | 0.41 | 49 | 5.42 | 4.44 | 0 | 25.892 | -48.8623 | -29.6498 |
| | | | | 539 | 64 | 5.24 | 4.34 | 0 | 20.792 | -76.124 | -51.5484 |
| | | | | 717 | 89 | 5.02 | 4.2 | 0 | 14.3224 | -46.9776 | -25.7211 |
| | | | | 915 | 149 | 4.7 | 4 | 0 | 45.3495 | -39.7624 | -19.6692 |
| | | | | 998 | 305 | 4.25 | 3.7 | 0 | 29.3306 | -64.9248 | -58.3143 |
| | | | | 1 | 396 | 4.08 | 3.58 | 0 | 10.6962 | -73.9313 | -54.1152 |
| | | | | 1 | 438 | 4.01 | 3.54 | 0 | 42.2955 | -50.4533 | -43.7993 |
| | | | | 1 | 448 | 4 | 3.53 | 0 | 25.862 | -35.0089 | -38.3974 |
| 0 | 0 | 5481 | 0 | 52 | 11 | 6.46 | 5.01 | 0 | 38.8807 | 50.5709 | 30.423 |
| | | | | 325 | 41 | 5.55 | 4.52 | 0 | 40.9707 | 40.8506 | 31.7197 |
| | | | | 616 | 73 | 5.14 | 4.28 | 0 | 33.0549 | 50.0452 | 18.5296 |
| | | | | 799 | 105 | 4.9 | 4.13 | 0 | 46.3884 | 31.9641 | 36.7315 |
| | | | | 869 | 126 | 4.79 | 4.06 | 0 | 43.4746 | 45.7741 | 18.7049 |
| | | | | 944 | 166 | 4.63 | 3.95 | 0 | 51.6862 | 45.1238 | 2.32999 |
| | | | | 1 | 534 | 3.87 | 3.43 | 0 | 36.8029 | 45.4088 | 2.57921 |
| 0 | 0 | 1330 | 0 | 56 | 12 | 6.42 | 4.99 | 0 | 34.9502 | -3.56308 | -42.7006 |
| | | | | 441 | 52 | 5.37 | 4.42 | 0 | 49.5885 | 2.85831 | -41.4139 |
| | | | | 995 | 267 | 4.33 | 3.76 | 0 | 40.6295 | 5.51121 | -39.1059 |
| 121 | 27 | 442 | 4 | 85 | 15 | 6.23 | 4.89 | 0 | -15.4096 | -12.0884 | 10.3092 |
| | | | | 595 | 72 | 5.17 | 4.3 | 0 | -15.0049 | -4.28105 | 15.7429 |
| 484 | 112 | 260 | 21 | 338 | 41 | 5.53 | 4.51 | 0 | -29.0031 | -6.56102 | -35.7525 |
| | | | | 999 | 327 | 4.21 | 3.67 | 0 | -35.7689 | 1.84955 | -35.5544 |
| 576 | 129 | 234 | 28 | 456 | 54 | 5.35 | 4.4 | 0 | -20.333 | -57.3485 | 72.9489 |
| 965 | 338 | 114 | 109 | 606 | 72 | 5.16 | 4.29 | 0 | 15.2038 | 5.03613 | 14.9866 |
| 25 | 6 | 654 | 1 | 741 | 93 | 4.99 | 4.18 | 0 | 52.2838 | 9.61062 | -25.9273 |
| 898 | 295 | 145 | 74 | 764 | 98 | 4.95 | 4.16 | 0 | 16.9388 | -43.4869 | 25.249 |
| 429 | 101 | 277 | 18 | 795 | 105 | 4.91 | 4.14 | 0 | -21.6549 | 57.1741 | -8.74474 |
| 0.95 | 338 | 123 | 97 | 814 | 109 | 4.88 | 4.12 | 0 | -28.751 | -42.4849 | -50.0982 |
| 808 | 224 | 173 | 53 | 961 | 177 | 4.57 | 3.92 | 0 | 11.0589 | -3.31574 | 8.00021 |
| 936 | 324 | 130 | 89 | 973 | 193 | 4.52 | 3.88 | 0 | 35.9427 | -6.36623 | 66.6886 |
| 917 | 307 | 138 | 0.08 | 978 | 203 | 4.49 | 3.87 | 0 | -30.0083 | -82.9657 | -44.2865 |
| 508 | 113 | 253 | 23 | 981 | 209 | 4.47 | 3.85 | 0 | 1.11825 | -69.1976 | -36.8164 |
| 964 | 338 | 115 | 107 | 988 | 229 | 4.42 | 3.82 | 0 | 12.2427 | -59.7351 | -23.4391 |
| 137 | 29 | 426 | 5 | 1 | 431 | 4.03 | 3.55 | 0 | 50.8327 | -0.192024 | 49.0204 |
| | | | | 1 | 0.53 | 3.87 | 3.44 | 0 | 53.0012 | 6.91137 | 40.9558 |
| 983 | 393 | 100 | 131 | 1 | 466 | 3.97 | 3.5 | 0 | -22.2513 | 49.7235 | 1.11177 |
| 768 | 209 | 184 | 47 | 1 | 507 | 3.9 | 3.46 | 0 | 9.40876 | -37.7489 | -17.8857 |
| | | | | 1 | 711 | 3.67 | 3.29 | 1 | 16.4722 | -31.3473 | -26.1163 |
| 965 | 338 | 114 | 109 | 1 | 552 | 3.83 | 3.41 | 0 | -10.3862 | -38.3173 | -17.4234 |

HC-Reward_OFF

| p | c | p(FWE-corr) | p(FDR-corr) | equivk | p(unc) | p(FWE-corr).1 | p(FDR-corr).1 | T | equivZ | p(unc).1 | MNI_x | MNI_y | MNI_z |
|---|---|---|---|---|---|---|---|---|---|---|---|---|---|
| **0** | 49 | 0 | 0 | 6890 | 0 | 24 | 132 | 7.22 | 5.21 | 0 | -12.307 | 6.71714 | -13.7294 |
| | | | | | | 28 | 132 | 7.13 | 5.17 | 0 | -22.1728 | 11.8809 | -6.90861 |
| | | | | | | 0.04 | 132 | 6.93 | 5.09 | 0 | 17.0008 | 13.3491 | -10.3265 |
| | | | | | | 321 | 162 | 5.79 | 4.54 | 0 | 10.94 | 18.0375 | -2.37865 |
| | | | | | | 375 | 0.17 | 5.69 | 4.48 | 0 | -10.2565 | 14.2921 | -7.31481 |
| | | | | | | 0.66 | 241 | 5.27 | 4.25 | 0 | -4.42557 | 2.80718 | -10.7655 |
| | | | | | | 0.67 | 241 | 5.26 | 4.25 | 0 | 8.78909 | 8.53701 | -1.32618 |
| | | | | | | 0.71 | 254 | 5.2 | 4.21 | 0 | -25.9272 | 19.5205 | -10.4486 |
| | | | | | | 944 | 359 | 4.77 | 3.96 | 0 | -12.017 | 17.496 | 3.34304 |
| | | | | | | 1 | 0.55 | 4.2 | 3.6 | 0 | 17.1137 | 26.058 | -9.11917 |
| | | | | | | 1 | 588 | 4.04 | 3.5 | 0 | -2.46307 | 26.5221 | -12.8172 |
| | | | | | | 1 | 588 | 4.04 | 3.5 | 0 | -10.7361 | 5.40129 | 4.16443 |
| | | | | | | 1 | 722 | 3.81 | 3.34 | 0 | -8.09401 | 29.4517 | -2.43492 |
| | | 0 | 0 | 26786 | 0 | 42 | 132 | 6.92 | 5.08 | 0 | -22.9736 | -100.506 | 16.0281 |
| | | | | | | 56 | 132 | 6.76 | 5.01 | 0 | 7.63515 | -94.7867 | 14.6302 |
| | | | | | | 88 | 0.16 | 6.52 | 4.9 | 0 | -16.5914 | -91.3939 | -7.63062 |
| | | | | | | 0.13 | 0.16 | 6.31 | 4.8 | 0 | -12.5004 | -73.3771 | 25.5333 |
| | | | | | | 132 | 0.16 | 6.3 | 4.79 | 0 | 14.7658 | -99.1065 | 23.107 |
| | | | | | | 135 | 0.16 | 6.29 | 4.79 | 0 | -45.6386 | -80.9513 | 24.4009 |
| | | | | | | 139 | 0.16 | 6.27 | 4.78 | 0 | 17.321 | -97.3931 | 0.225023 |
| | | | | | | 246 | 162 | 5.95 | 4.62 | 0 | 8.33654 | -91.7027 | 34.9869 |
| | | | | | | 274 | 162 | 5.89 | 4.59 | 0 | 18.8136 | -59.5816 | 22.3943 |
| | | | | | | 293 | 162 | 5.85 | 4.57 | 0 | 45.3595 | -76.8364 | 33.513 |
| | | | | | | 311 | 162 | 5.81 | 4.55 | 0 | -28.0505 | -93.6006 | 32.0022 |
| | | | | | | 336 | 162 | 5.76 | 4.52 | 0 | -42.2487 | -88.7343 | 25.1531 |
| | | | | | | 345 | 163 | 5.74 | 4.51 | 0 | 11.9086 | -87.598 | 17.2469 |
| | | | | | | 363 | 0.17 | 5.71 | 4.5 | 0 | -15.2547 | -101.501 | 20.6277 |
| | | | | | | 473 | 211 | 5.54 | 4.4 | 0 | 14.3544 | -88.6166 | 8.28554 |
| | | | | | | 541 | 222 | 5.44 | 4.35 | 0 | -52.6882 | -80.1611 | 16.3018 |
| | | | | | | 587 | 239 | 5.37 | 4.31 | 0 | -18.6555 | -103.427 | 3.37716 |
| | | | | | | 677 | 241 | 5.25 | 4.24 | 0 | -15.0608 | -96.487 | 31.7779 |
| | | | | | | 902 | 316 | 4.88 | 4.03 | 0 | -0.324489 | -89.7654 | 30.8702 |
| | | | | | | 0.91 | 319 | 4.86 | 4.02 | 0 | 9.66236 | -55.3573 | 12.9468 |
| | | | | | | 927 | 338 | 4.82 | 3.99 | 0 | 36.9751 | -84.7762 | 32.8493 |
| | | | | | | 954 | 359 | 4.73 | 3.94 | 0 | 60.1867 | -63.3171 | 24.5053 |
| | | | | | | 955 | 359 | 4.73 | 3.94 | 0 | 56.0632 | -70.0271 | 29.2765 |
| | | | | | | 969 | 385 | 4.67 | 3.9 | 0 | -35.8649 | -84.5698 | 19.4677 |
| | | | | | | 987 | 423 | 4.55 | 3.83 | 0 | 41.6679 | -76.8288 | 17.3676 |
| | | | | | | 988 | 425 | 4.54 | 3.82 | 0 | -13.9612 | -87.0069 | 46.4259 |
| | | | | | | 994 | 452 | 4.46 | 3.77 | 0 | 31.1656 | -84.851 | 13.4821 |
| | | | | | | 995 | 463 | 4.44 | 3.76 | 0 | -37.1101 | -71.8493 | 20.5153 |
| | | | | | | 998 | 483 | 4.36 | 3.71 | 0 | -42.5313 | -71.7446 | 29.4684 |
| | | | | | | 998 | 483 | 4.36 | 3.7 | 0 | 31.6256 | -78.1647 | 36.6155 |
| | | | | | | 999 | 483 | 4.34 | 3.69 | 0 | 0.372216 | -63.5049 | 23.5369 |
| | | | | | | 999 | 494 | 4.32 | 3.68 | 0 | 22.8146 | -88.4377 | -2.423 |
| | | | | | | 999 | 499 | 4.28 | 3.66 | 0 | 35.5688 | -93.2654 | 21.9468 |
| | | | | | | 1 | 556 | 4.19 | 3.59 | 0 | 25.747 | -83.0906 | 26.4629 |
| | | | | | | 1 | 563 | 4.17 | 3.58 | 0 | 24.1199 | -99.7587 | 9.42453 |
| | | | | | | 1 | 563 | 4.13 | 3.55 | 0 | -12.4668 | -95.9874 | 10.4531 |

| | | | | | | | | | | | | | |
|---|---|---|---|---|---|---|---|---|---|---|---|---|---|
| | | | | | | 1 | 563 | 4.12 | 3.55 | 0 | 48.8919 | -63.282 | 10.2188 |
| | | | | | | 1 | 675 | 3.88 | 3.38 | 0 | -35.3576 | -95.377 | 19.8837 |
| | | | | | | 1 | 706 | 3.84 | 3.35 | 0 | 0.438625 | -92.2827 | 0.493819 |
| | | | | | | 1 | 752 | 3.76 | 3.3 | 0 | 33.6707 | -81.8617 | 6.65514 |
| | | | | | | 1 | 864 | 3.63 | 3.21 | 1 | 35.4268 | -96.8584 | 9.25676 |
| | | | | | | 1 | 869 | 3.62 | 3.2 | 1 | 15.1824 | -91.4785 | -11.9089 |
| | | | | | | 1 | 923 | 3.56 | 3.16 | 1 | 23.8172 | -67.3778 | 24.264 |
| | | 0 | 0 | 12805 | 0 | 77 | 0.16 | 6.59 | 4.93 | 0 | -40.775 | 46.7306 | 20.9726 |
| | | | | | | 179 | 162 | 6.13 | 4.71 | 0 | -46.2122 | 51.6128 | 6.78371 |
| | | | | | | 195 | 162 | 6.08 | 4.69 | 0 | 0.324227 | 40.5318 | 6.32164 |
| | | | | | | 227 | 162 | 6 | 4.64 | 0 | 2.2403 | 33.2907 | 19.4155 |
| | | | | | | 287 | 162 | 5.86 | 4.57 | 0 | -7.79272 | 62.4773 | -3.88864 |
| | | | | | | 544 | 222 | 5.43 | 4.34 | 0 | -38.7284 | 36.1634 | 13.7089 |
| | | | | | | 548 | 222 | 5.43 | 4.34 | 0 | -24.1952 | 65.0344 | 7.08297 |
| | | | | | | 626 | 241 | 5.32 | 4.28 | 0 | 6.41165 | 43.4986 | -5.22219 |
| | | | | | | 683 | 241 | 5.24 | 4.24 | 0 | -39.1996 | 39.9125 | 25.3346 |
| | | | | | | 687 | 241 | 5.23 | 4.23 | 0 | -7.69827 | 69.9642 | 6.11086 |
| | | | | | | 792 | 0.28 | 5.08 | 4.14 | 0 | 6.37112 | 54.0813 | 4.45002 |
| | | | | | | 795 | 0.28 | 5.07 | 4.14 | 0 | -51.5669 | 36.205 | 17.5372 |
| | | | | | | 807 | 283 | 5.06 | 4.13 | 0 | -1.04955 | 45.2159 | -12.1884 |
| | | | | | | 0.92 | 332 | 4.84 | 4 | 0 | -9.83387 | 36.4906 | -11.6584 |
| | | | | | | 963 | 367 | 4.7 | 3.92 | 0 | -12.5093 | 43.7582 | 16.1988 |
| | | | | | | 997 | 478 | 4.39 | 3.73 | 0 | -29.7646 | 55.5373 | 17.0053 |
| | | | | | | 998 | 478 | 4.38 | 3.72 | 0 | -16.722 | 69.3322 | 4.01831 |
| | | | | | | 999 | 494 | 4.32 | 3.68 | 0 | -1.84334 | 57.6522 | -12.297 |
| | | | | | | 999 | 0.5 | 4.27 | 3.65 | 0 | 1.83138 | 65.1654 | -0.317411 |
| | | | | | | 1 | 563 | 4.14 | 3.56 | 0 | -48.7517 | 30.3686 | 30.1743 |
| | | | | | | 1 | 579 | 4.09 | 3.53 | 0 | -15.8093 | 68.0657 | 14.7073 |
| | | 0 | 0 | 8993 | 0 | 104 | 0.16 | 6.43 | 4.86 | 0 | 2.61887 | -62.414 | 59.6056 |
| | | | | | | 139 | 0.16 | 6.27 | 4.78 | 0 | -4.60568 | -38.3258 | 49.871 |
| | | | | | | 225 | 162 | 6 | 4.65 | 0 | -7.2145 | -63.2177 | 63.3121 |
| | | | | | | 382 | 0.17 | 5.68 | 4.48 | 0 | -0.725018 | -48.6208 | 55.6294 |
| | | | | | | 521 | 218 | 5.47 | 4.36 | 0 | 6.48053 | -43.3807 | 56.1645 |
| | | | | | | 628 | 241 | 5.32 | 4.28 | 0 | -12.4937 | -36.0966 | 56.0467 |
| | | | | | | 749 | 268 | 5.15 | 4.18 | 0 | 4.80531 | -21.1556 | 27.0628 |
| | | | | | | 803 | 283 | 5.06 | 4.13 | 0 | -0.441007 | -30.1287 | 41.0362 |
| | | | | | | 881 | 316 | 4.92 | 4.05 | 0 | -12.8413 | -38.8129 | 41.1807 |
| | | | | | | 945 | 359 | 4.76 | 3.96 | 0 | -16.872 | -81.2841 | 56.992 |
| | | | | | | 989 | 426 | 4.53 | 3.81 | 0 | 13.2476 | -58.687 | 52.893 |
| | | | | | | 997 | 478 | 4.39 | 3.73 | 0 | -6.11705 | -45.1994 | 62.0062 |
| | | | | | | 999 | 499 | 4.29 | 3.66 | 0 | -22.6839 | -73.0419 | 60.5568 |
| | | | | | | 999 | 504 | 4.26 | 3.64 | 0 | 1.24396 | -45.8592 | 66.2273 |
| | | | | | | 1 | 563 | 4.12 | 3.55 | 0 | -1.53922 | -75.518 | 51.984 |
| | | | | | | 1 | 588 | 4.04 | 3.5 | 0 | 19.4811 | -68.2862 | 62.4294 |
| | | | | | | 1 | 653 | 3.92 | 3.41 | 0 | 10.1097 | -36.5604 | 35.711 |
| | | 0 | 0 | 2940 | 0 | 155 | 0.16 | 6.21 | 4.75 | 0 | 43.2802 | -3.18247 | 49.1942 |
| | | | | | | 484 | 214 | 5.52 | 4.39 | 0 | 28.6534 | 9.87077 | 62.7497 |
| | | | | | | 902 | 316 | 4.88 | 4.03 | 0 | 22.6104 | -4.67004 | 55.3393 |
| | | | | | | 996 | 464 | 4.44 | 3.75 | 0 | 35.6997 | -1.53687 | 61.7461 |
| | | | | | | 1 | 752 | 3.76 | 3.3 | 0 | 31.7461 | 18.3558 | 56.4116 |
| | | | | | | 1 | 784 | 3.72 | 3.27 | 1 | 24.3311 | 6.16685 | 53.2299 |

| | | | | | | | | | | | | | |
|---|---|---|---|---|---|---|---|---|---|---|---|---|---|
| | | 49 | 12 | 574 | 2 | 221 | 162 | 6.01 | 4.65 | 0 | -58.955 | -30.0116 | 33.1043 |
| | | | | | | 999 | 494 | 4.32 | 3.68 | 0 | -51.5285 | -24.9602 | 36.5377 |
| | | 849 | 288 | 164 | 62 | 222 | 162 | 6.01 | 4.65 | 0 | -15.3893 | -23.4764 | 65.6833 |
| | | 14 | 5 | 749 | 0 | 299 | 162 | 5.83 | 4.56 | 0 | 52.2961 | -43.1548 | 23.4592 |
| | | | | | | 1 | 588 | 4.07 | 3.51 | 0 | 65.0853 | -51.023 | 21.7303 |
| | | | | | | 1 | 588 | 4.06 | 3.51 | 0 | 57.886 | -33.8874 | 32.2228 |
| | | | | | | 1 | 718 | 3.82 | 3.34 | 0 | 59.6279 | -45.2347 | 29.6676 |
| | | 5 | 2 | 914 | 0 | 319 | 162 | 5.79 | 4.54 | 0 | -44.0675 | -53.6369 | -28.7577 |
| | | | | | | 1 | 588 | 4.04 | 3.49 | 0 | -51.9973 | -47.5848 | -39.9107 |
| | | 187 | 45 | 394 | 7 | 321 | 162 | 5.79 | 4.54 | 0 | 33.6005 | -46.7312 | 62.3466 |
| | | 37 | 11 | 612 | 1 | 394 | 172 | 5.66 | 4.47 | 0 | 27.1691 | 70.6161 | 10.8621 |
| | | | | | | 1 | 563 | 4.14 | 3.56 | 0 | 28.3741 | 56.4012 | 10.0675 |
| | | 969 | 414 | 113 | 114 | 457 | 206 | 5.56 | 4.41 | 0 | -13.6382 | -16.9696 | -14.4687 |
| | | 5 | 2 | 915 | 0 | 518 | 218 | 5.47 | 4.36 | 0 | 48.0099 | -12.8343 | -2.77816 |
| | | | | | | 0.87 | 316 | 4.95 | 4.07 | 0 | 68.4584 | -9.4991 | 2.40174 |
| | | | | | | 992 | 0.44 | 4.5 | 3.79 | 0 | 57.3735 | -5.92183 | -3.88319 |
| | | | | | | 999 | 499 | 4.29 | 3.66 | 0 | 55.0869 | -12.5542 | 4.49805 |
| | | | | | | 1 | 642 | 3.94 | 3.42 | 0 | 64.1483 | -16.3096 | -0.181113 |
| | | 1 | 0 | 1249 | 0 | 605 | 241 | 5.35 | 4.3 | 0 | 35.5005 | 35.2868 | 39.8821 |
| | | | | | | 1 | 588 | 4.04 | 3.5 | 0 | 31.4944 | 32.2716 | 29.4483 |
| | | 198 | 46 | 387 | 7 | 0.62 | 241 | 5.33 | 4.29 | 0 | 31.4576 | 19.5731 | -7.81177 |
| | | | | | | 1 | 563 | 4.17 | 3.58 | 0 | 45.9206 | 17.8522 | -8.46594 |
| | | 1 | 0 | 1287 | 0 | 629 | 241 | 5.31 | 4.28 | 0 | -24.5801 | 31.5844 | -13.8161 |
| | | | | | | 0.64 | 241 | 5.3 | 4.27 | 0 | -34.3213 | 36.2495 | -4.44418 |
| | | | | | | 954 | 359 | 4.73 | 3.94 | 0 | -38.0034 | 48.2243 | -9.31583 |
| | | | | | | 996 | 464 | 4.44 | 3.75 | 0 | -33.9846 | 36.7072 | -14.4308 |
| | | | | | | 1 | 563 | 4.16 | 3.58 | 0 | -21.5987 | 42.4638 | -13.087 |
| | | 0 | 0 | 1496 | 0 | 629 | 241 | 5.31 | 4.28 | 0 | -49.6666 | 3.93889 | 34.3283 |
| | | | | | | 674 | 241 | 5.25 | 4.24 | 0 | -52.6961 | -1.75561 | 49.9498 |
| | | | | | | 959 | 361 | 4.71 | 3.93 | 0 | -54.7304 | 1.85868 | 20.0625 |
| | | | | | | 976 | 393 | 4.64 | 3.88 | 0 | -35.4286 | 1.50681 | 24.7269 |
| | | | | | | 1 | 563 | 4.12 | 3.55 | 0 | -45.9113 | -14.6214 | 48.5797 |
| | | | | | | 1 | 588 | 4.05 | 3.5 | 0 | -34.5856 | -18.6956 | 38.5757 |
| | | | | | | 1 | 644 | 3.93 | 3.42 | 0 | -42.6344 | -7.85362 | 55.172 |
| | | 0 | 0 | 3871 | 0 | 666 | 241 | 5.26 | 4.25 | 0 | -45.982 | -49.8208 | 49.9826 |
| | | | | | | 752 | 268 | 5.14 | 4.18 | 0 | -34.6058 | -43.2818 | 58.3317 |
| | | | | | | 761 | 268 | 5.13 | 4.17 | 0 | -26.1956 | -46.9571 | 45.5055 |
| | | | | | | 883 | 316 | 4.92 | 4.05 | 0 | -20.4123 | -35.9377 | 74.8543 |
| | | | | | | 973 | 392 | 4.65 | 3.89 | 0 | -37.4579 | -43.4383 | 40.0465 |
| | | | | | | 979 | 396 | 4.61 | 3.87 | 0 | -31.1608 | -39.9764 | 69.0662 |
| | | | | | | 995 | 458 | 4.45 | 3.76 | 0 | -31.3399 | -37.163 | 50.7555 |
| | | | | | | 1 | 563 | 4.13 | 3.56 | 0 | -42.573 | -33.5034 | 37.8379 |
| | | | | | | 1 | 563 | 4.12 | 3.55 | 0 | -40.7502 | -46.5647 | 63.9897 |
| | | 2 | 1 | 1078 | 0 | 686 | 241 | 5.23 | 4.23 | 0 | -27.7247 | 1.53533 | 66.0439 |
| | | | | | | 0.99 | 428 | 4.52 | 3.81 | 0 | -30.3735 | -4.05811 | 54.3694 |
| | | | | | | 1 | 833 | 3.66 | 3.23 | 1 | -29.0702 | 14.875 | 59.918 |
| | | 242 | 56 | 360 | 9 | 749 | 268 | 5.14 | 4.18 | 0 | -16.2817 | -51.2971 | 71.3499 |
| | | | | | | 998 | 483 | 4.35 | 3.7 | 0 | -7.82916 | -48.2892 | 75.9557 |
| | | 31 | 9 | 638 | 1 | 751 | 268 | 5.14 | 4.18 | 0 | 25.3852 | -34.2415 | 11.4323 |
| | | | | | | 782 | 277 | 5.1 | 4.15 | 0 | 15.9466 | -50.5315 | -2.44857 |
| | | | | | | 997 | 478 | 4.39 | 3.72 | 0 | 26.27 | -38.4211 | 2.987 |

| | | | | | | | | | | | | | |
|---|---|---|---|---|---|---|---|---|---|---|---|---|---|
| | | 2 | 1 | 1068 | 0 | 776 | 275 | 5.1 | 4.16 | 0 | -27.4321 | 23.011 | 51.2865 |
| | | | | | | 998 | 483 | 4.36 | 3.71 | 0 | -24.7453 | 43.8839 | 44.5841 |
| | | | | | | 1 | 563 | 4.16 | 3.57 | 0 | -20.7892 | 33.5594 | 49.2793 |
| | | | | | | 1 | 0.68 | 3.87 | 3.38 | 0 | -23.8679 | 32.1157 | 36.0681 |
| | | 337 | 0.08 | 316 | 13 | 825 | 297 | 5.03 | 4.11 | 0 | -11.3311 | -27.1883 | 80.3457 |
| | | 19 | 6 | 705 | 1 | 836 | 301 | 5.01 | 4.1 | 0 | 22.2523 | -35.9624 | 73.6451 |
| | | | | | | 853 | 0.31 | 4.98 | 4.09 | 0 | 18.3981 | -30.0486 | 61.6943 |
| | | | | | | 999 | 497 | 4.31 | 3.67 | 0 | 15.4319 | -25.3749 | 70.6371 |
| | | | | | | 1 | 816 | 3.68 | 3.24 | 1 | 27.6338 | -22.1845 | 72.8738 |
| | | 581 | 144 | 237 | 29 | 865 | 316 | 4.95 | 4.07 | 0 | -21.0559 | -32.8142 | 8.08291 |
| | | 49 | 12 | 574 | 2 | 883 | 316 | 4.92 | 4.05 | 0 | -45.1937 | 3.95415 | -8.81873 |
| | | | | | | 979 | 396 | 4.62 | 3.87 | 0 | -47.6593 | 5.8793 | 2.95046 |
| | | | | | | 1 | 722 | 3.81 | 3.34 | 0 | -37.7456 | 7.10282 | 2.04132 |
| | | 0 | 0 | 1434 | 0 | 893 | 316 | 4.9 | 4.04 | 0 | -4.10857 | 22.1431 | 24.0257 |
| | | | | | | 997 | 468 | 4.41 | 3.74 | 0 | 3.06005 | 14.2579 | 28.588 |
| | | | | | | 999 | 504 | 4.26 | 3.64 | 0 | -0.854521 | 12.8765 | 43.7427 |
| | | 724 | 207 | 199 | 42 | 898 | 316 | 4.89 | 4.03 | 0 | 16.0139 | 11.0616 | 18.0806 |
| | | 938 | 373 | 131 | 91 | 0.9 | 316 | 4.88 | 4.03 | 0 | -22.149 | -45.4179 | 5.70333 |
| | | 1 | 1 | 1115 | 0 | 926 | 338 | 4.82 | 3.99 | 0 | -20.8499 | -77.1173 | -12.5866 |
| | | | | | | 998 | 483 | 4.36 | 3.7 | 0 | -33.9216 | -69.0365 | -19.1693 |
| | | | | | | 999 | 497 | 4.31 | 3.67 | 0 | -12.4263 | -77.3474 | -9.15254 |
| | | | | | | 1 | 774 | 3.73 | 3.28 | 1 | -5.83907 | -81.1872 | -3.99767 |
| | | 864 | 289 | 159 | 65 | 974 | 392 | 4.64 | 3.88 | 0 | 41.3514 | -8.26357 | -26.2849 |
| | | 535 | 135 | 250 | 25 | 975 | 393 | 4.64 | 3.88 | 0 | 44.5869 | -70.6761 | -42.9032 |
| | | | | | | 1 | 567 | 4.11 | 3.54 | 0 | 51.6647 | -69.4946 | -31.5354 |
| | | 567 | 143 | 241 | 27 | 979 | 396 | 4.61 | 3.86 | 0 | 44.5181 | -53.7674 | 16.5943 |
| | | 819 | 267 | 173 | 56 | 979 | 396 | 4.61 | 3.86 | 0 | 17.8851 | -10.218 | 13.138 |
| | | 2 | 1 | 1066 | 0 | 982 | 398 | 4.6 | 3.86 | 0 | -6.31656 | 2.32052 | 62.2421 |
| | | | | | | 982 | 398 | 4.59 | 3.85 | 0 | 11.2729 | -5.9599 | 42.9877 |
| | | | | | | 993 | 444 | 4.48 | 3.78 | 0 | 1.55363 | -2.94452 | 69.3505 |
| | | | | | | 1 | 563 | 4.12 | 3.55 | 0 | -7.6604 | -5.88621 | 44.0101 |
| | | | | | | 1 | 603 | 4.01 | 3.47 | 0 | 2.46953 | -4.25512 | 38.5672 |
| | | 44 | 12 | 588 | 1 | 988 | 424 | 4.55 | 3.82 | 0 | 59.7054 | -27.9497 | 18.1151 |
| | | | | | | 1 | 0.67 | 3.89 | 3.39 | 0 | 41.7737 | -35.0523 | 13.3781 |
| | | | | | | 1 | 692 | 3.86 | 3.37 | 0 | 61.5499 | -18.7324 | 20.0971 |
| | | 494 | 124 | 262 | 22 | 0.99 | 428 | 4.52 | 3.81 | 0 | 30.8177 | 5.35179 | 15.808 |
| | | | | | | 1 | 0.56 | 4.17 | 3.58 | 0 | 35.3275 | 8.28082 | 6.24441 |
| | | 968 | 414 | 114 | 113 | 993 | 444 | 4.49 | 3.78 | 0 | 1.49193 | -3.35455 | 4.40769 |
| | | 115 | 28 | 458 | 4 | 994 | 452 | 4.46 | 3.77 | 0 | -7.43044 | -51.4138 | 25.9169 |
| | | | | | | 998 | 483 | 4.35 | 3.7 | 0 | -12.377 | -52.8393 | 9.15009 |
| | | | | | | 1 | 718 | 3.81 | 3.34 | 0 | -2.50485 | -53.6858 | 11.4579 |
| | | | | | | 1 | 735 | 3.79 | 3.32 | 0 | -10.6777 | -57.3757 | 18.8467 |
| | | 905 | 331 | 145 | 77 | 996 | 467 | 4.42 | 3.74 | 0 | 12.2159 | -18.8713 | 14.9782 |
| | | 0.07 | 17 | 525 | 2 | 998 | 0.48 | 4.38 | 3.72 | 0 | -70.1383 | -23.7401 | 12.645 |
| | | | | | | 999 | 499 | 4.29 | 3.66 | 0 | -67.453 | -16.5285 | 8.56853 |
| | | | | | | 1 | 659 | 3.91 | 3.4 | 0 | -69.4205 | -17.2744 | 21.8499 |
| | | | | | | 1 | 752 | 3.77 | 3.3 | 0 | -60.536 | -28.2042 | 13.8643 |
| | | 971 | 414 | 112 | 116 | 998 | 483 | 4.37 | 3.71 | 0 | 18.7069 | -2.89499 | 73.4374 |
| | | 963 | 414 | 117 | 108 | 999 | 499 | 4.29 | 3.66 | 0 | -4.80432 | -53.3869 | -15.8504 |
| | | 378 | 89 | 300 | 16 | 999 | 499 | 4.29 | 3.66 | 0 | -32.99 | -45.1727 | -35.3193 |
| | | | | | | 1 | 972 | 3.51 | 3.12 | 1 | -25.7336 | -51.7676 | -29.2105 |

| | | | | | | | | | | | | | |
|---|---|---|---|---|---|---|---|---|---|---|---|---|---|
| | | 974 | 422 | 109 | 0.12 | 999 | 0.5 | 4.27 | 3.65 | 0 | -30.942 | -11.9413 | -17.5625 |
| | | 932 | 369 | 134 | 88 | 1 | 521 | 4.23 | 3.62 | 0 | 46.8253 | -52.5648 | -27.8798 |
| | | 952 | 397 | 124 | 99 | 1 | 563 | 4.13 | 3.55 | 0 | -1.18201 | -19.3338 | 7.31102 |
| | | 864 | 289 | 159 | 65 | 1 | 563 | 4.12 | 3.55 | 0 | -46.0846 | -11.2361 | -0.094845[illegible] |
| | | 968 | 414 | 114 | 113 | 1 | 625 | 3.96 | 3.44 | 0 | -30.2348 | 13.2987 | 8.77845 |

PDnoLID-Reward_OFF

| p | c | p(FWE-corr) | p(FDR-corr) | equivk | p(unc) | p(FWE-corr).1 | p(FDR-corr).1 | T | equivZ | p(unc).1 | MNI_x | MNI_y | MNI_z |
|---|---|---|---|---|---|---|---|---|---|---|---|---|---|
| 0 | 23 | 96 | 55 | 418 | 3 | 383 | 592 | 6.04 | 4.55 | 0 | -18.8085 | 7.64256 | -12.0834 |
| | | 117 | 55 | 396 | 3 | 454 | 592 | 5.91 | 4.49 | 0 | 28.6849 | 21.9265 | 6.00613 |
| | | 0.95 | 489 | 116 | 81 | 561 | 592 | 5.73 | 4.4 | 0 | 53.0167 | -15.333 | -16.205 |
| | | 946 | 489 | 118 | 79 | 655 | 592 | 5.59 | 4.33 | 0 | -11.3199 | -74.2207 | -11.4862 |
| | | 142 | 55 | 374 | 4 | 686 | 592 | 5.54 | 4.3 | 0 | -17.5481 | -98.0773 | 2.18999 |
| | | | | | | 1 | 994 | 3.57 | 3.12 | 1 | -19.5073 | -94.86 | -10.5715 |
| | | 137 | 55 | 378 | 4 | 747 | 592 | 5.45 | 4.25 | 0 | 16.0969 | 10.0789 | -12.77 |
| | | | | | | 1 | 918 | 3.89 | 3.34 | 0 | 22.5709 | 14.5263 | -5.45101 |
| | | 127 | 55 | 387 | 4 | 801 | 592 | 5.36 | 4.21 | 0 | 28.1993 | -94.8323 | 12.6286 |
| | | 958 | 489 | 112 | 86 | 867 | 677 | 5.23 | 4.14 | 0 | -8.89408 | -11.11 | 74.7775 |
| | | 964 | 489 | 109 | 0.09 | 918 | 737 | 5.11 | 4.08 | 0 | 25.344 | -87.3772 | 22.8282 |
| | | 0.09 | 55 | 425 | 3 | 923 | 737 | 5.1 | 4.07 | 0 | -57.8248 | -49.5017 | 42.8091 |
| | | 7 | 21 | 738 | 0 | 948 | 789 | 5.02 | 4.02 | 0 | 14.5427 | -93.5082 | -7.21896 |
| | | | | | | 1 | 876 | 4.31 | 3.61 | 0 | 10.5729 | -84.3928 | -2.39842 |
| | | 0.95 | 489 | 116 | 81 | 972 | 789 | 4.91 | 3.96 | 0 | -28.0897 | -3.11006 | -11.6004 |
| | | | | | | 1 | 789 | 4.46 | 3.7 | 0 | -23.3234 | -10.1253 | -12.1221 |
| | | 0.92 | 489 | 128 | 68 | 982 | 789 | 4.85 | 3.93 | 0 | 25.5674 | 23.5091 | 40.4239 |
| | | 23 | 37 | 582 | 1 | 982 | 789 | 4.85 | 3.93 | 0 | 6.47041 | -70.1908 | 58.8061 |
| | | | | | | 1 | 916 | 4.23 | 3.56 | 0 | 10.5037 | -66.08 | 50.3108 |
| | | 75 | 55 | 445 | 2 | 991 | 789 | 4.75 | 3.87 | 0 | -34.9702 | 27.2345 | 35.851 |
| | | 0.64 | 326 | 198 | 28 | 999 | 789 | 4.53 | 3.74 | 0 | 0.00521841 | -94.4967 | -4.49468 |
| | | 914 | 489 | 130 | 67 | 999 | 789 | 4.52 | 3.74 | 0 | 25.0152 | -24.7649 | 12.5889 |
| | | 954 | 489 | 114 | 83 | 1 | 789 | 4.47 | 3.7 | 0 | 23.9988 | -69.8472 | -15.1933 |
| | | 764 | 0.42 | 170 | 39 | 1 | 789 | 4.41 | 3.67 | 0 | -41.5578 | 48.036 | 25.5149 |
| | | | | | | 1 | 918 | 3.85 | 3.31 | 0 | -38.5851 | 40.1499 | 14.9813 |
| | | 966 | 489 | 108 | 91 | 1 | 789 | 4.4 | 3.66 | 0 | -8.98637 | -66.6351 | 53.284 |
| | | 883 | 489 | 140 | 58 | 1 | 915 | 4.25 | 3.57 | 0 | -7.52968 | 13.1017 | -3.46107 |
| | | 973 | 504 | 103 | 98 | 1 | 915 | 4.24 | 3.56 | 0 | 27.3507 | -63.1065 | 58.7952 |
| | | 943 | 489 | 119 | 78 | 1 | 918 | 4.13 | 3.49 | 0 | 61.0786 | -45.8201 | 21.1999 |

LID-Reward_OFF

| p | c | p(FWE-corr) | p(FDR-corr) | equivk | p(unc) | p(FWE-corr).1 | p(FDR-corr).1 | T | equivZ | p(unc).1 | MNI_x | MNI_y | MNI_z |
|---|---|---|---|---|---|---|---|---|---|---|---|---|---|
| **0** | 27 | 0.01 | 0.01 | 769 | 0 | 88 | 484 | 6.22 | 4.89 | 0 | -20.056 | 8.73581 | -13.2316 |
| | | | | | | 999 | 728 | 4.21 | 3.67 | 0 | -23.3959 | 17.4875 | -1.93123 |
| | | 32 | 21 | 610 | 1 | 326 | 498 | 5.56 | 4.53 | 0 | -27.8957 | 39.5399 | -13.3055 |
| | | | | | | 758 | 565 | 4.97 | 4.17 | 0 | -18.2817 | 48.1213 | -11.2478 |
| | | 728 | 346 | 192 | 41 | 354 | 498 | 5.51 | 4.5 | 0 | -24.1976 | -20.274 | -18.1965 |
| | | 775 | 371 | 180 | 47 | 412 | 498 | 5.43 | 4.45 | 0 | -40.9604 | -83.8302 | -2.83397 |
| | | 5 | 9 | 886 | 0 | 419 | 498 | 5.41 | 4.44 | 0 | 35.2971 | -93.0625 | 17.2383 |
| | | | | | | 977 | 0.68 | 4.51 | 3.88 | 0 | 23.5086 | -101.816 | 13.9141 |
| | | | | | | 985 | 0.68 | 4.46 | 3.84 | 0 | 8.81975 | -93.8438 | 20.3256 |
| | | | | | | 1 | 781 | 4.01 | 3.53 | 0 | 19.9925 | -92.842 | 23.5635 |
| | | 3 | 9 | 936 | 0 | 446 | 498 | 5.38 | 4.42 | 0 | -3.42371 | 13.0142 | 36.2951 |
| | | | | | | 1 | 919 | 3.82 | 3.39 | 0 | -2.27232 | 0.146246 | 35.7692 |
| | | | | | | 1 | 928 | 3.76 | 3.36 | 0 | 5.72995 | 10.7176 | 40.0035 |
| | | 46 | 27 | 560 | 1 | 522 | 498 | 5.27 | 4.36 | 0 | 5.22163 | 54.7684 | -4.10422 |
| | | 863 | 416 | 155 | 63 | 531 | 498 | 5.26 | 4.35 | 0 | 32.5116 | -39.737 | -8.10833 |
| | | | | | | 1 | 781 | 4.02 | 3.54 | 0 | 25.2057 | -44.412 | -5.81666 |
| | | 375 | 208 | 291 | 15 | 572 | 498 | 5.21 | 4.32 | 0 | 29.5074 | 23.5904 | 43.4962 |
| | | | | | | 1 | 956 | 3.63 | 3.26 | 1 | 32.1177 | 16.6573 | 49.3339 |
| | | 0.42 | 217 | 276 | 17 | 761 | 565 | 4.97 | 4.17 | 0 | 27.4047 | 42.5668 | 35.7184 |
| | | 962 | 516 | 115 | 104 | 773 | 565 | 4.95 | 4.16 | 0 | 26.9963 | -22.3901 | 19.4981 |
| | | 8 | 0.01 | 803 | 0 | 776 | 565 | 4.95 | 4.16 | 0 | 9.13197 | 14.8388 | -6.62257 |
| | | | | | | 989 | 0.68 | 4.42 | 3.82 | 0 | 18.3672 | 7.88826 | -7.85294 |
| | | | | | | 999 | 728 | 4.18 | 3.65 | 0 | 26.1907 | 12.6166 | -3.28695 |
| | | 17 | 14 | 692 | 1 | 0.88 | 0.57 | 4.78 | 4.05 | 0 | -14.1169 | -58.6895 | 14.0467 |
| | | | | | | 1 | 757 | 4.14 | 3.63 | 0 | -21.983 | -63.583 | 10.238 |
| | | 913 | 0.44 | 138 | 77 | 881 | 0.57 | 4.78 | 4.05 | 0 | -35.8834 | -7.65577 | -4.19615 |
| | | 897 | 431 | 144 | 72 | 0.9 | 0.57 | 4.74 | 4.03 | 0 | -66.8172 | -27.9698 | 39.3672 |
| | | 0.52 | 244 | 246 | 23 | 907 | 0.57 | 4.73 | 4.02 | 0 | 14.2368 | -30.4142 | 41.4807 |
| | | 876 | 416 | 151 | 66 | 913 | 0.57 | 4.72 | 4.01 | 0 | 22.345 | -24.5761 | 70.1649 |
| | | 634 | 286 | 216 | 32 | 965 | 0.68 | 4.57 | 3.91 | 0 | -24.5951 | 2.56253 | 66.6281 |
| | | | | | | 1 | 781 | 4 | 3.53 | 0 | -22.4381 | 11.2524 | 64.0731 |
| | | 0.98 | 559 | 102 | 123 | 967 | 0.68 | 4.56 | 3.91 | 0 | 28.7701 | -2.13258 | 13.221 |
| | | 854 | 416 | 158 | 61 | 0.98 | 0.68 | 4.49 | 3.87 | 0 | -39.768 | 3.88449 | 8.00482 |
| | | 622 | 286 | 219 | 31 | 988 | 0.68 | 4.44 | 3.83 | 0 | 8.87242 | 36.6667 | -6.99081 |
| | | | | | | 1 | 781 | 4.06 | 3.57 | 0 | 3.7305 | 35.9524 | -15.5973 |
| | | 137 | 74 | 419 | 5 | 999 | 728 | 4.24 | 3.69 | 0 | -5.79073 | -32.0246 | 53.0043 |
| | | | | | | 999 | 728 | 4.22 | 3.68 | 0 | -13.3729 | -29.7998 | 40.2679 |
| | | 965 | 516 | 113 | 106 | 999 | 728 | 4.22 | 3.68 | 0 | -21.0684 | -55.7705 | -9.1074 |
| | | 921 | 0.44 | 135 | 0.08 | 999 | 728 | 4.18 | 3.65 | 0 | -25.4307 | -62.2876 | -13.3594 |
| | | 833 | 416 | 164 | 57 | 1 | 781 | 4.01 | 3.53 | 0 | 22.2401 | 2.20301 | 62.9147 |
| | | 952 | 503 | 121 | 96 | 1 | 855 | 3.9 | 3.45 | 0 | -10.1313 | -94.085 | -2.15172 |
| | | | | | | 1 | 929 | 3.7 | 3.31 | 0 | -4.27184 | -97.6676 | 7.03833 |
| | | 502 | 244 | 251 | 22 | 1 | 919 | 3.8 | 3.38 | 0 | -36.6377 | 32.709 | 38.7225 |
| | | | | | | 1 | 931 | 3.67 | 3.29 | 1 | -29.6533 | 34.9678 | 29.07 |

HC-Motivational_Context_OFF

| p | c | p(FWE-corr) | p(FDR-corr) | equivk | p(unc) | p(FWE-corr).1 | p(FDR-corr).1 | T | equivZ | p(unc).1 | MNI_x | MNI_y | MNI_z |
|---|---|---|---|---|---|---|---|---|---|---|---|---|---|
| **0** | 26 | 0 | 0 | 11572 | 0 | 1 | 5 | 9.08 | 5.93 | 0 | -3.18847 | -77.5336 | -0.447524 |
| | | | | | | 25 | 65 | 7.26 | 5.23 | 0 | -6.92468 | -104.026 | 0.761105 |
| | | | | | | 64 | 65 | 6.75 | 5 | 0 | -20.6499 | -71.896 | -8.61178 |
| | | | | | | 171 | 115 | 6.22 | 4.75 | 0 | -29.6189 | -75.1184 | -16.3869 |
| | | | | | | 247 | 157 | 6.01 | 4.65 | 0 | 27.4147 | -73.341 | -12.2022 |
| | | | | | | 716 | 462 | 5.25 | 4.24 | 0 | -21.0806 | -103.975 | -4.93576 |
| | | | | | | 922 | 486 | 4.89 | 4.03 | 0 | 15.885 | -71.8898 | -12.5235 |
| | | | | | | 929 | 486 | 4.87 | 4.02 | 0 | -13.1701 | -84.8862 | -13.3899 |
| | | | | | | 976 | 593 | 4.7 | 3.92 | 0 | -20.9158 | -80.5859 | -16.5676 |
| | | | | | | 1 | 734 | 4.23 | 3.62 | 0 | 34.112 | -60.6445 | -17.8089 |
| | | | | | | 1 | 734 | 4.19 | 3.59 | 0 | 34.6754 | -51.009 | -10.2055 |
| | | | | | | 1 | 888 | 3.8 | 3.33 | 0 | -28.7669 | -86.3837 | -18.0064 |
| | | | | | | 1 | 913 | 3.76 | 3.3 | 0 | 0.571934 | -66.4604 | 12.2422 |
| | | | | | | 1 | 913 | 3.72 | 3.27 | 1 | -2.44963 | -77.4485 | -15.2532 |
| | | 0 | 0 | 5001 | 0 | 48 | 65 | 6.91 | 5.08 | 0 | 8.302 | -91.9646 | 10.0778 |
| | | | | | | 61 | 65 | 6.78 | 5.02 | 0 | 13.2611 | -97.8972 | 18.2551 |
| | | | | | | 277 | 163 | 5.94 | 4.61 | 0 | 9.71486 | -98.5768 | -2.15757 |
| | | | | | | 363 | 207 | 5.77 | 4.53 | 0 | 12.9986 | -100.834 | 8.27081 |
| | | | | | | 791 | 0.47 | 5.14 | 4.18 | 0 | 21.5428 | -100.582 | 1.39264 |
| | | | | | | 826 | 0.47 | 5.08 | 4.15 | 0 | 29.4174 | -99.1322 | -5.6449 |
| | | | | | | 999 | 684 | 4.37 | 3.71 | 0 | 28.6747 | -98.7473 | 9.30872 |
| | | | | | | 1 | 815 | 4.05 | 3.5 | 0 | 20.4435 | -96.0789 | -11.8617 |
| | | 14 | 12 | 679 | 0 | 0.16 | 115 | 6.25 | 4.77 | 0 | 10.0177 | 21.4569 | 49.5521 |
| | | | | | | 564 | 344 | 5.46 | 4.36 | 0 | 19.4029 | 24.6086 | 37.9976 |
| | | | | | | 1 | 0.74 | 4.13 | 3.55 | 0 | 15.4976 | 29.532 | 30.6769 |
| | | 369 | 124 | 280 | 13 | 582 | 344 | 5.44 | 4.35 | 0 | -43.9238 | -55.7247 | -42.738 |
| | | 19 | 13 | 638 | 1 | 776 | 0.47 | 5.17 | 4.19 | 0 | -15.3249 | -42.3151 | -30.4948 |
| | | | | | | 923 | 486 | 4.89 | 4.03 | 0 | -3.90939 | -45.6574 | -28.1941 |
| | | | | | | 1 | 0.83 | 3.98 | 3.45 | 0 | -16.933 | -50.8502 | -33.9235 |
| | | 58 | 32 | 500 | 2 | 802 | 0.47 | 5.12 | 4.17 | 0 | -21.6146 | -48.715 | -5.76395 |
| | | | | | | 1 | 734 | 4.23 | 3.62 | 0 | -34.8623 | -53.5426 | -9.9813 |
| | | | | | | 1 | 817 | 4.03 | 3.49 | 0 | -23.0158 | -38.6191 | -8.38777 |
| | | 939 | 412 | 124 | 82 | 0.88 | 486 | 4.99 | 4.09 | 0 | -26.7683 | -62.4672 | 45.063 |
| | | 0.5 | 158 | 241 | 0.02 | 885 | 486 | 4.98 | 4.08 | 0 | -47.4791 | 15.4063 | 33.1141 |
| | | 965 | 471 | 111 | 98 | 901 | 486 | 4.94 | 4.06 | 0 | 46.5378 | -43.4191 | -40.8636 |
| | | 234 | 102 | 335 | 8 | 918 | 486 | 4.9 | 4.04 | 0 | 17.8167 | -36.3223 | -18.9799 |
| | | 137 | 68 | 398 | 4 | 0.92 | 486 | 4.9 | 4.04 | 0 | 41.1523 | -21.9103 | -14.8803 |
| | | | | | | 1 | 894 | 3.79 | 3.32 | 0 | 46.5775 | -19.0289 | -25.6797 |
| | | 329 | 119 | 294 | 12 | 962 | 563 | 4.76 | 3.96 | 0 | -17.8024 | -34.8946 | 12.7877 |
| | | 853 | 311 | 153 | 56 | 977 | 593 | 4.69 | 3.91 | 0 | -7.38186 | -49.1302 | -38.5163 |
| | | | | | | 1 | 931 | 3.62 | 3.2 | 1 | -8.89181 | -52.9962 | -29.3559 |
| | | 846 | 311 | 155 | 55 | 988 | 0.62 | 4.61 | 3.86 | 0 | -36.2206 | -49.8873 | 57.1387 |
| | | 745 | 0.26 | 181 | 0.04 | 989 | 0.62 | 4.59 | 3.85 | 0 | -14.4829 | -74.4105 | 47.0755 |
| | | 332 | 119 | 293 | 12 | 0.99 | 0.62 | 4.58 | 3.85 | 0 | 52.4775 | -57.9799 | -34.4147 |
| | | | | | | 1 | 734 | 4.23 | 3.62 | 0 | 45.6759 | -69.5535 | -27.2491 |
| | | | | | | 1 | 913 | 3.67 | 3.24 | 1 | 37.5419 | -74.3064 | -25.7492 |
| | | 436 | 143 | 259 | 17 | 992 | 0.62 | 4.55 | 3.83 | 0 | -22.1271 | -79.9844 | -34.5174 |
| | | | | | | 1 | 832 | 3.93 | 3.42 | 0 | -21.9067 | -88.6185 | -32.4821 |
| | | 0.01 | 11 | 730 | 0 | 993 | 0.62 | 4.54 | 3.82 | 0 | 7.15647 | -90.4452 | -25.3398 |

| | | | | | | | | | | | | | |
|---|---|---|---|---|---|---|---|---|---|---|---|---|---|
| | | | | | | 998 | 642 | 4.44 | 3.76 | 0 | 16.3682 | -88.1875 | -26.4555 |
| | | | | | | 1 | 734 | 4.18 | 3.59 | 0 | -4.45605 | -79.5174 | -29.1783 |
| | | | | | | 1 | 797 | 4.07 | 3.52 | 0 | 3.37423 | -81.4785 | -28.2699 |
| | | 0.98 | 487 | 100 | 114 | 995 | 0.62 | 4.5 | 3.8 | 0 | 33.415 | -39.7387 | -6.71147 |
| | | 978 | 487 | 102 | 111 | 998 | 642 | 4.43 | 3.75 | 0 | -7.92486 | -72.8607 | 45.7303 |
| | | 832 | 311 | 159 | 52 | 998 | 659 | 4.41 | 3.74 | 0 | 24.7654 | 43.6916 | 41.8325 |
| | | 248 | 102 | 328 | 8 | 999 | 684 | 4.37 | 3.71 | 0 | 42.9571 | -48.9299 | 56.2943 |
| | | 554 | 163 | 227 | 24 | 1 | 734 | 4.26 | 3.64 | 0 | 4.23663 | 47.689 | 41.7818 |
| | | | | | | 1 | 913 | 3.74 | 3.29 | 1 | 2.69118 | 33.8687 | 28.9357 |
| | | 867 | 312 | 149 | 59 | 1 | 734 | 4.24 | 3.63 | 0 | -29.5362 | -82.0399 | 13.2884 |
| | | 519 | 158 | 236 | 21 | 1 | 0.74 | 4.14 | 3.56 | 0 | 60.836 | -54.491 | -11.3779 |
| | | | | | | 1 | 832 | 3.95 | 3.44 | 0 | 53.8843 | -52.6714 | -18.9039 |
| | | 979 | 487 | 101 | 113 | 1 | 832 | 3.95 | 3.44 | 0 | 36.9229 | -48.5632 | -30.0055 |

PDnoLID-Motivational_Context_OFF

| p | c | p(FWE-corr) | p(FDR-corr) | equivk | p(unc) | p(FWE-corr).1 | p(FDR-corr).1 | T | equivZ | p(unc).1 | MNI_x | MNI_y | MNI_z |
|---|---|---|---|---|---|---|---|---|---|---|---|---|---|
| **272** | 5 | 0 | 0 | 1250 | 0 | 111 | 137 | 6.92 | 4.94 | 0 | 10.0351 | -98.8007 | 14.2387 |
| | | | | | | 1 | 946 | 4.24 | 3.57 | 0 | 10.596 | -90.2804 | 13.8271 |
| | | 0 | 0 | 1461 | 0 | 0.28 | 191 | 6.33 | 4.68 | 0 | -9.12932 | -104.225 | 4.34158 |
| | | | | | | 1 | 946 | 4.34 | 3.62 | 0 | -12.9976 | -103.637 | 20.6709 |
| | | | | | | 1 | 946 | 4.17 | 3.52 | 0 | -26.3839 | -101.037 | 18.5576 |
| | | 0 | 0 | 2078 | 0 | 573 | 0.24 | 5.79 | 4.43 | 0 | -7.0737 | -78.5704 | -1.29513 |
| | | | | | | 678 | 0.24 | 5.63 | 4.35 | 0 | -11.2642 | -76.6552 | -9.22834 |
| | | | | | | 692 | 0.24 | 5.61 | 4.34 | 0 | 13.0594 | -76.024 | -0.488959 |
| | | | | | | 1 | 946 | 4.23 | 3.56 | 0 | -21.248 | -73.6027 | -5.74555 |
| | | | | | | 1 | 946 | 3.99 | 3.4 | 0 | 4.87448 | -79.6924 | 4.45718 |
| | | 742 | 0.2 | 162 | 32 | 0.95 | 499 | 5.08 | 4.06 | 0 | -29.3837 | -49.2041 | 3.74245 |
| | | 667 | 0.2 | 177 | 26 | 1 | 946 | 4.55 | 3.76 | 0 | -2.77981 | -45.8293 | -26.9482 |

LID-Motivational_Context_OFF

| p | c | p(FWE-corr) | p(FDR-corr) | equivk | p(unc) | p(FWE-corr).1 | p(FDR-corr).1 | T | equivZ | p(unc).1 | MNI_x | MNI_y | MNI_z |
|---|---|---|---|---|---|---|---|---|---|---|---|---|---|
| **0.31** | 5 | 0 | 0 | 3040 | 0 | 1 | 1 | 8.29 | 5.84 | 0 | -4.40515 | -78.5793 | 0.754075 |
| | | | | | | 15 | 4 | 7.12 | 5.33 | 0 | -20.8818 | -70.6299 | -2.46866 |
| | | | | | | 0.69 | 0.09 | 5.14 | 4.27 | 0 | 6.63409 | -77.0573 | 2.58512 |
| | | 0 | 0 | 1917 | 0 | 37 | 7 | 6.71 | 5.14 | 0 | 7.5831 | -92.6767 | 15.6364 |
| | | | | | | 566 | 75 | 5.29 | 4.37 | 0 | 9.99052 | -101.001 | 11.9093 |
| | | 969 | 0.28 | 104 | 93 | 555 | 75 | 5.31 | 4.38 | 0 | -19.0633 | -42.6613 | 35.2971 |
| | | 708 | 157 | 179 | 33 | 912 | 164 | 4.8 | 4.06 | 0 | -27.7351 | -51.7708 | -7.43214 |
| | | 789 | 157 | 161 | 42 | 999 | 415 | 4.29 | 3.73 | 0 | -11.5257 | -102.292 | -1.20944 |

**Supplementary Table 2. Whole-brain SPM results for the contrasts of interest within each group.** For the right-go and left-go contrasts, results are reported at a voxel-level threshold of $P_{\mathrm{FWE}} < 0.05$, with a minimum cluster extent of $k = 100$ voxels. For the no-go, reward, and motivational context contrasts, results are reported at an uncorrected voxel-level threshold of $P_{\mathrm{uncorr}} < 0.001$, with a minimum cluster extent of $k = 100$ voxels. Coordinates are reported in MNI space. FWE, family-wise error; MNI, Montreal Neurological Institute; SPM, Statistical Parametric Mapping.
